# Acute stress reorganizes pregenual cingulate and subthalamic coupling to approach-avoidance decisions in remitted depression

**DOI:** 10.64898/2026.10.08.757641

**Authors:** Manuel Kuhn, Nadja R. Ging-Jehli, Emma H. Palermo, Pranavan Chanthrakumar, Swarag Thaikkandi, Zeyang Yu, Jacob M. Blank, David C. Steinberger, Yinru Long, Genevieve P. Nowicki, Todd M. Herrington, Daniel G. Dillon, Ann M. Graybiel, Michael R. Bruchas, Michael J. Frank, Diego A. Pizzagalli

## Abstract

Major depressive disorder (MDD) is highly recurrent, and psychosocial stress is a central precipitant of relapses. Daily choices require weighing potential rewards against costs, a process disrupted in MDD that may contribute to relapse. Here, we tested whether a psychosocial stressor affects neural and computational mechanisms underlying approach-avoidance decision making in individuals with remitted MDD (rMDD, N=50) and healthy controls (N=74), prioritizing pregenual anterior cingulate cortex (pgACC) and subthalamic nucleus (STN) due to their roles in value integration and decision caution, respectively. Participants performed a dynamic approach-avoidance conflict task with fMRI before and after psychosocial stress. Hierarchical Bayesian drift-diffusion modeling evaluated whether stress modulated computational and neural mechanisms, with trial-wise pgACC and STN activity entered as neural regressors on evidence accumulation and decision threshold, respectively. Before stress, groups did not differ in any decision parameter, but pgACC activity contributed to evidence accumulation toward avoidance in controls only. After stress, rMDD showed larger increases than controls in the impact of both reward and aversiveness on accumulation toward approach and avoidance, respectively, predominantly for net-aversive offers. Furthermore, in rMDD, stress rendered both approach and avoidance decisions more sensitive to pgACC activity, and STN activity was more strongly coupled to the decision threshold than in controls. Finally, stress-induced increases in reward sensitivity in rMDD predicted one-year course of depressive symptoms. Thus, MDD remission is not characterized by a generalized blunting of decision-making, but by a latent circuit difference that acute stress reorganizes and predicts future symptoms.

---

Major depressive disorder (MDD) is among the most recurrent psychiatric conditions, with relapse risk rising with each successive episode^1,2^. Among factors precipitating recurrence, psychosocial stress is among the most robust^3^. Cognitive^4^, physiological^5^ and neural^6–8^ evidence indicates that individuals with remitted MDD (rMDD) remain particularly sensitive to stress, yet the computational and circuit-level processes underlying this vulnerability are largely unknown. Identifying these mechanisms is important because they may represent intervention targets to build resilience and reduce recurrence risk.

One hallmark of MDD is an imbalance between blunted sensitivity to reward and heightened sensitivity to threat/cost^9,10^. Approach-avoidance (Ap-Av) conflict paradigms are particularly well suited to probe this imbalance because they simultaneously present potential gains against costs, capturing the motivational dysregulation that characterizes MDD, particularly within fronto-cingulo-striatal circuits^11,12^. Such conflict is resolved by competing approach and avoidance signals rather than by collapsing reward and cost into a single net value, so the critical computation lies outside classical value-integration regions. The ventromedial prefrontal cortex, for instance, encodes a domain-general subjective value signal that has already collapsed reward and cost into a single quantity^13^. In turn, preclinical work points to a set of regions that arbitrate the competition itself. Specifically, studies in macaques elucidated that microstimulation of the pregenual anterior cingulate cortex (pgACC) biases cost-benefit decisions toward avoidance^14^, and that pgACC decision-period activity follows caudal orbitofrontal signals carrying both appetitive and aversive information^15^. Similarly, studies in mice demonstrated that a medial prefrontal-to-striosomal striatal pathway is causally necessary for resolving cost-benefit conflict decisions^16^.

Critically, such circuitry is also where MDD pathophysiology and stress effects converge. Thus, human neuroimaging has identified abnormal pgACC and striatal activation during Ap-Av conflict in MDD^17^ – regions prominently recruited during acute stress^18,19^. Moreover, acute stress reduces reward responsiveness in humans, most markedly in anhedonic individuals^20–22^. As such, rMDD individuals may be particularly vulnerable to stress-induced disruption of these processes. In line with this, rMDD individuals show altered fronto-cingulo-striatal function, including potentiated striatal responses during psychological stress^6^ and blunted behavioral and neural responsiveness to reward^23,24^. Of note, the degree of stress-induced impairment in fronto-cingulate function prospectively predicted higher stress and depressive symptoms 18 months later in rMDD^7^. Collectively, these findings indicate that fronto-cingulo-striatal abnormalities relevant to cost-benefit decision-making can persist into remission in a latent form, which might become manifest under stress. Which computational processes might drive this vulnerability and their specificity to rMDD remain unknown.

Establishing whether and how stress disrupts cost-benefit decision-making in rMDD requires approaches that go beyond choice proportions and reaction times. Reduced approach behavior alone can arise from multiple factors. It could reflect lower sensitivity to offer value, a more cautious or more urgent decision policy, an *a priori* bias toward avoidance, or slowed sensorimotor processing, and these engage distinct neural substrates. Sequential sampling models resolve this ambiguity by decomposing choices and response times into latent processes that map onto each of these factors, including the rate of evidence accumulation (drift rate), the decision threshold, a starting-point bias, and non-decision time^25^. In model variants where the threshold declines over the course of a trial, the rate of that decline captures the growing urgency to commit^26^. Because these processes can be estimated as a function of offered incentives and trial-wise neural signals, they identify the computations and brain circuit affected by stress.

Prior work using such a framework has established that benefits and costs push evidence accumulation in opposite directions, with larger benefits driving the drift rate toward approach and larger costs toward avoidance. Critically, drift rate varies with underlying fronto-cingulo-striatal signals of benefits and costs, including the pgACC^17,27^. Decision policy, by contrast, has been linked to a distinct circuit. In particular, the mediofrontal cortex and the subthalamic nucleus (STN) track decision conflict and adjust the decision threshold, determining how much evidence is required before a choice is made and how rapidly that requirement declines within a trial^28–31^. Subthalamic stimulation causally modulates this threshold in humans^28,32–34^. Of note, drift rate and decision threshold also track anhedonic and depressive symptoms during Ap-Av conflict^26^. Whether acute stress selectively disrupts these processes in rMDD, through which computational parameters, and whether the degree of disruption relates to how symptoms develop over time, remains to be established.

To fill these gaps, we combined fMRI, a validated acute psychosocial stress induction (Maastricht Acute Stress Test, MAST)^35^, and a neurocomputational approach that combines drift diffusion modeling with activity in two *a priori* brain regions, the STN and pgACC, to characterize Ap-Av conflict decision-making in unmedicated individuals with rMDD and HC. Acute stress served as an experimental probe because it reliably activates the HPA axis and alters fronto-cingulo-striatal function^7,18,19,36^, offering a controlled model of how stress engages the circuits implicated in rMDD – potentially unveiling vulnerabilities. Participants completed a human adaptation of the macaque Ap-Av conflict task^14^ during fMRI before and after the stress induction. Participants were then followed for one year, with clinician-rated and self-reported symptom assessments at 6 and 12 months. We integrated single-trial BOLD signals as neural regressors to test whether stress differentially modulated brain-behavior coupling in rMDD relative to controls. This provides a direct test of whether latent circuit abnormalities in rMDD manifest selectively under stress as altered neurocomputational regulation of motivational conflict, and whether the magnitude of that stress-induced change relates to the subsequent course of depressive symptoms.

Computationally, given the blunted reward and heightened cost sensitivity that characterize MDD, we expected acute stress to shift evidence accumulation in rMDD toward avoidance, and to alter decision policy, both the evidence required to commit to a choice and how rapidly that requirement declines. Neurally, we expected stress to dysregulate pgACC coupling with evidence accumulation in rMDD relative to HC, and to alter STN coupling with the decision threshold and its rate of collapse. Because pgACC arbitrates the competition between appetitive and aversive signals rather than tracking either alone, we further hypothesized its contribution to depend on whether an offer was net rewarding or net aversive. Clinically, we expected that the magnitude of stress-induced changes would relate to future depressive symptoms. To foreshadow the main findings, we found that rMDD is characterized by a stress-dependent reorganization of how motivational and cognitive-control circuits shape evidence accumulation with consequences for clinical course, with such reorganization predicting symptoms 1 year later.

## Results

### Stress induction

Participants completed a dynamic approach-avoidance conflict task before and after acute stress induction, with choices and response times modeled using a hierarchical Bayesian drift-diffusion model with collapsing bounds (see **Supplementary Fig. 1-3** for study design and procedure, task, and full model specification, respectively). The MAST successfully induced robust physiological and subjective stress responses in both groups. Salivary cortisol showed a significant main effect of *Time* (F(2.80,341.2)=29.55, p<0.001, partial η²=0.20, **Supplementary Fig. 4**), with no group differences (see **Supplement** for details). Both groups showed stress-induced increases in negative affect and decreases in positive affect. However, the rMDD group reported higher overall negative affect (F(1,122)=6.66, p=0.011, partial η²=0.05) and a significantly stronger stress-induced increase in negative affect (*Group*×*Time*: F(1,122)=5.76, p=0.018, partial η²=0.05, **Supplementary Fig. 5**). The rMDD group also appraised the stressor as more threatening (*Group*×*Time*: F(1,122)=6.56, p=0.012, partial η²=0.05, **Supplementary Fig. 6**). Further, Visual Analogue Mood Scale (VAMS) ratings confirmed stress-related affective changes in both groups, with the rMDD group again reporting greater overall negative affect and a stronger stress-induced increase in hostility than HC (**Supplementary Fig. 7**).

### Computational model

Model selection was performed on a behavioral-only model paralleling our prior work in an independent (clinically unselected) sample^26^. Replicating those findings, the best fitting model was a modified hierarchical DDM in which drift rate was modulated by offered reward and aversion and the decision threshold collapsed linearly over the trial (Angle model; **Figure 1**; see the *Behavioral-Only Model Results section* in the **Supplementary Information**, with parameter estimates in **Supplementary Table 1**, key parameter effects in **Supplementary Fig. 8**, and posterior predictive checks in **Supplementary Fig. 9**). Next, single-trial BOLD signals extracted from the pgACC and STN were added as regressors on drift rate and on the decision threshold and its rate of collapse, respectively, yielding the neural model (**Figure 1, Supplementary Fig. 3)**. Estimates of the value-integration parameters were consistent across the behavioral-only and neural models, indicating that the inclusion of trial-wise BOLD regressors did not alter how offered reward and aversion entered evidence accumulation (**Supplementary Table 1** and **Supplementary Fig. 8**).

**Figure 1.**
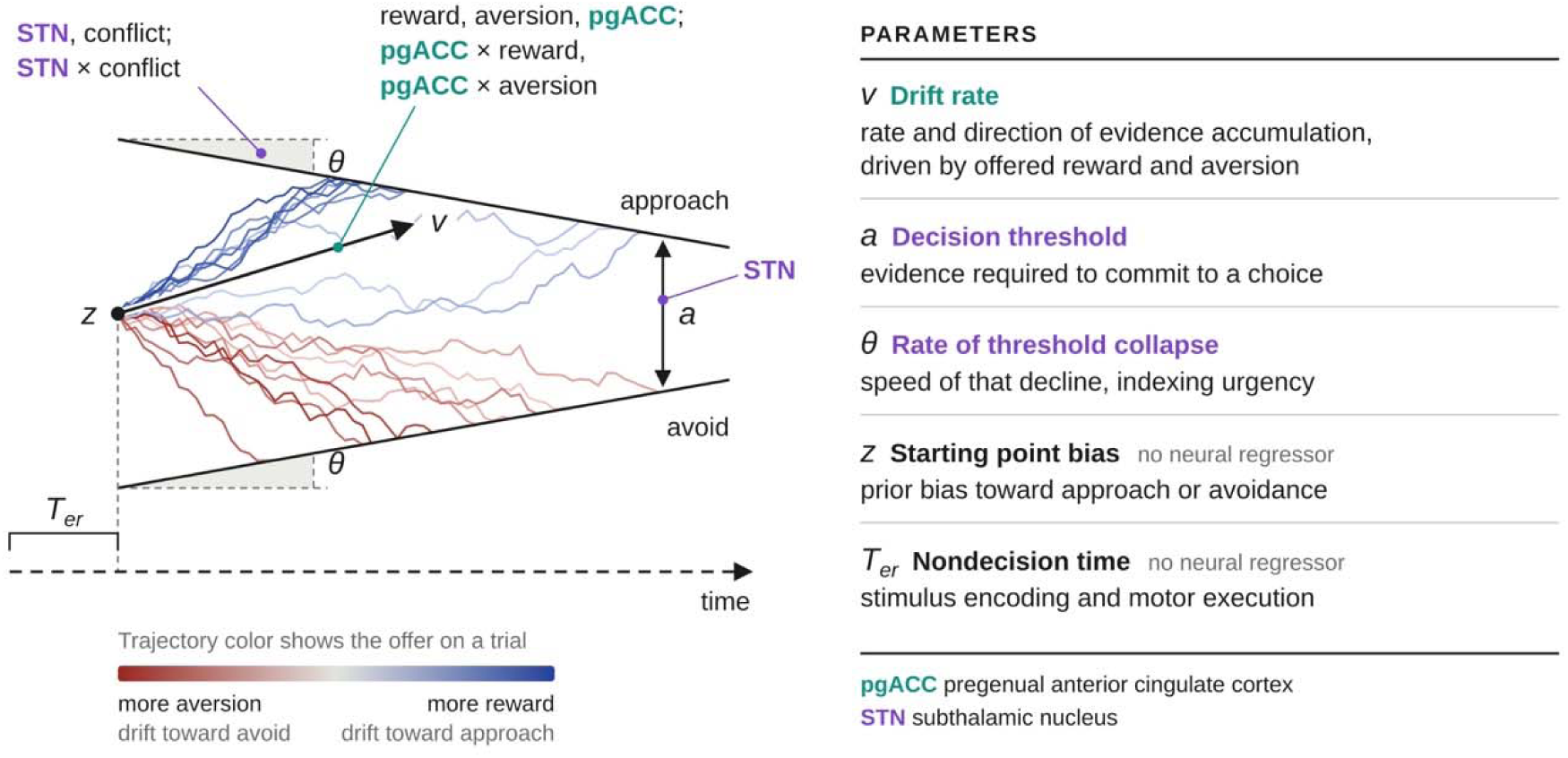
Model structure and neural modulation. **Left**: illustration of the drift diffusion model with a linearly collapsing decision threshold, adapted to the approach-avoidance conflict task. The model was used to estimate how trial-by-trial changes in offered reward and aversion, as well as their neural correlates, altered the speed and sign of evidence accumulation (controlled by the drift rate, v) toward approaching (upper bound) or avoiding (lower bound) an offer. Trajectory color indicates the offer on a trial, from more aversion (red, drift toward avoidance) to more reward (blue, drift toward approach). The decision threshold (a) measures the distance between the two bounds and thus the evidence required to commit to a choice, the rate of threshold collapse (θ) measures how rapidly that requirement declines within a trial, the starting point bias (z) measures a priori tendencies to approach or avoid offers, while the nondecision time (Ter) parameter captures time spent on stimulus encoding and motor response. Annotations indicate where trial-wise BOLD activity entered each parameter, with pgACC in teal and STN in purple. Within an annotation, terms before the semicolon are main effects and terms joined by × are interactions. **Right**: Summary of the model parameters and their interpretation. Parameter names are colored by the region whose activity modulated them, and names in black had no neural regressor. Trial-wise pgACC activity modulated the drift rate both directly and through separate interactions with offered reward and offered aversion, and all drift-rate terms were additionally estimated by domain. Trial-wise STN activity modulated the decision threshold, and both STN activity and trial-wise conflict modulated the rate of threshold collapse, individually and in interaction. Starting point bias varied by Pavlovian response congruency. All terms were estimated for both stress conditions within a single model, with post-stress as the reference. Model equations are given in **Online Methods** (Computational Modeling Approach), and the full graphical model is shown in **Supplementary** Fig. 3.

Based on the neural model, we first characterized decision dynamics before stress. We then examined group-specific differences in how stress alters decision dynamics, value integration, and their neural correlates. Further details are reported in **Supplementary Tables 2** and **3**. Interaction terms of fourth order and above were also tested but are omitted from the reported model. Estimates from the full model that retains them are summarized in **Supplementary Table 4**, posterior predictive checks for both are shown in **Supplementary Figs. 10** and **11**, and their parameter estimates are compared in **Supplementary Figs. 12-15**.

### Both groups show largely comparable approach-avoidance dynamics before stress

A characteristic of our Ap-Av conflict paradigm (see **Online Methods**) is that offers differ in whether their overall magnitude (= reward minus cost) is positive or negative, which we refer to as domain^26^. In general, larger rewards induce greater drift rate, leading to faster and more frequent approaches, and larger aversive offers induce greater drift toward avoidance^27^, but the relative impact of rewards and aversion can differ by domain^26^. In the negative domain (net-aversive offers), both groups showed positive impacts of reward (HC: β=0.662, 95%-HDI=[0.588, 0.738]; rMDD: β=0.619, 95%-HDI=[0.521, 0.718]) and negative impacts of aversion (HC: β=−0.658, 95%-HDI=[−0.731, −0.586]; rMDD: β=−0.619, 95%-HDI=[−0.705, −0.535]), with no credible group differences in either of these effects (**Supplementary Table 2**). Offers in the positive domain elevated drift rate toward approach in both groups (HC: β=1.067, 95%-HDI=[0.872, 1.259]; rMDD: β=1.082, 95%-HDI=[0.881, 1.284]), replicating the expected domain effect^26^ without a credible group difference. The influence of additional units of reward and aversion was attenuated in the positive domain, in both groups and without a credible group difference (**Supplementary Table 2**).

Previous work has documented effects of conflict on the collapsing decision threshold^30^, whereby higher conflict (here, smaller difference between reward and aversion within a trial’s offer) leads to a prolonged (slower) rate of decision threshold collapse. This prolonging effect was credible in rMDD but not in HC, with the group difference approaching our criterion (HC: β=−0.003, 95%-HDI=[−0.023, 0.016]; rMDD: β=−0.025, 95%-HDI=[−0.042, −0.007], posterior probability PP[rMDD>HC]=0.052). Before stress, groups did not differ credibly on any other model parameter (**Supplementary Table 2**).

### Pre-stress group differences reveal latent motivational neural differences in rMDD

Next, we asked whether groups differed in the neural circuits supporting decision-making by focusing on how trial-by-trial pgACC and STN activity related to the latent cognitive processes indexed by the model parameters (see **Online Methods** and **Figure 1**). Credible pre-stress group differences were confined to pgACC–drift couplings. Specifically, on net-aversive offers, higher pgACC activity was associated with more negative drift rate (i.e., stronger accumulation toward avoidance) in the HC but not rMDD group (HC: β=−0.076, 95%-HDI=[−0.126, −0.028]; rMDD: β=−0.007, 95%-HDI=[−0.063, 0.049], PP[rMDD>HC]=0.968; see **Supplementary Table 2** for all pre-stress parameter estimates). These results suggest that in HC, pgACC activity contributes a subjective aversive signal beyond the objective offer. However, as objective aversion increased, higher pgACC activity attenuated the impact of aversion on drift rate in the HC group, again with no effect in the rMDD group (HC: β=0.050, 95%-HDI=[0.010, 0.090]; rMDD: β=−0.026, 95%-HDI=[−0.072, 0.021], PP[rMDD>HC]=0.010). This interaction is consistent with a sublinear combination of pgACC- and aversion-related influences on evidence accumulation. Notably, this latter effect was the strongest group difference observed before stress. In HC, the negative pgACC–drift coupling was specific to net-aversive offers and was attenuated on net-rewarding offers, with no credible group difference in this domain dependence (**Supplementary Table 2**). Together, two credible group differences suggest that rMDD is characterized not by a global blunting of approach-avoidance sensitivity or evidence accumulation, but by a specific decoupling of pgACC from the drift-rate computation.

The subthalamic terms showed no comparable group difference. Consistent with prior work^29,30,32,34^, however, higher STN activity was linked to higher overall decision threshold and slower (more prolonged) collapse in both groups, with no credible group difference (**Supplementary Table 2**). This pattern is consistent with a role for the STN in prolonging evidence accumulation and indicates that the pre-stress group difference does not reflect a generic difference in how the subthalamic nucleus regulates decision policy.

### Stress alters value integration and decision policy differently by group

We next asked how stress altered evidence accumulation, and whether it did so differently by group. We focus first on value integration (drift rate) and then turn to decision policy (overall threshold and rate of threshold collapse).

Stress altered drift rate differently across groups, revealing an asymmetry between reward- and aversion-related processing. On net-aversive offers, reward sensitivity increased credibly in both groups following stress, but this increase was larger in rMDD than HC (post minus pre-stress, rMDD: β=0.188, 95%-HDI=[0.124, 0.252]; HC: β=0.098, 95%-HDI=[0.044, 0.152]; PP[rMDD>HC]=0.982). This direction is consistent with evidence that acute stress can actually enhance reward-related learning and selection^37^. On these same offers, aversion sensitivity also became stronger (more negative, i.e., toward avoidance) in rMDD but not HC (rMDD: β=−0.106, 95%-HDI=[−0.178, −0.032], HC: β=−0.019, 95%-HDI=[−0.083, 0.042]; PP[rMDD>HC]=0.035). Stress therefore increased the weight of both reward and cost on evidence accumulation for net-aversive offers, and in both cases credibly more in rMDD than HC, with the cost effect credible only within the rMDD group. These stress-induced changes in value integration were themselves domain-dependent. On net-rewarding offers, this increase in reward sensitivity was credibly smaller in rMDD but not in HC (difference from net-aversive offers, rMDD: β=−0.228, 95%-HDI=[−0.330, −0.120]; HC: β=−0.084, 95%-HDI=[−0.190, 0.020]; PP[rMDD>HC]=0.029). The stress-induced change on net-rewarding offers is the sum of the change on net-aversive offers and this difference, and it was close to zero in both groups (rMDD: β=−0.040; HC: β=0.014). Stress therefore increased reward sensitivity when costs outweighed rewards and left it largely unchanged when rewards outweighed costs, and this domain dependence was sharper in rMDD.

Stress also changed how pgACC activity related to evidence accumulation, and it did so in rMDD only (**Figure 2**). Whereas before stress higher pgACC activity was associated with accumulation toward avoidance on net-aversive offers in HC but not rMDD, after stress this coupling was credible in rMDD but not in HC, with the group difference approaching our criterion (post-stress, rMDD: β=−0.079, 95%-HDI=[−0.126, −0.034]; HC: β=−0.027, 95%-HDI=[−0.071, 0.015]; PP[rMDD>HC]=0.052; **Figure 2b**). The underlying stress-induced change was credible in rMDD, absent in HC, and credibly different between groups (post minus pre-stress, rMDD: β=−0.073, 95%-HDI=[−0.142, −0.011]; HC: β=0.049, 95%-HDI=[−0.009, 0.107]; PP[rMDD>HC]=0.001; **Figure 2**a). On net-rewarding offers, stress shifted the coupling in the opposite direction in rMDD. Relative to net-aversive offers, stress made the coupling more positive in rMDD but not in HC (additional stress-induced change on net-rewarding offers, rMDD: β=0.174, 95%-HDI=[0.069, 0.290]; HC: β=−0.035, 95%-HDI=[−0.133, 0.060]; PP[rMDD>HC]=0.998; **Figure 2c**). After stress, this additional coupling was credible in rMDD only (rMDD: β=0.171, 95%-HDI=[0.103, 0.243]; HC: β=0.050, 95%-HDI=[−0.018, 0.117]; PP[rMDD>HC]=0.992; **Figure 2d**). Adding it to the coupling on net-aversive offers, the implied post-stress coupling in rMDD was negative on net-aversive offers (β=−0.079) and positive on net-rewarding offers (β=0.092), whereas in HC it was close to zero on both (β=−0.027 and β=0.023). After stress, higher pgACC activity in rMDD was therefore associated with stronger accumulation toward avoidance when costs outweighed rewards and toward approach when rewards outweighed costs, a pattern not seen in HC.

**Figure 2.**
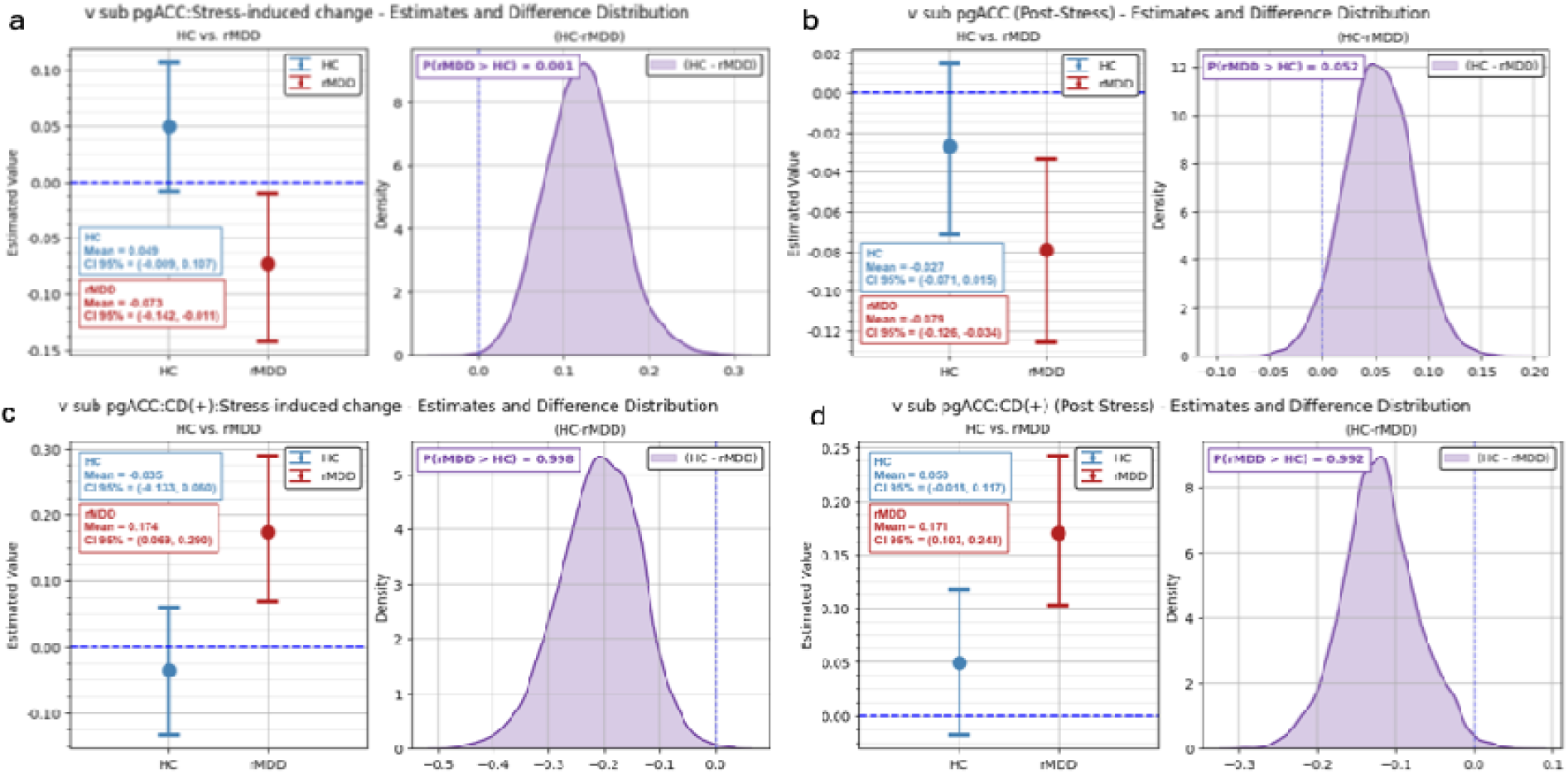
Stress altered pgACC–drift rate coupling in rMDD but not in HC. (a) Stress-induced change (post minus pre-stress) in the coupling on net-aversive offers. (b) Post-stress coupling on net-aversive offers. (c) Additional stress-induced change on net-rewarding offers, relative to net-aversive offers. (d) Additional post-stress coupling on net-rewarding offers, relative to net-aversive offers. In each panel, the left plot shows group posterior means with 95% highest density intervals, and the right plot the posterior distribution of the group difference (HC - rMDD), with the posterior probability that the estimate is larger in rMDD than in HC, PP[rMDD>HC].

Stress also altered the pgACC modulation of aversion sensitivity on net-aversive offers, the strongest group difference before stress. This interaction shifted credibly in rMDD but not in HC (post minus pre-stress, rMDD: β=0.067, 95%-HDI=[0.018, 0.117]; HC: β=0.002, 95%-HDI=[−0.041, 0.042]; PP[rMDD>HC]=0.978), so that the groups no longer differed after stress (PP[rMDD>HC]=0.370; **Supplementary Table 3**). For this interaction, rMDD therefore moved toward the pattern shown by HC before stress, and the groups converged rather than diverged.

Beyond value integration, stress may also shift the decision policies governing caution and urgency. Specifically, stress reduced the decision threshold in both groups, with no credible group difference, indicating a shared shift toward urgency (faster, less cautious decision strategies; HC: β=−0.134, 95%-HDI=[−0.226, −0.047]; rMDD: β=−0.149, 95%-HDI=[−0.259, −0.042]; **Supplementary Table 3**). Stress also changed how the decision threshold tracked STN activity, and it did so differently by group. Whereas the two groups did not differ before stress as reported above, post-stress the coupling between STN activity and decision threshold was substantially stronger in rMDD than HC (HC: β=0.024, 95%-HDI=[−0.001, 0.048]; rMDD: β=0.081, 95%-HDI=[0.051, 0.107]; PP[rMDD>HC]=0.999, **Figure 3**). This divergence emerged from stress-induced changes that were credibly different between groups even though neither was credible on its own, with the coupling shifting upward in rMDD and downward in HC (HC: β=−0.019, 95%-HDI=[−0.053, 0.020]; rMDD: β=0.031, 95%-HDI=[−0.012, 0.075]; PP[rMDD>HC]=0.952). Thus, after stress, rMDD required more evidence to commit as STN activity increased, whereas in HC this relationship was no longer credible.

**Figure 3.**
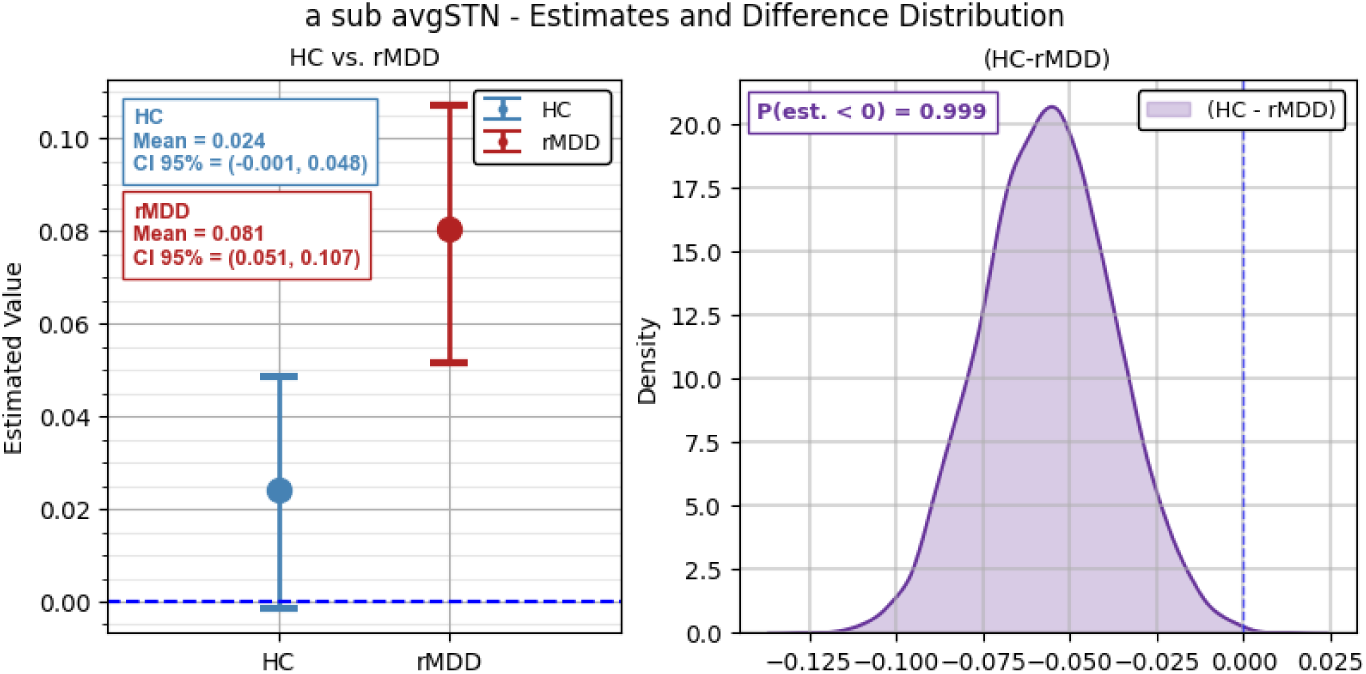
Credible group difference in the STN sensitivity of the decision threshold post-stress. Plots on the left reflect the posterior estimates from MCMC trace samples for each group, while the plot on the right reflects the posterior distribution of the sampled difference. Post-stress, higher STN activity was associated with a higher decision threshold in rMDD, whereas this relationship was no longer credible in HC (PP[rMDD>HC]=0.999).

Stress also reorganized the rate of threshold collapse, and here the groups diverged in two ways. First, only rMDD showed a stress-induced increase in the rate of threshold collapse (again, increasing urgency) specifically on high-conflict trials, while HC showed no such effect (HC: β=0.002, 95%-HDI=[−0.022, 0.025]; rMDD: β=0.033, 95%-HDI=[0.007, 0.056]; PP[rMDD>HC]=0.961). Indeed, the conflict-related prolongation observed before stress was no longer credible in rMDD after stress (β=0.008, 95%-HDI=[−0.012, 0.028]). Second, stress also changed how the rate of threshold collapse tracked STN activity, and it did so differently by group. In HC, higher STN activity continued to prolong collapse (post-stress β=−0.037, 95%-HDI=[−0.055, −0.020]), whereas in rMDD this effect was no longer credible (β=0.014, 95%-HDI=[−0.008, 0.033]). Thus stress shifted the impact of STN activity toward faster collapse in rMDD, whereas in HC it did not change (rMDD: β=0.038, 95%-HDI=[0.008, 0.068]; HC: β=−0.007, 95%-HDI=[−0.030, 0.018]; PP[rMDD>HC]=0.987). Consequently, the two groups differed credibly in post-stress STN-collapse coupling (PP[rMDD>HC]>0.999, **Figure 4**). Collectively, these two effects describe single change. Pre-stress, two independent influences prolonged the collapse of the decision threshold in rMDD, one driven by conflict and one by STN activity. After stress neither remained credible, whereas in HC the STN influence was preserved and the conflict influence was not credible at either timepoint. Thus, stress abolished, selectively in rMDD, the mechanisms that prolong decisions when evidence is conflicting.

**Figure 4.**
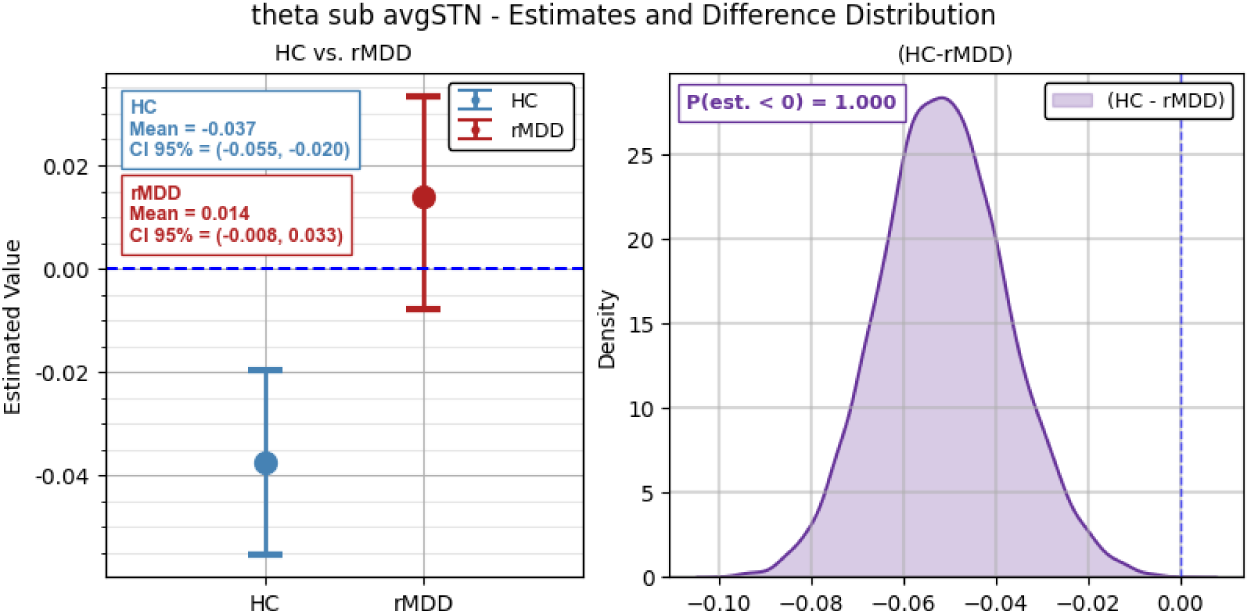
Highly credible difference between rMDD and HC in the STN sensitivity of the rate of threshold collapse post-stress. Plots on the left reflect the posterior estimates from MCMC trace samples for each group, while the plot on the right reflects the posterior distribution of the sampled difference. HC show slower collapse with increasing STN activity, whereas in rMDD this relationship is no longer credible after stress. The group difference is highly credible (PP[rMDD>HC]>0.999).

Group differences in the effects of stress were confined to drift rate and decision threshold. Stress also shortened nondecision time and introduced a Pavlovian congruency effect on the starting point bias, but these effects were shared across groups, with no credible group differences (**Supplementary Results** and **Supplementary Table 3**).

### Stress-induced reward sensitivity is associated with the subsequent course of depressive symptoms in rMDD

Because stress increased reward sensitivity credibly more in rMDD than in HC, we asked whether its magnitude was related to how depressive symptoms developed over the following year. Clinician-rated symptoms were assessed with the Hamilton Depression Rating Scale (HAMD) at baseline and again at 6 months, in 47 (of 50) rMDD and 62 (of 74) HC participants, and at 12 months, in 45 rMDD and 60 HC participants. HAMD scores across all three visits were modeled using generalized estimating equations, entering the stress-induced change in reward sensitivity on net-aversive offers, standardized across th whole sample, together with its interaction with time. Within rMDD, a larger stress-induced increase in reward sensitivity was associated with a steeper rise in symptoms (b=0.108 HAMD points per month per SD, 95% CI [0.002, 0.214], p=0.045), corresponding to a difference of 1.30 points [0.03, 2.57] over twelve months. No such association was present in HC (b=−0.013, 95% CI [−0.052, 0.026], p=0.52), and the interaction of predictor, time and group was significant (b=0.121, 95% CI [0.008, 0.234], p=0.036; **Figure 5**). The corresponding parameter for net-rewarding offers showed a similar but weaker association that did not reach significance within rMDD (b=0.080, 95% CI [−0.009, 0.168], p=0.078) or for the group interaction (b=0.084, 95% CI [−0.005, 0.173], p=0.065). Full coefficients for these models are given in Supplementary Table 5.

**Figure 5.**
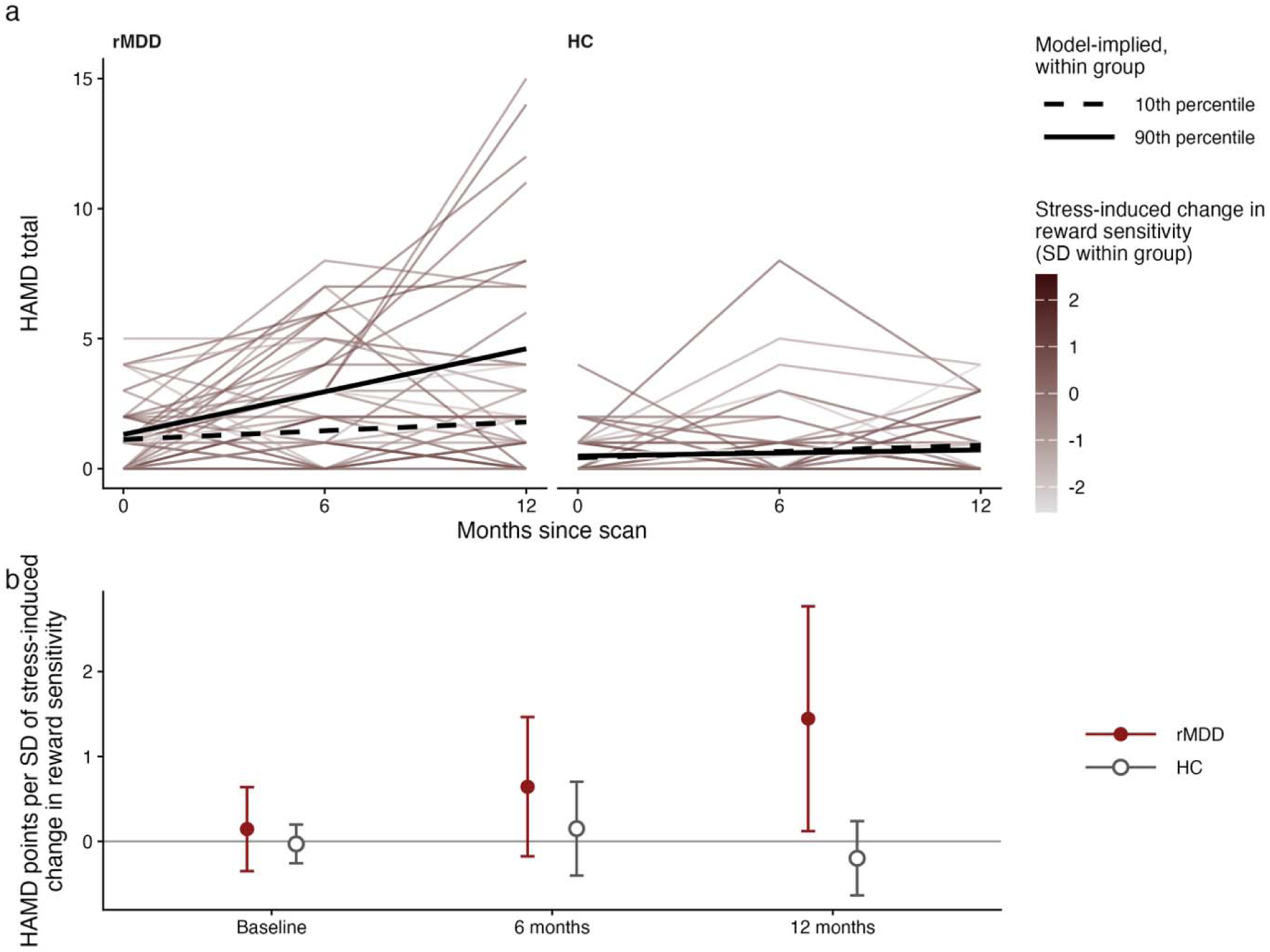
Within rMDD, a larger stress-induced increase in reward sensitivity was associated with a steeper rise in depressive symptoms over the following year. (a) Observed HAMD totals at baseline, 6 and 12 months for every participant, shaded by each participant’s stress-induced change in the influence of reward magnitude on drift rate for net-aversive offers, expressed relative to the mean and standard deviation of their own group. Bold lines represent the trajectory implied by the generalized estimating equation reported in the text, drawn at the 10th and 90th percentile of the predictor within each group. The predictor entered the model continuously and was standardized across the whole sample, so the coefficients quoted in the text are per sample-wide standard deviation. (b) Association between the same predictor and HAMD at each visit, with visit as a factor, expressed in HAMD points per sample-wide standard deviation of the predictor, with 95% confidence intervals from robust standard errors. Within rMDD the association was absent at baseline, intermediate at 6 months and present at 12 months, with no corresponding pattern in HC at any visit.

The association emerged over the course of follow-up rather than being present from the outset. Refitting the same model with visit entered as a factor (**Figure 5b**), the association within rMDD was absent at baseline (b=0.14, 95% CI [−0.35, 0.64], p=0.57), intermediate at 6 months (b=0.64, 95% CI [−0.18, 1.46], p=0.12) and present at 12 months (b=1.45, 95% CI [0.12, 2.77], p=0.032), with no corresponding pattern among HC at any visit. Analyzing each follow-up visit separately with baseline symptoms as a covariate yielded the same pattern, with no group difference in the association at 6 months (b=0.43, F(1,104)=0.75, p=0.39) and a significant one at 12 months (b=1.57, F(1,100)=4.95, p=0.026; **Supplementary Table 6** and **Supplementary Fig. 16**).

Notably, this association was confined to depression severity. Self-reported depressive symptoms showed the same pattern (BDI-II: b=0.155, 95% CI [0.003, 0.307], p=0.046; QIDS-SR16: b=0.076, 95% CI [0.002, 0.149], p=0.043), whereas anhedonia (SHAPS, b=−0.090, 95% CI [−0.366, 0.186], p=0.52), perceived stress (PSS-14, b=0.054, 95% CI [−0.210, 0.318], p=0.69) and quality of life (Q-LES-Q-SF, b=−0.114, 95% CI [−0.329, 0.101], p=0.30) showed no relationship (**Supplementary Table 7**).

## Discussion

We combined fMRI, an optimized task back-translated from non-human primate research, a validated acute psychosocial stressor, and hierarchical Bayesian modeling of approach-avoidance conflict to examine how acute stress alters cost-benefit decision-making in rMDD, and whether such metrics predicted future symptoms. The principal finding was a difference in motivational circuitry that behavior alone could not detect, and that acute stress reorganized. Before stress, individuals with rMDD and never-depressed controls were closely matched on the behavioral parameters of the decision process, including the sensitivity of evidence accumulation to reward and aversion and the amount of evidence required to commit to a choice. Yet, the pgACC was coupled to evidence accumulation in HC but not rMDD. Stress then reorganized such coupling and, separately, abolished the two influences (one driven by conflict itself and one by subthalamic activity) that had slowed the collapse of the decision threshold under conflict in rMDD. Finally, among fully remitted individuals with past MDD, the size of the stress-induced increase in reward sensitivity on net-aversive offers also tracked the one-year course of depressive symptoms. Collectively, these findings indicate that rMDD was not characterized by a generalized blunting of decision-making but by a latent difference in motivational circuitry that behavior does not express before the stressor, that stress reorganizes, and that relates to the later course of symptoms.

In detail, before acute stress, groups differed in brain-behavior coupling. Specifically, higher pgACC activity was associated with a more negative drift rate on net-aversive offers (i.e., accumulation biased toward avoidance) in HC and showed no such association in rMDD; similarly, pgACC activation modulated the influence of aversion on evidence accumulation in HC only. Both effects were present while no behavioral parameter of the decision process differed credibly between groups. As such, the difference manifested at the level of brain-behavior coupling rather than behavior itself. These findings also mirror prior findings of (1) reduced avoidance-related pgACC activation in currently depressed individuals performing a similar task^17^, and (2) pgACC microstimulation in macaques biasing cost-benefit decisions toward avoidance^14^; thus, a region with causal control over conflict decisions was the one decoupled from the decision computation in remitted depression.

Under stress, the groups diverged asymmetrically in how reward and aversion impacted evidence accumulation. On net-aversive offers, the stress-induced increase in reward sensitivity was credibly larger in rMDD than HC, and aversion sensitivity became credibly more negative in rMDD than HC, meaning aversion pushed evidence accumulation further toward avoidance in rMDD. The aversion result was hypothesized and is aligned with accounts of heightened cost sensitivity in depression. The direction of the reward effect runs, however, counter to a substantial literature showing that acute stress blunts reward responsiveness, particularly in individuals with elevated anhedonia^22^. However, these literatures likely probe different constructs. The blunting literature is anchored on response bias in probabilistic reward tasks, which indexes the capacity to modulate behavior by reinforcement history. Conversely, drift-rate reward sensitivity indexes how strongly an offered reward pulls evidence accumulation toward approach. These are dissociable, and how to reconcile stress-enhanced reward salience with blunted reward valuation remains unresolved^38^. As such, we observed an increase in reward salience and not reward valuation. In the same cohort studied here, rMDD participants retained heightened sensitivity to high-value rewards during effort-based decision-making^39^. Across two paradigms, then, reward sensitivity appears preserved and, in some respects, heightened in remission, and stress-vulnerable rather than blunted.

The stress-induced gain in reward sensitivity was not uniform across offers. In both groups it was largely confined to net-aversive offers, those on which the cost outweighed the reward, and the offset that removed it on net-rewarding offers was credibly larger in rMDD than HC. Stress therefore did not simply raise the weight of reward on evidence accumulation. It raised that weight only where reward had to compete against a cost that exceeded it. Critically, in rMDD, the degree of stress-induced shift of weighting of reward against costs tracked clinical course. Thus, individuals whose reward weighting shifted most under stress showed the steepest rise in clinician-rated and self-reported depressive symptoms over the ensuing year. This weighting reflects the pull of an offered reward on evidence accumulation, and not the capacity to modulate behavior by reinforcement history, in which blunting has been repeatedly reported in current and past MDD^22^. Further highlighting dissociations with prior findings linking stress-blunted reward responsiveness to anhedonia, this association did not extend to anhedonia, perceived stress or quality of life.

In rMDD, stress also introduced a change of sign in pgACC–drift coupling that, to our knowledge, has not been reported before. Because drift rate is already negative on net-aversive offers, the more negative coupling indicates stronger accumulation toward avoidance, consistent with heightened threat sensitivity, whereas the positive coupling on net-rewarding offers indicates stronger accumulation toward approach, consistent with stress-potentiated reward engagement. One interpretation is that, under stress, pgACC activity in rMDD reflected the subjective utility of the action favored by the offer, strengthening avoidance when costs dominated and approach when rewards dominated. Because this domain dependence had been present in HC and absent in rMDD before stress, stress introduced it in rMDD rather than inverting an existing one, so that under stress the pgACC contribution to decisions in rMDD differed in kind and not only in degree (see **Supplementary Discussion** for alternative interpretations of this pattern, and its relation to earlier work with this task).

Stress also altered how decisions were committed. Pre-stress, two independent influences prolonged the collapse of the decision threshold in rMDD, one driven by trial-wise conflict and one by subthalamic activity. After stress neither remained credible, whereas in HC the subthalamic influence was preserved and the conflict influence had not been credible at either timepoint. Post-stress, subthalamic activity was also more strongly coupled with the overall decision threshold in rMDD than in HC. Stress therefore removed, specifically in rMDD, the mechanisms that prolong decisions when evidence is conflicting, while preserving and indeed strengthening the mechanism that raises the overall evidence requirement. The subthalamic nucleus has long been described as implementing a “hold-your-horses” signal that raises the decision threshold under conflict^28,29^. The present findings do not contradict that account but indicate a state-dependent failure of one of its components in a clinical group.

Two observations constrain how these group differences should be interpreted. First, the strongest pre-stress group difference was pgACC attenuation of aversion’s influence on drift rate on net-aversive offers, present in HC but not rMDD. That group difference disappeared after stress, because rMDD moved toward the HC pattern. Stress therefore produced divergence on one pgACC term and convergence on another, and cannot be read as a general amplifier of group differences. Second, the two groups demonstrated equivalent physiological stress responses. Cortisol increased robustly and neither its magnitude nor its time course differed by group, which argues against the simplest alternative explanation that rMDD participants reacted with a stronger endocrine response rather than evaluating the challenge differently. That equivalence is itself notable, since prior work in recurrent remitted depression has reported larger cortisol responses in remitted individuals^6,8^. Appraisal did differ, with rMDD participants rating the stressor as more threatening and reporting a larger increase in negative affect despite a comparable endocrine response, which supports the notion that the group difference emerged in how the challenge was evaluated rather than in how strongly the HPA axis was engaged.

Several limitations should be noted. First, subthalamic signal at 3 Tesla potentially reflects a mixture of the nucleus and surrounding tissue^40^. Second, due to model complexity and computational constraints related to convergence, we considered only two brain regions so it is possible that other brain regions might show stress and group effects. Third, the domain-dependent effects sit on net-rewarding offers, where parameter recovery in this model family has previously been less favorable^26^, and we therefore weigh them less heavily than the net-aversive and pre-stress results. Fourth, symptom levels during follow-up remained low: mean HAMD in rMDD rose from 1.14 at baseline to 3.20 at 12 months, still below clinical levels (a score of 7 defines remission), so our findings concern variation within a subclinical range rather than a return of clinically significant depression. Finally, the sample was unmedicated and in remission, so the findings cannot be extrapolated to current MDD.

Collectively, these findings indicate that remitted depression is not a return to intact decision-making. Before being exposed to an acute stressor, the parameters of the decision process were closely matched between rMDD and never-depressed controls, yet pgACC activity shaped evidence accumulation in controls only. Acute stress then reorganized value weighting, introducing a domain-dependent change of sign in pgACC–drift coupling across net-aversive and net-rewarding offers, and removing the conflict- and STN-related brakes that had prolonged commitment under conflict. Critically, the stress-induced increase in reward sensitivity on net-aversive offers was associated with the course of depressive symptoms in rMDD over the following year, raising the possibility that stress reactivity unveiled residual vulnerability. Clinically, relapse-prevention approaches may need to stabilize cost-reward weighting and restore pgACC and STN control under stress, rather than only assessing function in rest conditions where the risk-relevant change is not expressed.

## Online Methods

### Participants

A total of 129 unmedicated participants were enrolled, including 79 healthy controls (HC) and 50 euthymic individuals with rMDD. Participants were recruited through local advertisements and clinical referrals. The rMDD group met the criteria for prior MDD but had been in remission for at least six months at the time of testing. Exclusion criteria included current or past bipolar disorder, psychotic symptoms, substance use disorders within the past year, neurological disorders, and contraindications for MRI scanning. Five healthy control participants were excluded from the computational analyses, three due to missing neural data for one or both task runs, one due to a technical failure of the shock delivery during task calibration, and one due to invariant responding during the post-stress run. No participants in the rMDD group were excluded. The final sample therefore comprised 124 participants (74 HC, 50 rMDD; see **Table 1**). Study procedures were approved by the Mass General Brigham Human Research Committee (Protocol #’s: 2014P001871, 2018P000318), which serves as the Institutional Review Board for McLean Hospital. Participants provided written informed consent.

**Table 1.**
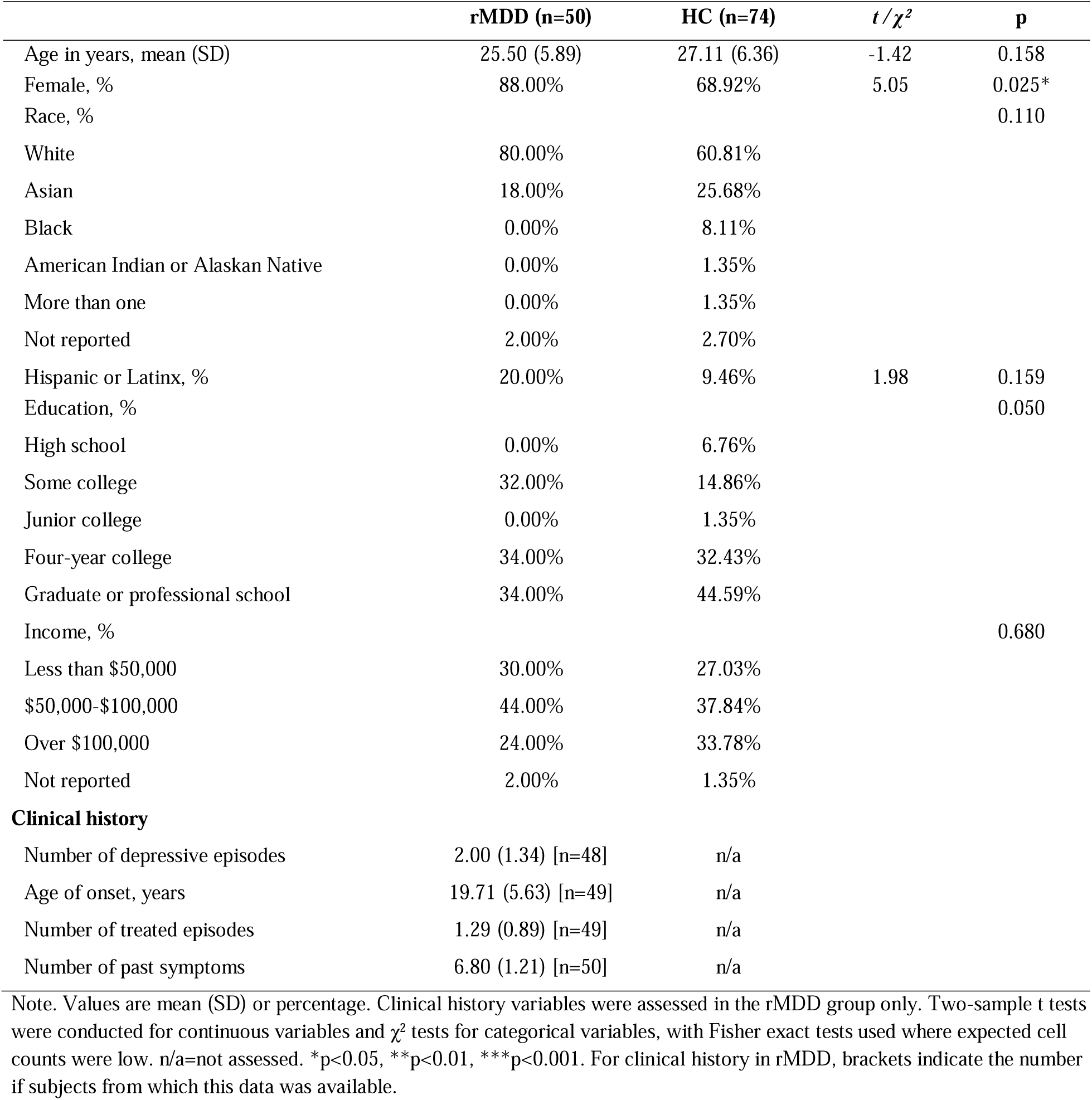
Demographic and clinical characteristics of the sample.

|  | **rMDD (n=50)** | **HC (n=74)** | ***t* / $\chi^2$** | ***p*** |
| --- | --- | --- | --- | --- |
| Age in years, mean (SD) | 25.50 (5.89) | 27.11 (6.36) | -1.42 | 0.158 |
| Female, % | 88.00% | 68.92% | 5.05 | 0.025* |
| Race, % |  |  |  | 0.110 |
| White | 80.00% | 60.81% |  |  |
| Asian | 18.00% | 25.68% |  |  |
| Black | 0.00% | 8.11% |  |  |
| American Indian or Alaskan Native | 0.00% | 1.35% |  |  |
| More than one | 0.00% | 1.35% |  |  |
| Not reported | 2.00% | 2.70% |  |  |
| Hispanic or Latinx, % | 20.00% | 9.46% | 1.98 | 0.159 |
| Education, % |  |  |  | 0.050 |
| High school | 0.00% | 6.76% |  |  |
| Some college | 32.00% | 14.86% |  |  |
| Junior college | 0.00% | 1.35% |  |  |
| Four-year college | 34.00% | 32.43% |  |  |
| Graduate or professional school | 34.00% | 44.59% |  |  |
| Income, % |  |  |  | 0.680 |
| Less than \$50,000 | 30.00% | 27.03% | | |
| \$50,000-\$100,000 | 44.00% | 37.84% | | |
| Over \$100,000 | 24.00% | 33.78% | | |
| Not reported | 2.00% | 1.35% |  |  |
| **Clinical history** |  |  |  |  |
| Number of depressive episodes | 2.00 (1.34) [n=48] | n/a |  |  |
| Age of onset, years | 19.71 (5.63) [n=49] | n/a |  |  |
| Number of treated episodes | 1.29 (0.89) [n=49] | n/a |  |  |
| Number of past symptoms | 6.80 (1.21) [n=50] | n/a |  |  |

Note. Values are mean (SD) or percentage. Clinical history variables were assessed in the rMDD group only. Two-sample *t* tests were conducted for continuous variables and $\chi^2$ tests for categorical variables, with Fisher exact tests used where expected cell counts were low. n/a=not assessed. \**p*<0.05, \*\**p*<0.01, \*\*\**p*<0.001. For clinical history in rMDD, brackets indicate the number of subjects from which this data was available.

### Experimental Design

Participants completed a dynamic approach-avoidance conflict task during fMRI in two conditions, a pre-stress condition that assessed decision-making under neutral conditions, and a post-stress condition that assessed stress-induced changes in approach-avoidance behavior. Between conditions, participants underwent an acute stress induction protocol. Clinical follow-up assessments took place 6 and 12 months after the scan, at which clinician-rated and self-report symptom measures were repeated (Table 2). For details of the full study procedure, see **Supplementary Fig. 1**.

**Table 2.** Clinical and self-report measures at baseline and at 6- and 12-month follow-ups.

| Measure | Baseline |  |  | 6-month follow-up |  |  | 12-month follow-up |  |  |
| --- | --- | --- | --- | --- | --- | --- | --- | --- | --- |
|  | rMDD | HC | *p* | rMDD | HC | *p* | rMDD | HC | *p* |
| HAMD | 1.14 (1.34) | 0.47 (0.80) | <0.001*** | 2.43 (2.27) | 0.61 (1.41) | <0.001*** | 3.20 (3.86) | 0.83 (1.12) | <0.001*** |
| QIDS-SR | 0.88 (1.25) | 0.41 (0.74) | 0.009** | 1.74 (1.74) | 0.65 (1.27) | <0.001*** | 2.16 (2.62) | 0.55 (0.87) | <0.001*** |
| BDI-II | 2.08 (2.77) | 0.74 (1.50) | <0.001*** | 4.60 (4.70) | 1.37 (2.85) | <0.001*** | 5.42 (5.93) | 1.66 (2.77) | <0.001*** |
| SHAPS | 21.48 (6.16) | 19.38 (5.79) | 0.053 | 19.91 (4.34) | 19.65 (5.50) | 0.782 | 21.29 (6.39) | 18.31 (4.62) | 0.005** |
| TEPS anticipatory | 48.38 (5.82) | 48.66 (5.71) | 0.789 | 48.64 (6.08) | 48.27 (6.74) | 0.771 | 47.95 (6.27) | 48.61 (6.35) | 0.602 |
| TEPS consummatory | 41.58 (4.25) | 40.91 (4.89) | 0.427 | 41.81 (4.54) | 41.23 (4.72) | 0.516 | 42.02 (4.23) | 41.20 (4.86) | 0.365 |
| PSS | 15.80 (7.11) | 12.11 (5.85) | 0.002** | 20.45 (7.71) | 13.94 (6.88) | <0.001*** | 19.64 (8.02) | 14.39 (6.46) | <0.001*** |
| QLESQ | 59.86 (5.58) | 63.36 (5.43) | <0.001*** | 57.00 (6.83) | 61.68 (5.76) | <0.001*** | 56.73 (7.62) | 62.10 (5.70) | <0.001*** |

Note. Values are mean (SD). All 50 rMDD and 74 HC participants were assessed at baseline, 47 and 62 attended the 6-month visit and 45 and 61 attended the 12-month visit. Not every attending participant completed every measure, so available n varied slightly by measure, from 48 to 50 (rMDD) and 74 (HC) at baseline, 47 (rMDD) and 62 (HC) at 6 months, and 44 to 45 (rMDD) and 60 to 61 (HC) at 12 months. *p* values are from two-sample *t* tests comparing groups at each visit. \**p*<0.05, \*\**p*<0.01, \*\*\**p*<0.001.
HAMD=Hamilton Depression Rating Scale; QIDS-SR16=Quick Inventory of Depressive Symptomatology, Self-Report, 16 items; BDI-II=Beck Depression Inventory-II; SHAPS=Snaith-Hamilton Pleasure Scale; TEPS=Temporal Experience of Pleasure Scale; PSS=Perceived Stress Scale; QLESQ=Quality of Life Enjoyment and Satisfaction Questionnaire.

### Dynamic Approach-Avoidance Conflict (AAC) Task

Decision-making was assessed using a dynamic approach-avoidance conflict task back-translated from non-human primate research^14,26^. For full details see Supplementary Information and **Supplementary Fig. 2**. In brief, on each trial, participants decided whether to approach or avoid an offer defined by a monetary reward component and an aversive electrical shock component, each varying in magnitude and displayed as horizontal bars. Offers were dynamically adjusted on a trial-by-trial basis to track each participant’s individual indifference point between approach and avoidance across the reward-aversion space, ensuring individually calibrated levels of conflict throughout the task. Participants indicated their choices using a joystick. On Pavlovian response-congruent trials, approach was indicated by pulling the joystick toward themselves, whereas on Pavlovian response-incongruent trials, approach was indicated by pushing the joystick away. These trial types were counterbalanced across the session. Following each choice, participants received feedback on earnings and, if they approached, the corresponding shock was administered. A total of 105 offers were presented per run across two fMRI runs completed before and after stress induction, providing sufficient trials to model individual differences in approach-avoidance decision-making under each condition.

### Stress Induction Procedure

Acute stress was induced using the Maastricht Acute Stress Test (MAST)^35^, a well-validated paradigm that combines a physiological and a psychosocial stressor. We have previously implemented the MAST for use in the MRI in prior independent MDD samples^7,19^. Participants alternated between hand immersion in ice-cold water (0-4°C) and performing a mental arithmetic test, during which they were monitored on their performance by two evaluators providing negative feedback in case of mistakes. The MAST reliably elicits autonomic and endocrine stress responses, which were confirmed in this study through salivary cortisol sampling at multiple time points. To sustain the stress induction throughout the post-stress task, participants were told that their performance had been below average and that they would have to repeat the stressor later in the session. They were debriefed after leaving the scanner. The post-stress Ap-Av task began immediately after the MAST and spanned approximately 15 to 45 minutes after stress onset, thus covering the ascending phase of the cortisol response up to its peak (**Supplementary Fig. 1**).

### fMRI Acquisition and Preprocessing

For details on fMRI data acquisition and preprocessing, see the “Functional magnetic resonance imaging (fMRI)” section in the Supplementary Information. To extract single-trial neural activation estimates, we applied the Least Squares Separate (LSS) approach^41^. This method improves the estimation of trial-specific activation by modeling each trial separately while accounting for collinearity with surrounding events. The resulting single-trial beta estimates were then used as input for our hierarchical Bayesian computational model.

### Computational Modeling Approach

A hierarchical Bayesian drift-diffusion model (HDDM) in which the decision threshold collapses linearly over the course of a trial was used to dissect approach-avoidance decision-making. The computational analysis was conducted in two stages. The first stage fitted a behavioral-only model, without neural regressors, to select the model and to establish whether the characterization of the decision process depended on the inclusion of trial-wise neural signals. Bayesian model comparison between a standard drift-diffusion model and the Angle model, both fitted to this dataset, identified the Angle model as the better fit, replicating the model selection reported for this task in an independent sample^26^ (see Behavioral-Only Model Results section of the Supplementary Information). Selection was based primarily on the deviance information criterion, evaluated together with posterior predictive checks in both groups. The Angle model is a widely used modification of the HDDM in which the rate of collapse is captured by a single parameter^42–45^. As such, model parameters included *drift rate* (v), representing the rate of information accumulation and indexing reward and aversion sensitivity; *decision threshold* (a; also termed boundary separation), indicating response caution and speed-accuracy trade-offs; *rate of threshold collapse* (θ), quantifying decision urgency; *starting point bias* (z), reflecting pre-decisional biases based on prior experiences; and *nondecision time* (Ter), measuring the time required for stimulus encoding and motor execution. In addition, this behavioral model accounts for the influence of the varying levels of offered reward, aversion, and conflict present on each trial^26^. These task variables were z-scored within each subject and condition before being incorporated as regressors in the model to estimate effects on drift and the rate of threshold collapse.

In the second stage, neural measures were integrated into the best-fitting computational model to assess the contribution of brain activity to approach-avoidance decision-making under stress, using a model that combined both stress conditions in order to capture stress-induced changes in neural modulation. Each model was fitted separately to the HC and rMDD groups. Because both conditions were modeled jointly, the post-stress condition served as the reference. Effects of the stressor are therefore estimated as the difference between conditions, and pre-stress estimates were derived from the same posteriors rather than from a separately fitted pre-stress model. Functional MRI data from two regions of interest (ROIs), a priori selected based on prior literature, were used as regressors in the hierarchical Bayesian model: pgACC, which contributes to the valuation of offers and to decision-making under conflict, and the STN, which regulates inhibitory control and response cautiousness. The pgACC was defined as described previously^17^, using a sphere centered on meta-analysis coordinates^46^. The STN was defined bilaterally from left and right middle-aged probabilistic maps^47^. The caudate nucleus and nucleus accumbens were additionally considered in earlier model iterations, given their roles in reinforcement learning and motivational bias, but were not retained because removing them improved model fit as indexed by the DIC. Mean single-trial LSS betas were extracted from each mask; ROI activity was then z-scored within each subject and condition and included as regressors to estimate neural contributions to drift rate, decision threshold, and the rate of threshold collapse across conditions.

Including trial-wise neural signals as predictors of decision parameters allows each region to be linked to a specific computational process, beyond what is explained by the offered reward and aversion alone^27^. Each region was assigned a priori to specific decision parameters on the basis of prior work rather than selected by model fit, and neural regressors were entered at the level of individual trials after within-subject standardization, so the resulting coupling estimates reflect trial-to-trial covariation within participants rather than group differences in mean activation. Accordingly, trial-wise drift rate was modeled as a function of offered reward and aversion, of pgACC activity, and of the interactions between them, with all terms additionally crossed with domain. The decision threshold was modeled as a function of STN activity, and the rate of threshold collapse as a function of STN activity, trial-wise conflict, and their interaction. Starting point bias was modeled as a function of Pavlovian response congruency. All terms were further allowed to vary by stress condition, so that each neural effect could differ before and after the stressor. Formally:

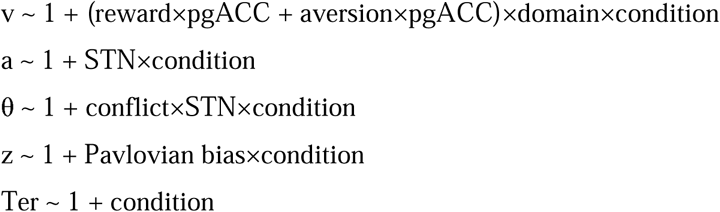

The reported neural model excludes interaction terms of fourth order and above. A comparison with the full model, including all higher-order interactions, is provided in the Supplementary Information (**Supplementary Figs. 10-15** and **Supplementary Table 4**).

### Statistical Analysis

#### Stress Response and Self-Report Measures

Physiological and subjective responses to the MAST were analyzed with repeated-measures analyses of variance including Group as a between-subjects factor and Time as a within-subjects factor. Salivary cortisol values were log-transformed prior to analysis. One missing cortisol value, representing 0.16% of all cortisol data points, was imputed using the mice package in R^48^ with *Group* and *Timepoint* as predictors. Where the assumption of sphericity was violated, degrees of freedom were adjusted using the Greenhouse-Geisser correction, and effect sizes are reported as partial eta squared. Visual Analogue Mood Scale ratings were analyzed with a repeated-measures multivariate analysis of variance across the three affective dimensions, followed by univariate tests. Interactions were followed up with pairwise comparisons corrected for multiple comparisons. Group differences in demographic and clinical characteristics were tested with two-sample *t* tests for continuous variables and chi-square or Fisher exact tests for categorical variables. Missing questionnaire and interview responses were likewise imputed at the item level within each instrument, using that instrument’s remaining items together with group and visit as predictors, following the procedure used previously in this cohort^49^. Missingness was minimal, affecting 59 of 40,543 item responses (0.15%). Assessments that were not attended were not imputed, so participants contributed every visit at which they were assessed. Full statistics are reported in the Supplementary Information.

#### Sequential Sampling Modeling

Hierarchical Bayesian models were estimated using Markov Chain Monte Carlo (MCMC) sampling, with three chains per model. Each chain drew 60,000 iterations, and a burn-in of 40,000 iterations. The full neural model in the rMDD group required 80,000 iterations per chain, with the same burn-in. Model convergence was assessed using Gelman-Rubin statistics, with R-hat values below 1.1 indicating convergence. Group differences were evaluated using the posterior distribution of the difference between the two groups’ estimates, computed from their separately sampled posteriors.. The posterior probability (PP) that a parameter was larger in rMDD than in HC, PP[rMDD>HC], was defined as the proportion of this posterior distribution lying above zero. Because this quantity is directional, values approaching 1 indicate a credibly larger estimate in rMDD and values approaching 0 a credibly larger estimate in HC. Our criterion for a credible group difference was PP≥0.95 or PP≤0.05. Within-group effects are reported as posterior means with 95% highest density intervals (HDI) and were considered credible when the HDI excluded zero.

### Prospective Analyses

Clinician-rated depressive symptoms on the HAMD^50^ were the primary prospective outcome, with self-reported depressive symptoms on the Beck Depression Inventory-II (BDI-II)^51^ and the 16-item Quick Inventory of Depressive Symptomatology, Self-Report (QIDS-SR16)^52^, anhedonia on the Snaith-Hamilton Pleasure Scale (SHAPS)^53^, perceived stress on the 14-item Perceived Stress Scale (PSS-14)^54^ and quality of life on the Quality of Life Enjoyment and Satisfaction Questionnaire, Short Form (Q-LES-Q-SF)^55^ examined as specificity checks. Anticipatory and consummatory pleasure were assessed with the Temporal Experience of Pleasure Scale (TEPS)^56^ for sample characterization only (**Table 2**).

The QIDS-SR16 was scored with the published nine-domain algorithm, in which sleep, appetite and weight change, and psychomotor disturbance each contribute the maximum of their constituent items (range 0–27). The predictor was the stress-induced change in the influence of reward magnitude on drift rate for net-aversive offers, taken as the subject-level posterior mean from the reported neural model and standardized across the whole sample. Symptom scores from all three assessments were modeled with generalized estimating equations assuming a Gaussian distribution, an exchangeable working correlation and robust sandwich standard errors, entering the predictor, time in months, and their interaction. Group was modeled explicitly rather than assuming homogeneity of regression slopes, so that the three-way interaction of predictor, time and group tests whether the association differs between rMDD and HC. Two confirmatory analyses are reported, one refitting the same model with visit entered as a factor and one analyzing each follow-up visit separately with baseline symptoms as a covariate. Analyses were conducted in R (version 4.5.2) using the geepack package (version 1.3.13), with imputation performed using mice (version 3.19.0) as described above.

## Declaration of Interest

Over the past three years, Dr. Pizzagalli has received consulting fees from Abbvie, Boehringer Ingelheim, Circular Genomics, Compass Pathways, Engrail Therapeutics, N1 Bio Corp., Neumora Therapeutics, Neurocrine Biosciences, Neuroscience Software, Syntropic Medical, Tap Sciences, and Xenon Pharmaceuticals; he has received honoraria from the American Psychological Association, Psychonomic Society and Springer (for editorial work) and from Alkermes; he has received research funding from the Brain and Behavior Research Foundation, Circular Genomics, Dana Foundation, Millennium Pharmaceuticals, NIMH, and Wellcome Leap; he has received stock options from Ceretype Neuromedicine, Compass Pathways, Engrail Therapeutics, GLX Analytix, Neumora Therapeutics, Neuroscience Software, and ReWire Neurotechnologies, Inc. No funding from these entities was used to support the current work, and all views expressed are solely those of the authors. The other authors declare no competing financial interests.

## Funding

The present work was supported by the National Institute of Mental Health grants P50 MH119467 and R37 MH068376 to Diego A. Pizzagalli. The funders had no role in study design, data collection and analysis.

## References

1. Bulloch, A., Williams, J., Lavorato, D. & Patten, S. Recurrence of major depressive episodes is strongly dependent on the number of previous episodes. Depress Anxiety 31, 72–76 (2014).

2. Breedvelt, J. J. F. et al. An individual participant data meta-analysis of psychological interventions for preventing depression relapse. Nat. Mental Health 2, 154–163 (2024).

3. LeMoult, J. et al. Advances in stress and depression research. Curr Opin Psychiatry 36, 8–13 (2023).

4. Segal, Z. V. et al. Cognitive Reactivity to Sad Mood Provocation and the Prediction of Depressive Relapse. Arch Gen Psychiatry 63, 749 (2006).

5. Kuhn, M. et al. Psychobiological Stress Response Profiles in Current and Remitted Depression: A Person-Centered, Multisystem Approach. Biological Psychiatry Global Open Science 5, 100400 (2025).

6. Admon, R. et al. Striatal Hypersensitivity During Stress in Remitted Individuals with Recurrent Depression. Biological Psychiatry 78, 67–76 (2015).

7. Whitton, A. E. et al. Acute stress impairs frontocingulate activation during error monitoring in remitted depression. Psychoneuroendocrinology 75, 164–172 (2017).

8. Ming, Q. et al. State-Independent and Dependent Neural Responses to Psychosocial Stress in Current and Remitted Depression. AJP 174, 971–979 (2017).

9. Eshel, N. & Roiser, J. P. Reward and Punishment Processing in Depression. Biological Psychiatry 68, 118–124 (2010).

10. Treadway, M. T. & Zald, D. H. Reconsidering anhedonia in depression: Lessons from translational neuroscience. Neuroscience & Biobehavioral Reviews 35, 537–555 (2011).

11. Aupperle, R. L., Melrose, A. J., Francisco, A., Paulus, M. P. & Stein, M. B. Neural substrates of approach-avoidance conflict decision-making. Hum Brain Mapp 36, 449–462 (2015).

12. Aupperle, R. L. & Paulus, M. P. Neural systems underlying approach and avoidance in anxiety disorders. Dialogues in Clinical Neuroscience 12, 517–531 (2010).

13. Bartra, O., McGuire, J. T. & Kable, J. W. The valuation system: A coordinate-based meta-analysis of BOLD fMRI experiments examining neural correlates of subjective value. NeuroImage 76, 412–427 (2013).

14. Amemori, K. & Graybiel, A. M. Localized microstimulation of primate pregenual cingulate cortex induces negative decision-making. Nat Neurosci 15, 776–785 (2012).

15. Papageorgiou, G. K. et al. Functional distinctions between orbitofrontal cortex and anterior cingulate cortex subregions in decision-making and autonomic regulation. Nat Commun 10.1038/s41467-026-69447-4 (2026) doi:10.1038/s41467-026-69447-4.

16. Friedman, A. et al. A Corticostriatal Path Targeting Striosomes Controls Decision-Making under Conflict. Cell 161, 1320–1333 (2015).

17. Ironside, M. et al. Approach-Avoidance Conflict in Major Depressive Disorder: Congruent Neural Findings in Humans and Nonhuman Primates. Biological Psychiatry 87, 399–408 (2020).

18. Dong, D. et al. Sex-specific neural responses to acute psychosocial stress in depression. Transl Psychiatry 12, 2 (2022).

19. Ironside, M. et al. Association of Lower Rostral Anterior Cingulate GABA+ and Dysregulated Cortisol Stress Response With Altered Functional Connectivity in Young Adults With Lifetime Depression: A Multimodal Imaging Investigation of Trait and State Effects. AJP 181, 639–650 (2024).

20. Bogdan, R. & Pizzagalli, D. A. Acute stress reduces reward responsiveness: implications for depression. Biol Psychiatry 60, 1147–1154 (2006).

21. Duda, J. M. et al. Effects of GABA, Sex, and Stress on Reward Learning in Current and Remitted Major Depression. Biol Psychiatry Cogn Neurosci Neuroimaging 9, 606–615 (2024).

22. Pizzagalli, D. A. Probing biomarkers and clinical utility of reward learning across species using the Probabilistic Reward Task: 20 years of findings. Nat. Mental Health 4, 1066–1087 (2026).

23. Pechtel, P., Dutra, S. J., Goetz, E. L. & Pizzagalli, D. A. Blunted reward responsiveness in remitted depression. J Psychiatr Res 47, 1864–1869 (2013).

24. Whitton, A. E. et al. Blunted Neural Responses to Reward in Remitted Major Depression: A High-Density Event-Related Potential Study. Biological Psychiatry: Cognitive Neuroscience and Neuroimaging 1, 87–95 (2016).

25. Ratcliff, R. & McKoon, G. The diffusion decision model: theory and data for two-choice decision tasks. Neural Comput 20, 873–922 (2008).

26. Ging-Jehli, N. R. et al. Cognitive Signatures of Depressive and Anhedonic Symptoms and Affective States Using Computational Modeling and Neurocognitive Testing. Biological Psychiatry: Cognitive Neuroscience and Neuroimaging 9, 726–736 (2024).

27. Pedersen, M. L. et al. Computational phenotyping of brain-behavior dynamics underlying approach-avoidance conflict in major depressive disorder. PLoS Comput Biol 17, e1008955 (2021).

28. Cavanagh, J. F. et al. Subthalamic nucleus stimulation reverses mediofrontal influence over decision threshold. Nat Neurosci 14, 1462–1467 (2011).

29. Frank, M. J. et al. fMRI and EEG predictors of dynamic decision parameters during human reinforcement learning. J Neurosci 35, 485–494 (2015).

30. Ging-Jehli, N. R. et al. Basal ganglia components have distinct computational roles in decision-making dynamics under conflict and uncertainty. PLoS Biol 23, e3002978 (2025).

31. Herz, D. M., Zavala, B. A., Bogacz, R. & Brown, P. Neural Correlates of Decision Thresholds in the Human Subthalamic Nucleus. Current Biology 26, 916–920 (2016).

32. Cole, R. C. et al. Theta-frequency subthalamic nucleus stimulation increases decision threshold. Brain Stimulation 18, 1021–1027 (2025).

33. Green, N. et al. Reduction of Influence of Task Difficulty on Perceptual Decision Making by STN Deep Brain Stimulation. Current Biology 23, 1681–1684 (2013).

34. Herz, D. M., Frank, M. J., Tan, H. & Groppa, S. Subthalamic control of impulsive actions: insights from deep brain stimulation in Parkinson’s disease. Brain 147, 3651–3664 (2024).

35. Smeets, T. et al. Introducing the Maastricht Acute Stress Test (MAST): a quick and non-invasive approach to elicit robust autonomic and glucocorticoid stress responses. Psychoneuroendocrinology 37, 1998–2008 (2012).

36. Ironside, M., Kumar, P., Kang, M.-S. & Pizzagalli, D. A. Brain mechanisms mediating effects of stress on reward sensitivity. Curr Opin Behav Sci 22, 106–113 (2018).

37. Lighthall, N. R., Gorlick, M. A., Schoeke, A., Frank, M. J. & Mather, M. Stress modulates reinforcement learning in younger and older adults. Psychology and Aging 28, 35–46 (2013).

38. Porcelli, A. J. & Delgado, M. R. Stress and decision making: effects on valuation, learning, and risk-taking. Current Opinion in Behavioral Sciences 14, 33–39 (2017).

39. Kuhn, M. et al. Computational Phenotyping of Effort-Based Decision Making in Unmedicated Adults With Remitted Depression. Biological Psychiatry: Cognitive Neuroscience and Neuroimaging 10.1016/j.bpsc.2025.02.006 (2025) doi:10.1016/j.bpsc.2025.02.006.

40. De Hollander, G., Keuken, M. C., Van Der Zwaag, W., Forstmann, B. U. & Trampel, R. Comparing functional MRI protocols for small, iron-rich basal ganglia nuclei such as the subthalamic nucleus at 7 T and 3 T. Human Brain Mapping 38, 3226–3248 (2017).

41. Mumford, J. A., Turner, B. O., Ashby, F. G. & Poldrack, R. A. Deconvolving BOLD activation in event-related designs for multivoxel pattern classification analyses. Neuroimage 59, 2636–2643 (2012).

42. Hawkins, G. E., Forstmann, B. U., Wagenmakers, E.-J., Ratcliff, R. & Brown, S. D. Revisiting the Evidence for Collapsing Boundaries and Urgency Signals in Perceptual Decision-Making. J Neurosci 35, 2476–2484 (2015).

43. Churchland, A. K., Kiani, R. & Shadlen, M. N. Decision-making with multiple alternatives. Nat Neurosci 11, 693–702 (2008).

44. Fengler, A., Govindarajan, L. N., Chen, T. & Frank, M. J. Likelihood approximation networks (LANs) for fast inference of simulation models in cognitive neuroscience. eLife 10, e65074 (2021).

45. Hanks, T. D., Mazurek, M. E., Kiani, R., Hopp, E. & Shadlen, M. N. Elapsed Decision Time Affects the Weighting of Prior Probability in a Perceptual Decision Task. J. Neurosci. 31, 6339–6352 (2011).

46. Marusak, H. A. et al. You say ‘prefrontal cortex’ and I say ‘anterior cingulate’: meta-analysis of spatial overlap in amygdala-to-prefrontal connectivity and internalizing symptomology. Transl Psychiatry 6, e944–e944 (2016).

47. Keuken, M. C. et al. Ultra-High 7T MRI of Structural Age-Related Changes of the Subthalamic Nucleus. J. Neurosci. 33, 4896–4900 (2013).

48. Buuren, S. van & Groothuis-Oudshoorn, K. mice: Multivariate Imputation by Chained Equations in R. Journal of Statistical Software 45, 1–67 (2011).

49. Palermo, E. H. et al. Characterizing emotion dynamics in remitted depression: A network approach using ecological momentary assessment. Journal of Mood & Anxiety Disorders 14, 100182 (2026).

50. Hamilton, M. A rating scale for depression. J Neurol Neurosurg Psychiatry 23, 56–62 (1960).

51. Beck, A. T., Steer, R. A. & Brown, G. Beck Depression Inventory–II. (American Psychological Association, 2011). doi:10.1037/t00742-000.

52. Rush, A. J. et al. The 16-Item quick inventory of depressive symptomatology (QIDS), clinician rating (QIDS-C), and self-report (QIDS-SR): a psychometric evaluation in patients with chronic major depression. Biological Psychiatry 54, 573–583 (2003).

53. Snaith, R. P. et al. A scale for the assessment of hedonic tone the Snaith-Hamilton Pleasure Scale. Br J Psychiatry 167, 99–103 (1995).

54. Cohen, S., Kamarck, T. & Mermelstein, R. A global measure of perceived stress. Journal of Health and Social Behavior 24, 385–396 (1983).

55. Endicott, J., Nee, J., Harrison, W. & Blumenthal, R. Quality of Life Enjoyment and Satisfaction Questionnaire: a new measure. Psychopharmacol Bull 29, 321–326 (1993).

56. Gard, D. E., Gard, M. G., Kring, A. M. & John, O. P. Anticipatory and consummatory components of the experience of pleasure: A scale development study. Journal of Research in Personality 40, 1086–1102 (2006).

