## Supplement for "Acute stress reorganizes pregenual cingulate and subthalamic coupling to approach-avoidance decisions in remitted depression"

Manuel Kuhn et al.

**Supplementary Methods**

**Experimental procedures**

The experimental fMRI session was performed at the McLean Imaging Center, Belmont, MA, USA. For a graphical representation of the session procedure, see **Supplementary Fig. 1**. In brief, the session included: participant check-in and preparation; a shock calibration procedure (see procedure description below), a functional magnetic resonance (fMRI) scan including clinical sequences that were independently reviewed by a board-certified radiologist to screen for incidental findings, structural MRI scan, two runs of the functional MRI approach-avoidance conflict task (dynamic AAC task pre- and post-stress, see below) as well as an acute stress-induction via the Maastricht Acute Stress Test (MAST)^1^; two offline computerized tasks (Probabilistic Rewards Task – PRT^2^ , Effort Expenditure for Rewards Task – EEfRT^3^), debrief with EMA setup and participant checkout. Throughout the experiment, salivary cortisol and affective self-report ratings were administered, including Visual Analogue Mood Scales (VAMS), the Positive and Negative Affect Schedule (PANAS)^4^, and the Threat–Challenge Questionnaire (TCQ)^5^. Notably, the MAST procedure was modified to induce prolonged stress during the post-MAST dynamic AAC task administration, as accomplished in prior independent MDD and rMDD samples (e.g., ^6–8^). Specifically, to sustain stress during the post-MAST run of the dynamic AAC, participants were told after the MAST procedure, regardless of their actual performance, that their performance had been below average and that they would have to repeat the task later in the experiment. After the task, participants were told that they would not have to repeat it, providing “stress relief”.

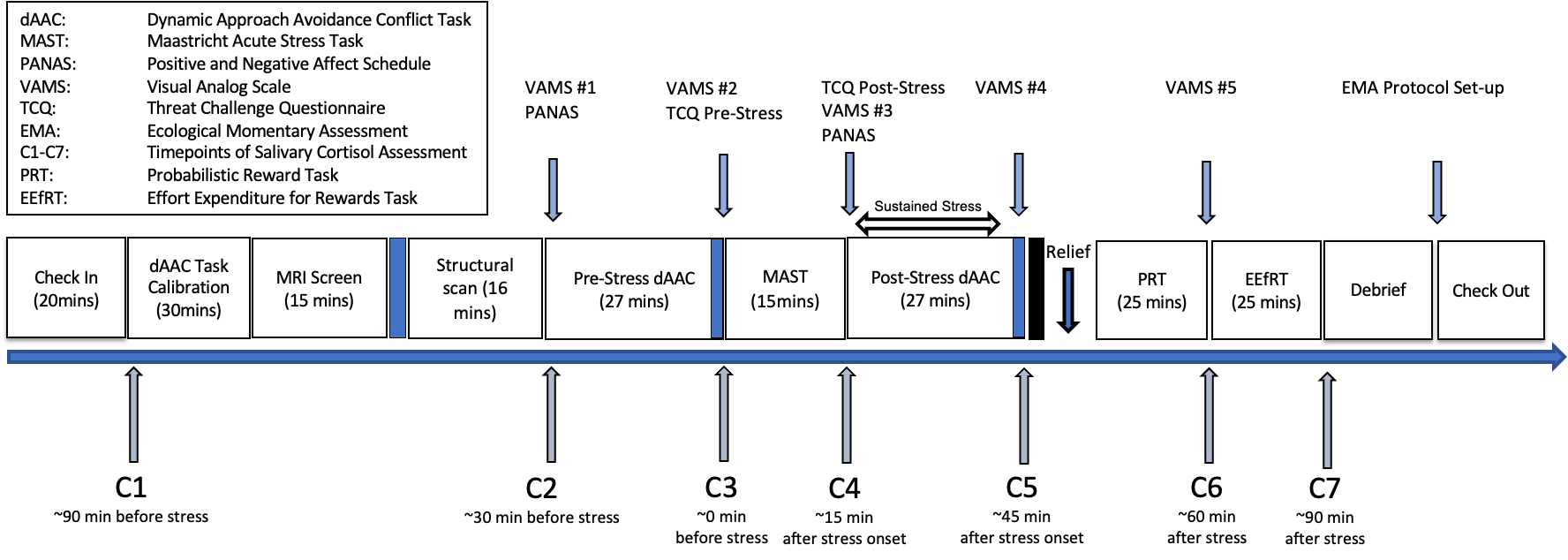

**Supplementary Fig. 1**. Schematic overview of the scan session study procedure.

**Dynamic Approach-Avoidance Conflict (AAC) Task Procedure**

*Preparation*

After the participant provided consent, the experimenter attached two rectangular surface electrodes (Coulburn Instruments, Holliston, MA, USA) to the participant’s right ankle. A calibration procedure (for details, see below) determined 10 stimulus intensities that induced shock levels from a slightly uncomfortable feeling that would still affect decision-making in the task (shock level 1) to a highly aversive but not painful feeling (shock level 10). After calibration, participants received task instructions displayed on a computer screen.

*Shock calibration*

The shock calibration was performed in a mock scanner environment to allow the participant to adjust to the MR environment. The calibration followed the procedure of our previous work^9^. In detail, the electrical stimulation was generated by a Digitimer DG2A Train/Delay generator (Digitimer, Welwyn Garden City, UK) that generated pulse trains of 500ms at 50Hz and delivered shocks at this frequency via a DS8R (Digitimer DS8R Biphasic Constant Current Stimulator, Digitimer, Welwyn Garden City, UK). The two electrodes attached to the participant’s right ankle were used for stimulation. At the beginning of the shock level calibration procedure, participants were instructed that the procedure determined their individual level 1 stimulation (minimum) and level 10 stimulation (maximum). Participants were told that level 1 should be slightly uncomfortable (like a light pinch) but still carry some weight in their decision-making and that level 10 should be maximally uncomfortable without being painful. The calibration procedure started by applying a 1mA stimulation and then proceeding in steps of 1mA (or smaller increments if necessary) until the participant expressed that the stimulation was slightly uncomfortable. At this point, participants were asked how they would feel about taking the slightly uncomfortable stimulation for 5 cents, to determine whether this level of stimulation carried some weight in their decision-making. If they reported that they would take this offer (stimulation level in question for 5 cents) 40% to 60% of the time (demonstrating that this stimulation carried some weight) this level was chosen as level 1. Once the level 1 was determined, stimulation was gradually increased in intensity to determine the level 10. The stimulation was increased in intensity until the participant reported that the stimulation resulted in their maximum level of discomfort (that they would be willing to take during the task) without being painful. At this point, they were asked how they would feel about taking the highly uncomfortable stimulation for additional 5 to 50 cents. If they reported that they would take this offer 40% to 60% of the time (demonstrating that this carried some weight) this level was chosen as the level 10. In cases where participants offered responses that seemed inconsistent (such as agreeing to endure greater electric shocks for smaller monetary rewards than for larger shocks) or indicated changes in their willingness to accept a particular shock level for a given monetary incentive, the calibration process was either repeated or necessary adjustments to individual shock levels were made. Next, participants completed 15 practice trials of the task while supervised by the experimenter, allowing for readjustment of the shock level calibration in case participants should, for example, choose approach on all practice trials.

*Dynamic AAC task*

The task was identical to the task used in our previous work^9^. For a graphical representation of the trial structure see **Supplementary Fig. 2A.** The task was implemented in Psychtoolbox running on MATLAB 2020a. Shock intensity was controlled via MATLAB on the DS8R stimulation device, which was triggered via MATLAB and a NI DAQ USB-6009 device (National Instruments, Austin, TX, USA) which triggered the DG2A pulse train generator. Participants used a MR-compatible joystick (NATA Technologies, Coquitlam, Canada) to indicate their responses. Specifically, on Pavlovian response congruent trials, approach decisions were indicated by pulling the joystick. On Pavlovian response incongruent trials, approach decisions were indicated by pushing away the joystick (**Supplementary Fig. 2B)**. Pavlovian response congruent and incongruent trials were counterbalanced. For each presented offer, we collected participants’ choice and RTs that served as inputs into our computational models. The dynamic AAC is a newly developed task based on our prior work in humans and non-human primates^10^. The task was optimized to induce individually calibrated levels for aversion and used a trial-by-trial dynamic (i.e., adaptive) procedure to optimally induce variable levels of approach-avoidance conflict. Moreover, ecologically valid stimuli (i.e., physically aversive stimulation and monetary rewards) were introduced to generate optimal individually adjusted degrees of conflict. As such, the dynamic AAC task’s design allows to induce approach-avoidance conflict on a reward-aversion continuum with varying degrees of conflict, while adjusting for individual decision indifference points within this continuum. Offers were dynamically created on a trial-by-trial case and for each participant separately. Specifically, the first nine trials were pseudo-randomly sampled on a 3x3 grid from the reward-aversion space (i.e., rewards and aversion could each range from 10 to 100, in steps of 10 with one trial selected from each subspace within the 3x3 grid). Afterwards, for each subsequent trial and for each participant their momentary indifference curve along which subjects are equally likely to approach or avoid were estimated. To do so, logistic regression with input of all prior choices (up to and excluding the current trial) as dependent variable (1=approach, 0=avoid) and the corresponding reward and aversion as independent variables was used. Offers were sampled from normal distributions centered on the estimated indifference curve, denoted 𝒩(0, σ) where σ reflects the spread of offers around the curve, with 40% probability drawn from close to the curve (𝒩(0, 5%)), 30% from farther away (𝒩(0, 20%)), and 30% drawn pseudo-randomly from subspaces within a 3×3 grid spanning the reward–aversion space, while ensuring balanced sampling across grid tiles. Response symbols and offers were simultaneously presented for 4 seconds. When participants made a choice, the circle was locked at the target until the 4s were up. A fixation cross was then presented and if participants decided to approach an offer, they were administered the corresponding shock level for 500ms during this period (jittered after fixation cross onset, range 0.5-10s; mean 1.77s). If participants decided to avoid or did not respond within the time window, no shock was delivered. Following, participants received a 2s feedback about the number of cents they earned in this trial (0 in case of avoid or no-response).

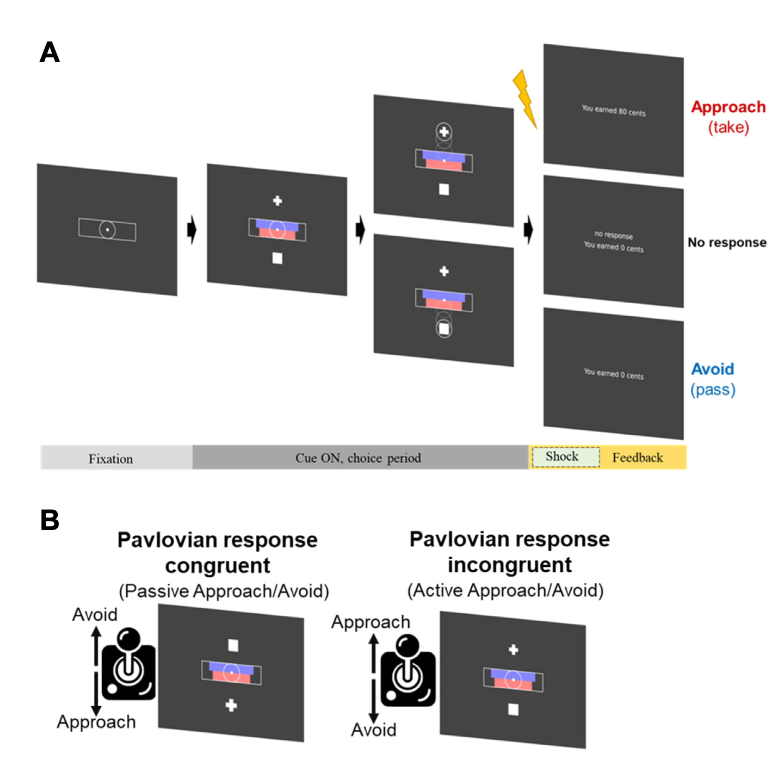

**Supplementary Fig. 2**. Approach-avoidance conflict task. (A) Participants decided to approach or avoid offers with monetary reward (magnitude represented by blue bars) and electrical shock (magnitude represented by red bars). (B) Trials were counterbalanced including either Pavlovian response-congruent trials or Pavlovian response-incongruent trials. Reproduced from ^9^, with permission from the publisher.

**Functional magnetic resonance imaging (fMRI)**

*fMRI acquisition*

MRI data were acquired on a 3T Siemens Prisma scanner (Siemens Medical Systems, Iselin, NJ) equipped with a 64-channel head coil. T1-weighted anatomical images were acquired using a 3D magnetization-prepared rapid gradient-echo (MPRAGE) sequence (TR = 2530 ms; TE = 1.69, 3.55, 5.41, 7.27 ms; field of view = 256 mm; voxel size = 1.0 × 1.0 × 1.0 mm³; 176 slices; GRAPPA acceleration factor = 2). Functional images were acquired using a T2*-weighted gradient-echo echo-planar imaging (EPI) sequence (TR = 800 ms; TE = 37 ms; field of view = 208 mm; voxel size = 2.0 × 2.0 × 2.0 mm³; 72 interleaved slices; multiband factor = 8).

*fMRI preprocessing*

Preprocessing was performed using fMRIPrep 23.1.4 ^11,12^, which is based on Nipype 1.8.6 ^13^. The following preprocessing procedure was applied:

*Preprocessing of B0 inhomogeneity mappings:* For each participant, available EPI fieldmaps in the BIDS dataset were used to estimate a B0-nonuniformity map (fieldmap) with topup ^14^.

*Anatomical data preprocessing:* The T1-weighted (T1w) image was corrected for intensity non-uniformity (INU) with N4BiasFieldCorrection ^15^, distributed with ANTs ^16^, and used as T1w-reference throughout the workflow. The T1w-reference was then skull-stripped with a Nipype implementation of the antsBrainExtraction.sh workflow (from ANTs), using OASIS30ANTs as target template. Brain tissue segmentation of cerebrospinal fluid (CSF), white-matter (WM) and gray-matter (GM) was performed on the brain-extracted T1w using fast (FSL, ^17^). Brain surfaces were reconstructed using recon-all (FreeSurfer 7.3.2, ^18^), and the brain mask estimated previously was refined with a custom variation of the method to reconcile ANTs-derived and FreeSurfer-derived segmentations of the cortical gray-matter of Mindboggle ^19^. Volume-based spatial normalization to two standard spaces (MNI152NLin6Asym, MNI152NLin2009cAsym) was performed through nonlinear registration with antsRegistration (ANTs), using brain-extracted versions of both T1w reference and the T1w template. The following templates were selected for spatial normalization and accessed with TemplateFlow (23.0.0, ^20^): FSL’s MNI ICBM 152 non-linear 6th Generation Asymmetric Average Brain Stereotaxic Registration Model [^21^, TemplateFlow ID: MNI152NLin6Asym], ICBM 152 Nonlinear Asymmetrical template version 2009c [^22^, TemplateFlow ID: MNI152NLin2009cAsym].

*Functional data preprocessing:* For each BOLD run a reference volume and its skull-stripped version were generated by aligning and averaging single-band references (SBRefs). Head-motion parameters with respect to the BOLD reference (transformation matrices, and six corresponding rotation and translation parameters) are estimated before any spatiotemporal filtering using mcflirt (FSL, ^23^). The estimated fieldmap was then aligned with rigid-registration to the target EPI (echo-planar imaging) reference run. The field coefficients were mapped on to the reference EPI using the transform. BOLD runs were slice-time corrected to 0.346s (0.5 of slice acquisition range 0s-0.693s) using 3dTshift from AFNI ^24^. The BOLD reference was then co-registered to the T1w reference using bbregister (FreeSurfer) which implements boundary-based registration^25^. Co-registration was configured with six degrees of freedom. Several confounding time-series were calculated based on the preprocessed BOLD: framewise displacement (FD), DVARS and three region-wise global signals. FD was computed using two formulations following Power (absolute sum of relative motions, ^26^) and Jenkinson (relative root mean square displacement between affines, ^23^). FD and DVARS are calculated for each functional run, both using their implementations in Nipype (following the definitions by ^26^). The three global signals were extracted within the CSF, the WM, and the whole-brain masks. Additionally, a set of physiological regressors were extracted to allow for component-based noise correction (CompCor, ^27^). Principal components were estimated after high-pass filtering the preprocessed BOLD time-series (using a discrete cosine filter with 128s cut-off) for the two CompCor variants: temporal (tCompCor) and anatomical (aCompCor). tCompCor components are then calculated from the top 2% variable voxels within the brain mask. For aCompCor, three probabilistic masks (CSF, WM and combined CSF+WM) are generated in anatomical space. The implementation differs from that of Behzadi et al. in that instead of eroding the masks by 2 pixels on BOLD space, a mask of pixels that likely contain a volume fraction of GM is subtracted from the aCompCor masks. This mask is obtained by dilating a GM mask extracted from the FreeSurfer’s aseg segmentation, and it ensures components are not extracted from voxels containing a minimal fraction of GM. Finally, these masks are resampled into BOLD space and binarized by thresholding at 0.99 (as in the original implementation). Components are also calculated separately within the WM and CSF masks. For each CompCor decomposition, the k components with the largest singular values are retained, such that the retained components’ time series are sufficient to explain 50 percent of variance across the nuisance mask (CSF, WM, combined, or temporal). The remaining components are dropped from consideration. The head-motion estimates calculated in the correction step were also placed within the corresponding confounds file. The confound time series derived from head motion estimates and global signals were expanded with the inclusion of temporal derivatives and quadratic terms for each ^28^. Frames that exceeded a threshold of 0.5 mm FD or 1.5 standardized DVARS were annotated as motion outliers. Additional nuisance timeseries were calculated by means of principal components analysis of the signal found within a thin band (crown) of voxels around the edge of the brain, as proposed by ^29^. The BOLD time-series were resampled into several standard spaces, correspondingly generating the following spatially-normalized, preprocessed BOLD runs: MNI152NLin6Asym, MNI152NLin2009cAsym. The BOLD time-series were resampled onto the following surfaces (FreeSurfer reconstruction nomenclature): fsaverage. All resamplings can be performed with a single interpolation step by composing all the pertinent transformations (i.e. head-motion transform matrices, susceptibility distortion correction when available, and co-registrations to anatomical and output spaces). Gridded (volumetric) resamplings were performed using antsApplyTransforms (ANTs), configured with Lanczos interpolation to minimize the smoothing effects of other kernels ^30^. Non-gridded (surface) resamplings were performed using mri_vol2surf (FreeSurfer). Subsequently, data spatially smoothed with a 6-mm full-width-at-half-maximum Gaussian kernel.

**Cortisol Assessment**

As a physiological marker of stress response, salivary cortisol was assessed at seven time points relative to the onset of the stress induction via the MAST (approximate time with respect to stress onset, C1 -90 min, C2 -30 min, C3 0 min / pre-stress baseline, C4 +15 min, C5 +45 min, C6 +60 min, C7 +90 min, see **Supplementary** **Fig. 1** for assessment timepoints). Consistent with prior work ^31^, the first two sampling points were excluded from analyses because they occurred well before the stress induction and may be more influenced by individual variation in pre-experimental factors (e.g., food intake, physical activity). The third time point C3, collected immediately before stress onset, was considered the optimal pre-stress baseline, while subsequent samples C4-C7 captured the acute stress response and recovery. Salivary cortisol was collected using Salivettes (Salivette®, Sarstedt, Nümbrecht, Germany), with participants instructed to place the cotton swabs in their mouth for approximately 2 minutes. Samples were stored at -80 °C until analysis, and assays of free cortisol were performed using a commercially available enzyme-linked immunosorbent assay (ELISA). Assay quality control across three batches indicated an average intra-assay coefficient of variation of 3.3% and an average inter-assay coefficient of variation of 5.2%, demonstrating high reliability of the cortisol measurements. To minimize diurnal variability, all fMRI scans took place in the afternoon (range: 12:06–5:56 pm, mean ≈ 2:20 pm), and all female participants were scheduled during the follicular phase of the menstrual cycle (days 1–10 after the onset of menses), confirmed verbally prior to scanning.

**Self-report questionnaires to assess stress response over the course of the session**

Affective mood states were assessed repeatedly throughout the experimental session to characterize stress responses to the MAST, including recovery, using VAMS, PANAS, and the TCQ. See **Supplementary Fig. 1** for a timeline of the assessment timepoints. In detail, for the VAMS, participants used a sliding cursor on three scales: Happy-to-Sad, Friendly-to-Hostile, and Relax-to-Tense, each rated from 0–100 across five time points during the session. The PANAS state version consists of 20 adjectives describing affective states (10 positive, 10 negative), each rated on a 5-point Likert scale ranging from 1 (“very slightly or not at all”) to 5 (“extremely”) and was completed immediately before and after the MAST to capture changes in positive and negative affect in response to the stressor. Lastly, participants completed the TCQ to assess subjective task appraisals in the context of the MAST as employed in our prior study ^32^ and both pre-task and post-task versions were administered, each consisting of 11 statements rated on a 7-point Likert scale from 1 (“strongly disagree”) to 7 (“strongly agree”). The pre-task version asked participants to evaluate their expectations about the upcoming task (e.g., anticipated effort, perceived abilities to perform successfully, perceived control), whereas the post-task version asked participants to reflect on their experiences of the completed task (e.g., task demands, perceived performance, perceived control). Following established scoring procedures (e.g. ^32^), items were divided into demand (e.g., effort, task difficulty) and resource (e.g., ability, control) subscales. A demand-to-resource ratio was then computed, with values greater than 1 reflecting more threat-like appraisals (perceived demands outweighing resources), and values less than 1 reflecting more challenge-like appraisals (perceived resources outweighing demands). Higher resource scores thus indicate more adaptive challenge appraisals, whereas higher demand scores indicate more maladaptive threat appraisals.

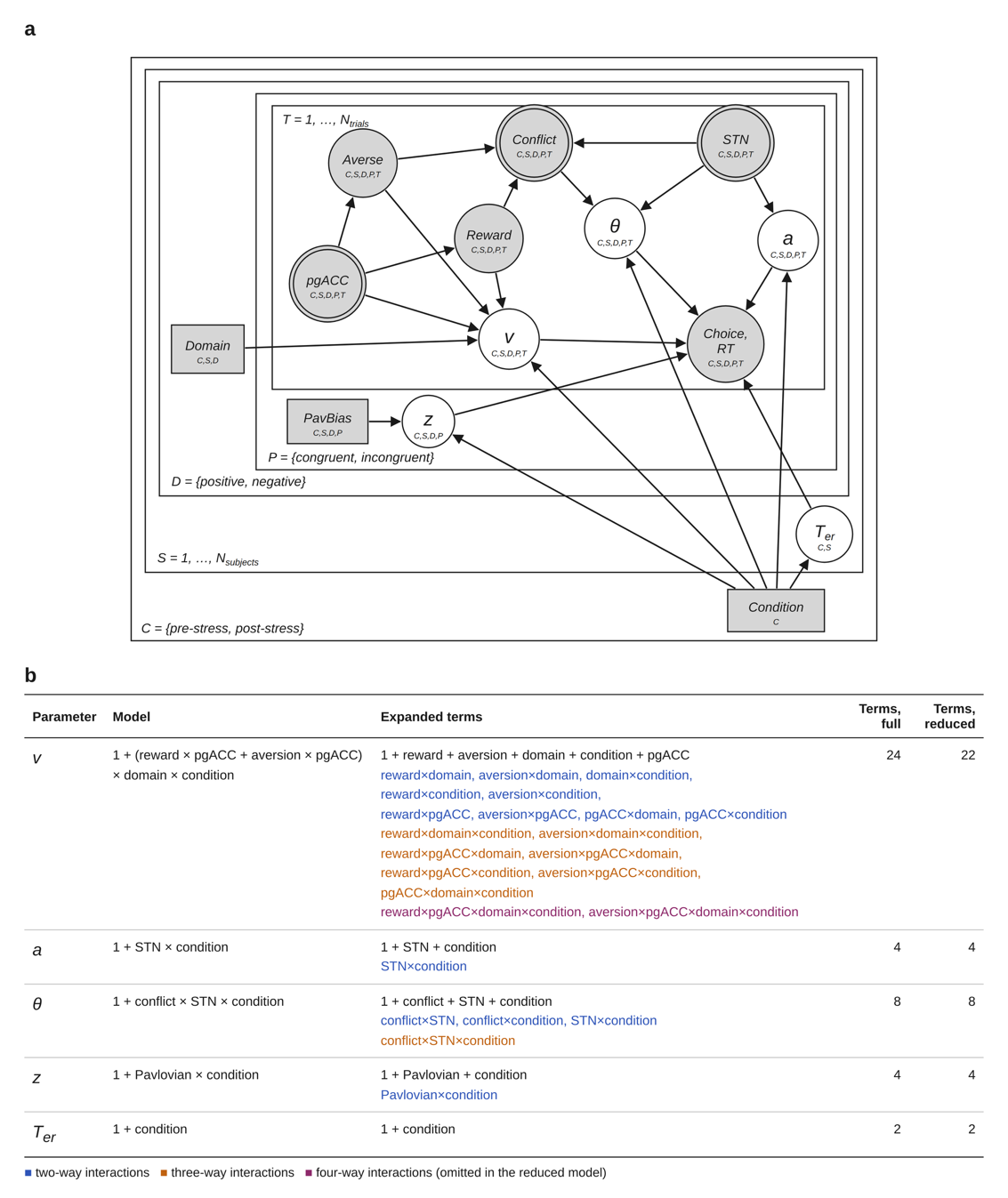

**Supplementary Fig. 3.** Neural model. (a) Graphical representation. Shaded nodes represent observed variables and unshaded nodes represent estimated parameters. Circles denote continuous variables and squares denote discrete variables. Trial-wise pgACC activity modulated drift rate (v) directly and in interaction with offered reward and aversion, while trial-wise STN activity modulated the decision threshold (a) and the rate of threshold collapse (θ). Starting point bias (z) varied by Pavlovian response congruency, and nondecision time (Ter) was estimated per participant. All parameters were allowed to vary by stress condition. Because the graph shows dependencies only, it applies both to the full model and to the reduced model reported in the main text. (b) Model definition for each parameter, listing all terms of the full model. The reduced model omits the two four-way interaction terms on drift rate (purple).

**Supplementary Results**

**Cortisol responses to the acute stress induction via the MAST**

To assess cortisol responses to the MAST, a repeated-measures ANOVA on log-transformed cortisol values was performed. One datapoint was missing at C5 for one healthy control (HC) participant (0.16% of all cortisol data points) and was imputed using the mice ^33^ package in R with Group and Timepoint as predictors. Results revealed a robust main effect of *Time* (F(2.80, 341.2) = 29.55, p < .001, partial η² = .20), reflecting the expected cortisol rise and recovery across the full session. There was no significant main effect of *Group* (F(1,122) = 1.47, p = .228, partial η² = .01) and no significant *Group* × *Time* interaction (F(2.80, 341.2) = 0.21, p = .875, partial η² = .002).

Follow-up pairwise comparisons confirmed that both groups showed significant cortisol increases from C3 (pre-stress baseline) to C4 (immediately post-MAST) and C5 (while under sustained stress and at expected peak, ~30 min post-stress offset). For example, in the HC group, cortisol rose significantly from C3 to C4 (p < .001) and C5 (p < .001) and then declined after sustained stress relief at C6 and C7 (all p < .001 vs. C5). A very similar temporal pattern was observed in the rMDD group, with robust increases from C3 to C4 (p < .001) and C5 (p < .001), followed by recovery after sustained stress relief at later time points. Importantly, none of the between-group contrasts at any time point reached significance after correction, indicating that cortisol reactivity and recovery trajectories were comparable across rMDD and HC participants. Notably, by the final sampling point (C7), cortisol values in both groups had declined to levels statistically indistinguishable from the pre-stress baseline, indicating full recovery of HPA-axis activity. Taken together, cortisol analyses demonstrated the expected robust stress-induced increase peaking at C5 and a successful induction of sustained stress, with no evidence for altered HPA-axis reactivity in the rMDD group relative to HC. See **Supplementary Fig. 4** for the cortisol response trajectory.

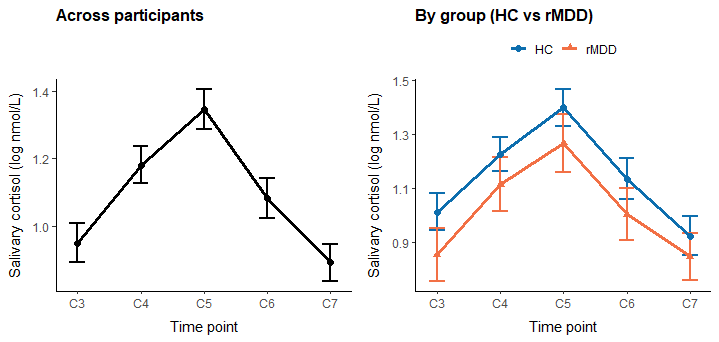

**Supplementary Fig. 4**. Salivary cortisol responses to the MAST. Mean log-transformed cortisol levels across participants (left) and separated by group (right) at pre-stress baseline (C3, ~0 min), immediately post-MAST (C4, ~+15 min), during the expected peak (C5, ~+45 min), and recovery (C6–C7; ~+60 and ~+90 min, respectively). Error bars represent standard error of the mean. Both groups showed a robust cortisol rise following stress induction, peaking at C5, followed by recovery to baseline levels. No significant group differences were observed at any time point.

**Subjective ratings pre/post-acute stress**

*PANAS Responses to the MAST*

For the PANAS, four missing datapoints were imputed in total for both subscales using the mice package ^33^. For negative affect, one datapoint was missing for one HC male participant before stress (0.4% of all observations). For positive affect, three datapoints were missing (two pre-stress from HC male participants and one post-stress from a female rMDD participant; 1.2% of all observations). The predictor matrix for imputation included Sex, Group, and Timepoint (pre/post stress), as well as the complementary PANAS subscale (positive or negative affect).

A repeated-measures ANOVA on Negative Affect revealed significant main effects of *Time* (F(1,122) = 93.01, p < .001, partial η² = .43) and *Group* (F(1,122) = 6.66, p = .011, partial η² = .05), as well as a significant *Group* × *Time* interaction (F(1,122) = 5.76, p = .018, partial η² = .05). These results indicate that the MAST successfully induced increases in negative affect, and that rMDD participants reported higher overall negative affect and, critically, exhibited a stronger stress-induced increase compared to HC.

Positive Affect analyses revealed a robust main effect of *Time* (F(1,122) = 57.27, p < .001, partial η² = .32), reflecting a clear decrease from pre- to post-MAST across participants, further confirming effective stress induction. There was also a trend-level main effect of *Group* (F(1,122) = 3.28, p = .073, partial η² = .03), suggesting that rMDD participants tended to report lower overall positive affect. However, the *Group* × *Time* interaction was not significant (F(1,122) = 0.28, p = .596, partial η² = .002), indicating that both groups exhibited a comparable decline in positive affect during the stressor.

See **Supplementary Fig. 5** for the PANAS negative affect (left) and positive affect (right) split by pre- and post-stress condition and group.

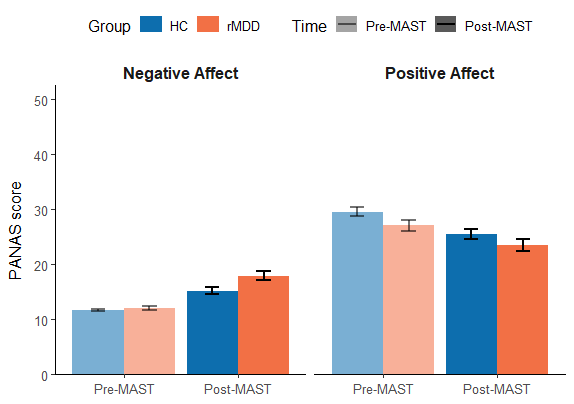

**Supplementary Fig. 5**. Positive and Negative Affect Schedule (PANAS) ratings pre- and post-MAST. Bar plots depict mean negative affect (left) and positive affect (right) in HC and rMDD. Both groups showed robust increases in negative affect and decreases in positive affect following the MAST, indicative of successful stress induction. rMDD participants reported higher overall negative affect, and the increase in negative affect was more pronounced relative to HC. Error bars represent standard error of the mean.

*Threat–Challenge Questionnaire (TCQ)*

Repeated-measures ANOVAs revealed robust main effects of *Time* for all TCQ indices, indicating that task appraisals shifted from pre- to post-MAST: Resources (F(1,122) = 114.50, p < .001, partial η² = .48), Demands (F(1,122) = 65.79, p < .001, partial η² = .35), and the demand-to-resource ratio (TCQ Score; F(1,122) = 63.53, p < .001, partial η² = .34). There were no significant overall *Group* differences (Resources: F(1,122) = 0.09, p = .762, partial η² = .001; Demands: F(1,122) = 2.72, p = .102, partial η² = .02; TCQ Score: F(1,122) = 1.99, p = .160, partial η² = .02). Critically, the *Group* × *Time* interaction was significant for Demands (F(1,122) = 5.43, p = .021, partial η² = .04) and the TCQ Score (F(1,122) = 6.56, p = .012, partial η² = .05), with a trend for Resources (F(1,122) = 3.75, p = .055, partial η² = .03). These results indicate that, while both groups reported the expected post-stress increase in perceived demands and increase in the demand-to-resource ratio, these post-stress shifts were more pronounced in the rMDD group. Because the TCQ score is the demand-to-resource ratio, higher values reflect more threat-like appraisals (perceived demands outweighing resources). The significant *Group* × *Time* effect indicates a stronger post-MAST shift toward threat-oriented appraisal in rMDD than in HC, consistent with the larger rise in the TCQ ratio in that group. See **Supplementary Fig. 6** for the TCQ facets of Resources (left), Demands (middle) and the TCQ Ratio (right) split by pre- and post-stress condition and group.

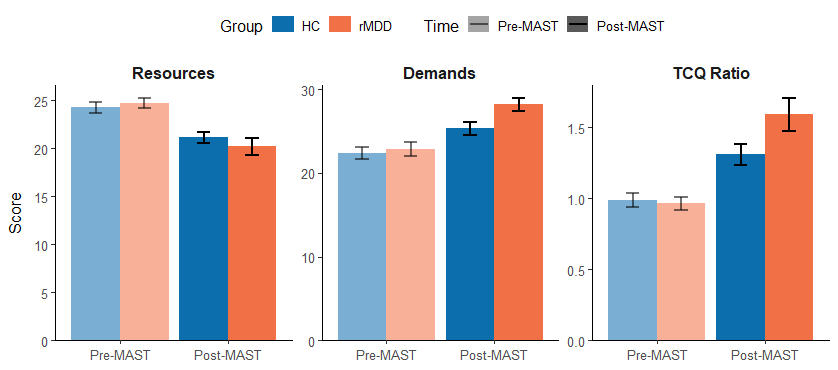

**Supplementary Fig. 6**. Threat–Challenge Questionnaire (TCQ) results before (Pre-MAST) and after (Post-MAST) stress induction, shown separately for Resources, Demands, and the TCQ Score ratio (Demands/Resources) indicating that rMDD participants appraised the stressor as more threatening, with a sharper shift in the demands/resources balance compared to HC. Error bars represent standard errors of the mean.

*Affective responding to MAST – Visual Analog Mood Scales (VAMS)*

To account for missing data, 18 datapoints (2.9% of all observations) were imputed using the mice ^33^ package. Missingness was limited to 9 HC participants (five male: one at T1, two at T3, two at T5; four female: one at T2, one at T4, two at T5) and, across these cases, involved all three VAMS subscales at least once. The imputation predictor matrix included Sex, Group, Timepoint (T1–T5), and the other subscales, respectively.

A repeated-measures MANOVA was conducted on the three affective VAMS scores across the five time points. At the multivariate level across the three outcomes’ joint distribution, there was a significant main effect of *Group* (Wilks’ lambda = .937, F(3,120) = 2.70, p = .049), with rMDD participants reporting overall higher negative affect. A main effect of *Time* also emerged (Wilks’ lambda = .306, F(12,111) = 20.95, p < .001). The *Group* × *Time* effect was not significant in this model (p = .109). In the multivariate within-subjects tests across the three VAMS measures, there was a robust main effect of *Time* (Wilks’ lambda = .507, F(12,1464) = 31.35, p < .001) and a significant *Group* × *Time* interaction (Wilks’ lambda = .952, F(12,1464) = 2.02, p = .020), indicating that the two groups differed in their affective trajectories across the task when mood scales were pooled. Follow-up univariate tests clarified the group differences. For between-subjects group differences, rMDD scored higher than HC on Happy-to-Sad (F(1,122) = 5.07, p = .026, partial η² = .040) and Relax-to-Tense (F(1,122) = 7.73, p = .006, partial η² = .060), but not Friendly-to-Hostile (F(1,122) = 2.72, p = .102, partial η² = .022). All three scales showed within-subjects *Time* effects (all p < .001). Critically, the *Group* × *Time* effect was significant for Friendly-to-Hostile (F(3.02,368.94) = 4.93, p = .002, partial η² = .039) and for Happy-to-Sad (F(3.41,416.54) = 2.52, p = .050, partial η² = .020), but not for Relax-to-Tense (F(3.47,423.41) = 1.49, p = .211, partial η² = .012). Inspection of estimated marginal means further revealed that the groups did not differ at the initial baseline on any of the scales (T1; all p > .10). Notably, at T2, after the first round of the dynamic AAC task and immediately prior to stress induction, rMDD participants already reported modestly higher negative affect than HC on the Happy-to-Sad and Relax-to-Tense scales (p = .041 and p = .027, respectively), while no difference emerged for Friendly-to-Hostile. Following the MAST (T3) and under sustained stress induction (T4), group differences were robust across all three scales (all p ≤ .024), indicating stronger stress-induced negative affect in rMDD. By the final assessment and after sustained stress relief (T5), no group differences were present (all p ≥ .387), consistent with recovery toward baseline. See **Supplementary Fig. 7** for the VAMS trajectories on dimensions of Happy-to-Sad (left), Friendly-to-Hostile (middle) and Relax-to-Tense (right) split by group.

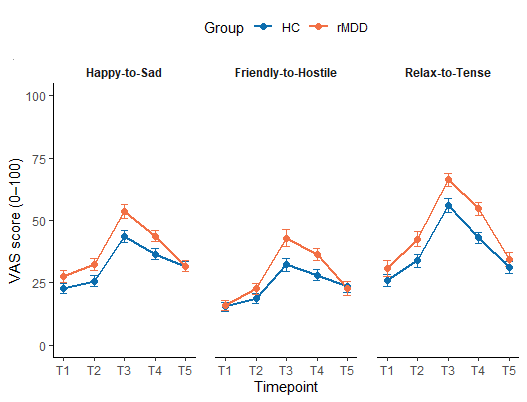

**Supplementary Fig. 7**. Visual Analog Mood Scale (VAMS) ratings across five time points (T1–T5; T1: session baseline before pre-stress task, T2: after pre-stress task and before MAST, T3: immediately after MAST, T4: during post-stress task under sustained stress, T5: after stress relief) for Happy-to-Sad, Friendly-to-Hostile, and Relax-to-Tense subscales in HC and rMDD participants. Both groups showed robust stress-related affective changes following the MAST (T3), with negative affect peaking under stress and declining toward baseline during recovery (T5). Compared to HC, rMDD participants reported higher negative affect overall, particularly at T2–T4, consistent with heightened stress reactivity. Error bars represent standard error of the mean.

**Behavioral-Only Model Results**

A behavioral-only model, fitted without neural regressors, was estimated to establish that the value-integration effects were not dependent on the inclusion of trial-wise BOLD signals. As in our prior work using this paradigm^9^, the Angle model provided the best behavioral fit, and the model included both pre- and post-stress conditions. Parameter estimates are reported in **Supplementary Table 1**. The behavioral-only model reproduced the asymmetric stress effect on value integration reported in the main text. On net-aversive offers, reward sensitivity increased credibly under stress in both groups, and this increase was credibly larger in rMDD than in HC (rMDD: β = 0.189, 95%-HDI = [0.124, 0.254]; HC: β = 0.100, 95%-HDI = [0.045, 0.155]; PP[rMDD > HC] = 0.977). As in the neural model, this increase was credibly offset in the positive domain in rMDD but not in HC (rMDD: β = -0.223, 95%-HDI = [-0.330, -0.120]; HC: β = -0.075, 95%-HDI = [-0.178, 0.027]; PP[rMDD > HC] = 0.023). Aversion sensitivity became credibly more negative in rMDD but not in HC on these offers, and the group difference was at our criterion (rMDD: β = -0.109, 95%-HDI = [-0.176, -0.039]; HC: β = -0.028, 95%-HDI = [-0.090, 0.031]; PP[rMDD > HC] = 0.050). The corresponding parameter estimates are visualized in **Supplementary Fig. 8**, and posterior predictive checks for this model are shown in **Supplementary Fig. 9.**

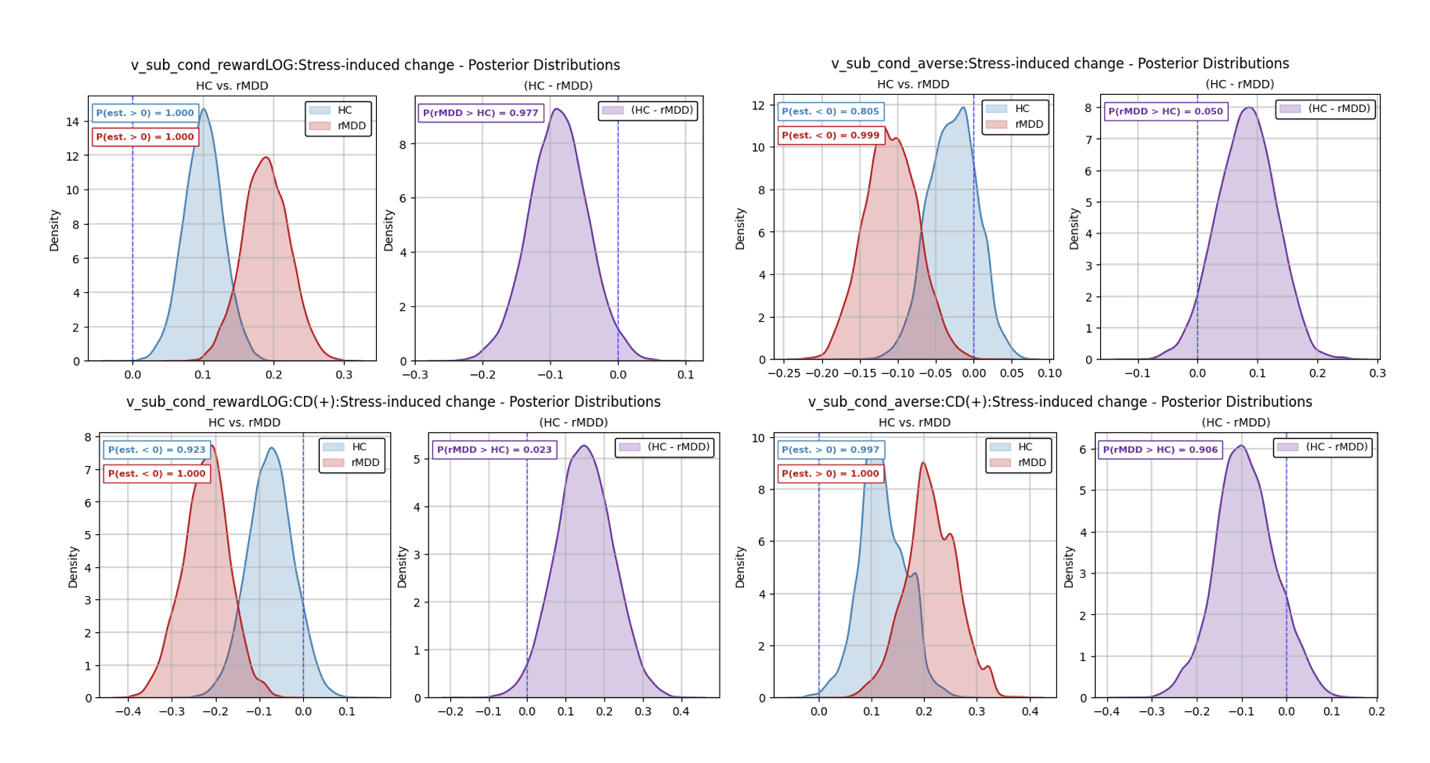

**Supplementary Fig. 8.** Stress-induced changes in drift-rate sensitivity in the behavioral-only model. Panels show the change in reward sensitivity (top-left) and aversion sensitivity (top-right) on net-aversive offers, and the additional change on net-rewarding offers, relative to net-aversive offers, for reward (bottom-left) and aversion (bottom-right). Single-group distributions (HC in blue, rMDD in red) show the posterior of each parameter within a group, annotated with the posterior probability that the estimate is above zero. Group-difference distributions (purple) show the posterior of the sampled difference HC − rMDD, annotated with the posterior probability that the estimate is larger in rMDD than in HC, PP[rMDD > HC].

**Neural Model Results**

**Supplementary Tables 2** and **3** report the full parameter estimates for the neural model presented in the main text. **Supplementary Table 2** shows pre-stress estimates, derived from the posterior distributions by subtracting the stress-induced change from the post-stress reference. **Supplementary Table 3** shows the post-stress estimates together with the stress-induced changes.

**Stress effects shared across groups**

Two further effects of stress were shared across groups. Both groups showed credibly shorter nondecision time following stress, with no credible group difference (HC: β = -0.053, 95%-HDI = [-0.089, -0.017]; rMDD: β = -0.050, 95%-HDI = [-0.085, -0.014]; PP[rMDD > HC] = 0.553), indicating a stress-induced speeding of sensorimotor processing. However, given the tradeoff between decision threshold and nondecision time in sequential sampling models, this effect may partly reflect the broader shift toward faster decision dynamics under stress rather than a fully distinct computational process. Stress also altered the starting point bias. Before stress, Pavlovian response congruency had no credible effect on the starting point in either group (rMDD: β = 0.004, 95%-HDI = [-0.014, 0.022]; HC: β = 0.003, 95%-HDI = [-0.015, 0.021]; PP[rMDD > HC] = 0.557). After stress, a credible congruency effect was present in both groups (rMDD: β = 0.027, 95%-HDI = [0.011, 0.044]; HC: β = 0.020, 95%-HDI = [0.005, 0.034]; PP[rMDD > HC] = 0.745). The stress-induced change did not differ between groups (PP[rMDD > HC] = 0.671), although within groups it was credible in rMDD (β = 0.023, 95%-HDI = [0.004, 0.041]) and fell just short in HC (β = 0.017, 95%-HDI = [-0.001, 0.035]). Stress therefore introduced a Pavlovian influence on the a priori tendency to approach or avoid that was not present beforehand, and it did so comparably in both groups.

**Comparison of the full and reduced neural models**

The neural model reported in the main text excludes interaction terms of fourth order and above (reduced model), while another model including such interactions (full model) was also tested. Posterior predictive checks for both models are shown in **Supplementary Figs. 10 and 11**. To verify that this reduction did not alter findings substantively, we compared parameter estimates between the full model and the reduced model across all model parameters (**Supplementary Figs. 12–15**).

Estimates were closely comparable between the two models throughout. For drift-rate terms based on task variables, posterior means differed by less than 0.01 in nearly all cases, and the posterior distributions of the group difference were almost completely overlapping (**Supplementary Fig. 12**). Differences were slightly larger for the pgACC regressors on drift rate, where posterior means shifted by up to 0.037 (**Supplementary Fig. 13**). Estimates were essentially unchanged for the STN regressors on the decision threshold and its rate of collapse (**Supplementary Fig. 14**) and for the starting-point parameters (**Supplementary Fig. 15**). Directional conclusions were unaffected.

Across the 59 parameters estimated in both models, no group difference changed credibility status. Four within-group pgACC estimates crossed the credibility boundary, each with an interval bound within 0.012 of zero in both models. Because the reduced model retains all terms up to third order, including the three-way interactions on which the reported effects depend, and because the higher-order terms omitted from it were not credible in the full model, we report the reduced model in the main text and present the full model here as a robustness check (**Supplementary Table 4**).

**Supplementary Discussion**

**Interpretation of the pgACC change of sign**

The change of sign constrains the interpretation of the pgACC signal, although less than it first appears. A salience signal that adds to evidence accumulation irrespective of valence would shift drift in the same direction in both domains, which is not what we observed. A signal that instead scales the evidence an offer already favors, acting as a gain, would amplify accumulation toward avoidance on net-aversive offers and toward approach on net-rewarding ones, which matches the observed pattern. Valuation and regulatory accounts accommodate the change of sign as well, since both permit the region's contribution to depend on whether the offer is net-rewarding or net-aversive. The present data therefore support valence-dependent coupling and are compatible with gain-like amplification, but not with the stronger claim that pgACC encodes aversion.

**Relation to earlier work with this task**

These results can be situated against earlier work, from which they differ in one respect and converge in another. In currently depressed individuals, the computational locus of the group difference was the starting point of accumulation, with nucleus accumbens activity biasing approach in HC and avoidance in MDD^34^. Here, in remission and under stress, the divergence lies in drift-rate coupling with pgACC and in threshold coupling with the subthalamic nucleus, while the starting point showed a stress effect that did not differ by group. The locus of computational group differences may therefore shift with clinical state. By contrast, in an unselected sample performing this task without a stressor, conflict prolonged threshold collapse across the whole group^9^, and an effect of comparable magnitude was present in rMDD pre-stress.

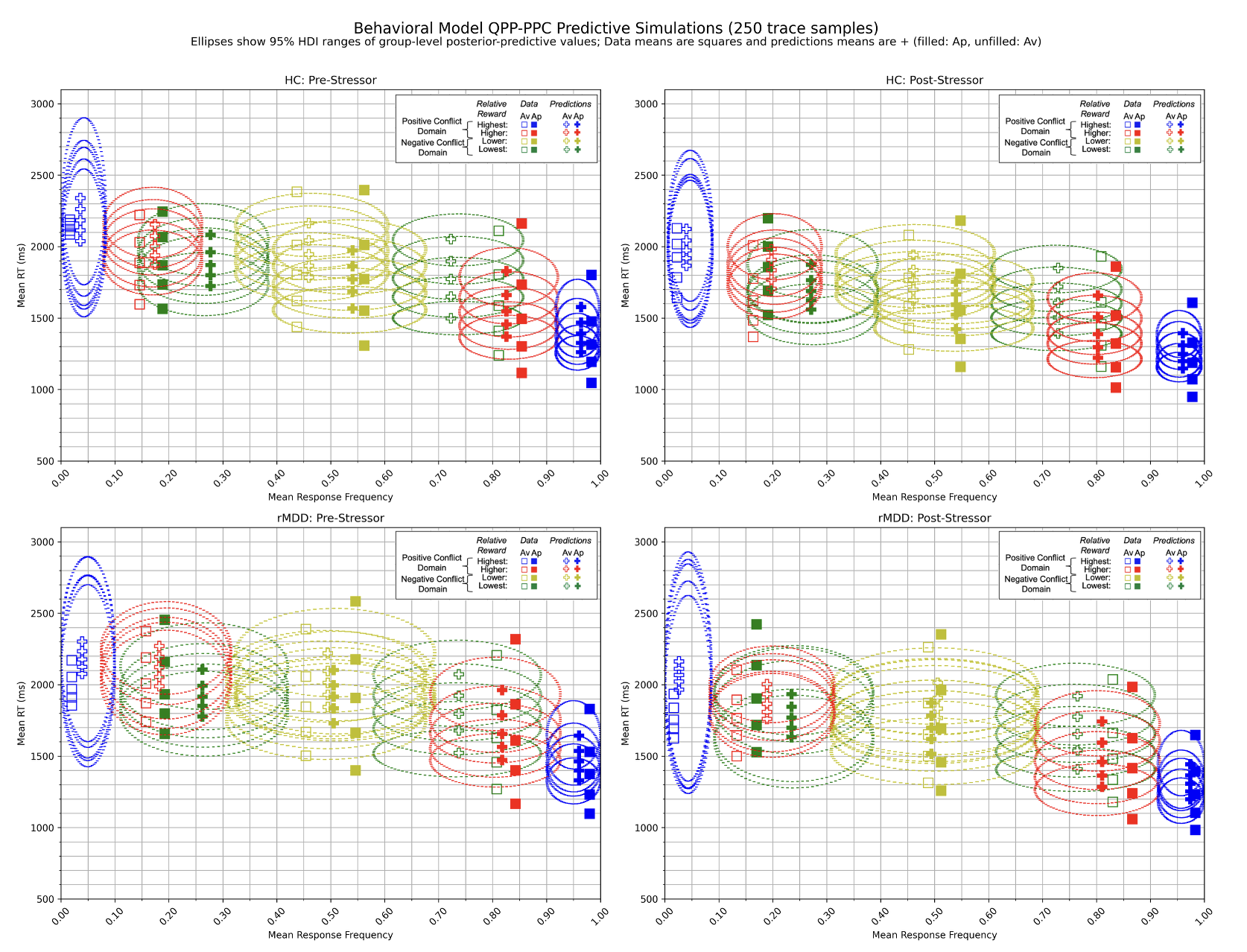

**Supplementary Fig. 9.** Posterior predictive checks by group of behavioral-only model for HC pre-stress (top-left), HC post-stress (top-right), rMDD pre-stress (bottom-left), and rMDD post-stress (bottom-right). Ellipse horizontal widths reflect the 95% HDI for response frequency, while vertical widths represent the 95% HDI for mean RT percentiles at 10%, 30%, 50%, 70%, and 90%. Squares indicate observed data means and plus symbols indicate model-predicted means, with filled symbols denoting approach and unfilled symbols denoting avoidance choices. Colors reflect the bin of relative reward evidence that each data series belongs to (highest and higher in the positive domain, lower and lowest in the negative domain), with bin boundaries computed within each participant. 250 sample traces were used to simulate the entire dataset, yielding 250 simulated datasets.

**Supplementary Table 1. Estimates from the behavioral model fitted without neural regressors.**

| **Model term** | **rMDD mean (SD)** | **rMDD 95% HDI** | **HC mean (SD)** | **HC 95% HDI** | **PP[rMDD > HC]** |
| --- | --- | --- | --- | --- | --- |
| ***Drift rate (v)*** |  |  |  |  |  |
| Intercept, pre-stress | -0.003 (0.114) | [-0.227, 0.222] | 0.053 (0.104) | [-0.152, 0.258] | 0.357 |
| Intercept | 0.025 (0.108) | [-0.186, 0.236] | 0.112 (0.096) | [-0.074, 0.302] | 0.270 |
| Intercept, stress-induced change | 0.028 (0.064) | [-0.098, 0.154] | 0.059 (0.059) | [-0.056, 0.176] | 0.363 |
| Reward, pre-stress | **0.623 (0.049)** | **[0.527, 0.718]** | **0.663 (0.038)** | **[0.588, 0.738]** | 0.257 |
| Reward | **0.811 (0.047)** | **[0.718, 0.905]** | **0.762 (0.037)** | **[0.690, 0.835]** | 0.791 |
| Reward, stress-induced change | **0.189 (0.033)** | **[0.124, 0.254]** | **0.100 (0.028)** | **[0.045, 0.155]** | **0.977** |
| Aversion, pre-stress | **-0.626 (0.043)** | **[-0.710, -0.543]** | **-0.661 (0.036)** | **[-0.731, -0.590]** | 0.737 |
| Aversion | **-0.735 (0.043)** | **[-0.819, -0.650]** | **-0.689 (0.038)** | **[-0.764, -0.614]** | 0.215 |
| Aversion, stress-induced change | **-0.109 (0.035)** | **[-0.176, -0.039]** | -0.028 (0.032) | [-0.090, 0.031] | **0.050** |
| Positive domain, pre-stress | **1.076 (0.106)** | **[0.866, 1.283]** | **1.063 (0.097)** | **[0.870, 1.252]** | 0.535 |
| Positive domain | **1.329 (0.102)** | **[1.131, 1.528]** | **1.199 (0.090)** | **[1.024, 1.378]** | 0.831 |
| Positive domain, stress-induced change | **0.253 (0.076)** | **[0.103, 0.402]** | 0.137 (0.071) | [-0.003, 0.276] | 0.870 |
| Reward, positive domain, pre-stress | **-0.325 (0.058)** | **[-0.438, -0.209]** | **-0.388 (0.062)** | **[-0.509, -0.267]** | 0.778 |
| Reward, positive domain | **-0.547 (0.058)** | **[-0.659, -0.434]** | **-0.463 (0.057)** | **[-0.575, -0.350]** | 0.148 |
| Reward, positive domain, stress-induced change | **-0.223 (0.053)** | **[-0.330, -0.120]** | -0.075 (0.052) | [-0.178, 0.027] | **0.023** |
| Aversion, positive domain, pre-stress | **0.218 (0.062)** | **[0.096, 0.341]** | **0.266 (0.057)** | **[0.155, 0.378]** | 0.283 |
| Aversion, positive domain | **0.430 (0.064)** | **[0.302, 0.556]** | **0.387 (0.059)** | **[0.272, 0.503]** | 0.695 |
| Aversion, positive domain, stress-induced change | **0.212 (0.050)** | **[0.112, 0.316]** | **0.121 (0.044)** | **[0.037, 0.206]** | 0.906 |
| ***Decision threshold (a)*** |  |  |  |  |  |
| Intercept, pre-stress | **1.882 (0.068)** | **[1.751, 2.019]** | **1.781 (0.051)** | **[1.681, 1.883]** | 0.885 |
| Intercept | **1.763 (0.069)** | **[1.633, 1.903]** | **1.658 (0.049)** | **[1.564, 1.757]** | 0.897 |
| Intercept, stress-induced change | **-0.119 (0.056)** | **[-0.235, -0.012]** | **-0.123 (0.045)** | **[-0.211, -0.034]** | 0.530 |
| Conflict, pre-stress | -0.036 (0.019) | [-0.073, 0.003] | **-0.050 (0.017)** | **[-0.083, -0.016]** | 0.705 |
| Conflict | **-0.054 (0.017)** | **[-0.088, -0.020]** | **-0.062 (0.016)** | **[-0.094, -0.031]** | 0.631 |
| Conflict, stress-induced change | -0.018 (0.024) | [-0.064, 0.028] | -0.012 (0.021) | [-0.056, 0.028] | 0.431 |
| ***Rate of threshold collapse (θ)*** |  |  |  |  |  |
| Intercept, pre-stress | **0.567 (0.026)** | **[0.516, 0.617]** | **0.555 (0.021)** | **[0.515, 0.595]** | 0.648 |
| Intercept | **0.567 (0.025)** | **[0.517, 0.615]** | **0.563 (0.020)** | **[0.524, 0.601]** | 0.549 |
| Intercept, stress-induced change | -0.000 (0.026) | [-0.053, 0.050] | 0.008 (0.022) | [-0.035, 0.051] | 0.410 |
| Conflict, pre-stress | **-0.040 (0.012)** | **[-0.064, -0.015]** | **-0.029 (0.012)** | **[-0.052, -0.003]** | 0.252 |
| Conflict | **-0.025 (0.012)** | **[-0.048, -0.000]** | **-0.041 (0.013)** | **[-0.067, -0.016]** | 0.817 |
| Conflict, stress-induced change | 0.016 (0.017) | [-0.016, 0.048] | -0.012 (0.017) | [-0.047, 0.019] | 0.877 |
| ***Starting point bias (z)*** |  |  |  |  |  |
| Intercept, pre-stress | **0.540 (0.009)** | **[0.522, 0.557]** | **0.543 (0.007)** | **[0.528, 0.557]** | 0.396 |
| Intercept | **0.541 (0.010)** | **[0.522, 0.560]** | **0.540 (0.008)** | **[0.525, 0.557]** | 0.486 |
| Intercept, stress-induced change | 0.005 (0.010) | [-0.016, 0.025] | -0.009 (0.008) | [-0.025, 0.008] | 0.842 |
| Pavlovian incongruent, pre-stress | 0.004 (0.009) | [-0.014, 0.021] | 0.001 (0.009) | [-0.017, 0.020] | 0.564 |
| Pavlovian incongruent | **0.027 (0.009)** | **[0.011, 0.044]** | **0.020 (0.008)** | **[0.005, 0.035]** | 0.738 |
| Pavlovian incongruent, stress-induced change | **0.024 (0.010)** | **[0.005, 0.042]** | **0.019 (0.009)** | **[0.000, 0.037]** | 0.644 |
| ***Nondecision time (Tₑᵣ)*** |  |  |  |  |  |
| Intercept, pre-stress | **0.701 (0.023)** | **[0.656, 0.744]** | **0.685 (0.021)** | **[0.644, 0.728]** | 0.688 |
| Intercept | **0.643 (0.019)** | **[0.604, 0.680]** | **0.630 (0.016)** | **[0.598, 0.662]** | 0.701 |
| Intercept, stress-induced change | **-0.058 (0.019)** | **[-0.094, -0.021]** | **-0.056 (0.019)** | **[-0.093, -0.020]** | 0.472 |

Note. Values are posterior means with standard deviations (SD) and 95% highest density intervals (HDI). Bold within-group values indicate that the 95% HDI excludes zero. PP[rMDD > HC] is the posterior probability that a parameter is larger in rMDD than in HC, defined as the proportion of the posterior distribution of the group difference lying above zero; bold values indicate PP >= 0.95 or PP <= 0.05. Terms are grouped by model parameter. Unless labelled positive domain, estimates refer to the negative domain, which served as the reference. Interactions are denoted by x. Terms labelled stress-induced change represent the difference between conditions (post minus pre-stress); terms labelled pre-stress are estimates for the pre-stress condition derived from the same posteriors.

**Supplementary Table 2. Pre-stress parameter estimates, neural model.**

| **Model term** | **rMDD mean (SD)** | **rMDD 95% HDI** | **HC mean (SD)** | **HC 95% HDI** | **PP[rMDD > HC]** |
| --- | --- | --- | --- | --- | --- |
| ***Drift rate (v)*** |  |  |  |  |  |
| Intercept | -0.007 (0.116) | [-0.236, 0.222] | 0.042 (0.101) | [-0.154, 0.241] | 0.371 |
| Reward | **0.619 (0.050)** | **[0.521, 0.718]** | **0.662 (0.038)** | **[0.588, 0.738]** | 0.243 |
| Aversion | **-0.619 (0.043)** | **[-0.705, -0.535]** | **-0.658 (0.037)** | **[-0.731, -0.586]** | 0.759 |
| Positive domain | **1.082 (0.102)** | **[0.881, 1.284]** | **1.067 (0.098)** | **[0.872, 1.259]** | 0.548 |
| Reward, positive domain | **-0.342 (0.057)** | **[-0.455, -0.229]** | **-0.400 (0.061)** | **[-0.519, -0.281]** | 0.753 |
| Aversion, positive domain | **0.222 (0.063)** | **[0.099, 0.345]** | **0.275 (0.057)** | **[0.163, 0.386]** | 0.269 |
| pgACC | -0.007 (0.028) | [-0.063, 0.049] | **-0.076 (0.025)** | **[-0.126, -0.028]** | **0.968** |
| Reward × pgACC | -0.016 (0.023) | [-0.062, 0.031] | -0.033 (0.020) | [-0.072, 0.005] | 0.700 |
| Aversion × pgACC | -0.026 (0.023) | [-0.072, 0.021] | **0.050 (0.021)** | **[0.010, 0.090]** | **0.010** |
| pgACC, positive domain | -0.003 (0.048) | [-0.096, 0.091] | **0.084 (0.040)** | **[0.009, 0.165]** | 0.076 |
| ***Decision threshold (a)*** |  |  |  |  |  |
| Intercept | **1.892 (0.069)** | **[1.759, 2.029]** | **1.818 (0.053)** | **[1.716, 1.923]** | 0.804 |
| STN | **0.050 (0.018)** | **[0.012, 0.085]** | **0.043 (0.014)** | **[0.015, 0.069]** | 0.622 |
| ***Rate of threshold collapse (θ)*** |  |  |  |  |  |
| Intercept | **0.581 (0.024)** | **[0.533, 0.629]** | **0.584 (0.021)** | **[0.543, 0.625]** | 0.472 |
| Conflict | **-0.025 (0.009)** | **[-0.042, -0.007]** | -0.003 (0.010) | [-0.023, 0.016] | 0.052 |
| STN | **-0.025 (0.012)** | **[-0.048, -0.002]** | **-0.030 (0.009)** | **[-0.049, -0.012]** | 0.653 |
| Conflict × STN | 0.015 (0.009) | [-0.002, 0.032] | 0.008 (0.007) | [-0.006, 0.022] | 0.736 |
| ***Starting point bias (z)*** |  |  |  |  |  |
| Intercept | **0.545 (0.009)** | **[0.527, 0.563]** | **0.545 (0.008)** | **[0.530, 0.560]** | 0.485 |
| Pavlovian incongruent | 0.004 (0.009) | [-0.014, 0.022] | 0.003 (0.009) | [-0.015, 0.021] | 0.557 |
| ***Nondecision time (Tₑᵣ)*** |  |  |  |  |  |
| Intercept | **0.703 (0.022)** | **[0.660, 0.747]** | **0.682 (0.021)** | **[0.640, 0.724]** | 0.753 |

Note. Pre-stress estimates were derived from the posterior distributions by subtracting the stress-induced change from the corresponding post-stress estimate. Values are posterior means with standard deviations (SD) and 95% highest density intervals (HDI). Bold within-group values indicate that the 95% HDI excludes zero. PP[rMDD > HC] is the posterior probability that a parameter is larger in rMDD than in HC, defined as the proportion of the posterior distribution of the group difference lying above zero; bold values indicate PP >= 0.95 or PP <= 0.05. Terms are grouped by model parameter. Unless labelled positive domain, estimates refer to the negative domain, which served as the reference. Interactions are denoted by x. Terms labelled stress-induced change represent the difference between conditions (post minus pre-stress); terms labelled pre-stress are estimates for the pre-stress condition derived from the same posteriors.

**Supplementary Table 3. Post-stress parameter estimates and stress-induced change, neural model.**

| **Model term** | **rMDD mean (SD)** | **rMDD 95% HDI** | **HC mean (SD)** | **HC 95% HDI** | **PP[rMDD > HC]** |
| --- | --- | --- | --- | --- | --- |
| ***Drift rate (v)*** |  |  |  |  |  |
| Intercept | 0.009 (0.108) | [-0.205, 0.220] | 0.102 (0.095) | [-0.081, 0.291] | 0.255 |
| Intercept, stress-induced change | 0.016 (0.064) | [-0.108, 0.143] | 0.060 (0.059) | [-0.055, 0.177] | 0.303 |
| Reward | **0.807 (0.048)** | **[0.712, 0.902]** | **0.760 (0.037)** | **[0.687, 0.834]** | 0.781 |
| Reward, stress-induced change | **0.188 (0.033)** | **[0.124, 0.252]** | **0.098 (0.028)** | **[0.044, 0.152]** | **0.982** |
| Aversion | **-0.725 (0.044)** | **[-0.813, -0.639]** | **-0.677 (0.039)** | **[-0.753, -0.599]** | 0.201 |
| Aversion, stress-induced change | **-0.106 (0.037)** | **[-0.178, -0.032]** | -0.019 (0.033) | [-0.083, 0.042] | **0.035** |
| Positive domain | **1.347 (0.100)** | **[1.153, 1.541]** | **1.208 (0.089)** | **[1.034, 1.383]** | 0.852 |
| Positive domain, stress-induced change | **0.265 (0.075)** | **[0.118, 0.411]** | 0.141 (0.072) | [-0.003, 0.281] | 0.883 |
| Reward, positive domain | **-0.570 (0.059)** | **[-0.684, -0.455]** | **-0.484 (0.057)** | **[-0.596, -0.371]** | 0.142 |
| Reward, positive domain, stress-induced change | **-0.228 (0.054)** | **[-0.330, -0.120]** | -0.084 (0.053) | [-0.190, 0.020] | **0.029** |
| Aversion, positive domain | **0.436 (0.064)** | **[0.311, 0.566]** | **0.396 (0.059)** | **[0.282, 0.514]** | 0.680 |
| Aversion, positive domain, stress-induced change | **0.214 (0.053)** | **[0.113, 0.330]** | **0.121 (0.046)** | **[0.028, 0.208]** | 0.918 |
| pgACC | **-0.079 (0.023)** | **[-0.126, -0.034]** | -0.027 (0.022) | [-0.071, 0.015] | 0.052 |
| pgACC, stress-induced change | **-0.073 (0.033)** | **[-0.142, -0.011]** | 0.049 (0.030) | [-0.009, 0.107] | **0.001** |
| Reward × pgACC | -0.006 (0.020) | [-0.046, 0.032] | 0.002 (0.016) | [-0.027, 0.034] | 0.374 |
| Reward × pgACC, stress-induced change | 0.010 (0.027) | [-0.043, 0.061] | 0.035 (0.021) | [-0.007, 0.076] | 0.239 |
| Aversion × pgACC | 0.041 (0.022) | [-0.000, 0.082] | **0.051 (0.019)** | **[0.012, 0.085]** | 0.370 |
| Aversion × pgACC, stress-induced change | **0.067 (0.026)** | **[0.018, 0.117]** | 0.002 (0.021) | [-0.041, 0.042] | **0.978** |
| pgACC, positive domain | **0.171 (0.036)** | **[0.103, 0.243]** | 0.050 (0.035) | [-0.018, 0.117] | **0.992** |
| pgACC, positive domain, stress-induced change | **0.174 (0.058)** | **[0.069, 0.290]** | -0.035 (0.048) | [-0.133, 0.060] | **0.998** |
| Reward × pgACC, positive domain | -0.013 (0.027) | [-0.064, 0.041] | -0.013 (0.022) | [-0.059, 0.028] | 0.486 |
| Aversion × pgACC, positive domain | -0.017 (0.025) | [-0.066, 0.026] | -0.041 (0.023) | [-0.084, 0.006] | 0.734 |
| ***Decision threshold (a)*** |  |  |  |  |  |
| Intercept | **1.742 (0.068)** | **[1.614, 1.879]** | **1.683 (0.051)** | **[1.585, 1.786]** | 0.756 |
| Intercept, stress-induced change | **-0.149 (0.055)** | **[-0.259, -0.042]** | **-0.134 (0.045)** | **[-0.226, -0.047]** | 0.418 |
| STN | **0.081 (0.014)** | **[0.051, 0.107]** | 0.024 (0.012) | [-0.001, 0.048] | **0.999** |
| STN, stress-induced change | 0.031 (0.022) | [-0.012, 0.075] | -0.019 (0.018) | [-0.053, 0.020] | **0.952** |
| ***Rate of threshold collapse (θ)*** |  |  |  |  |  |
| Intercept | **0.557 (0.025)** | **[0.509, 0.607]** | **0.580 (0.020)** | **[0.541, 0.618]** | 0.244 |
| Intercept, stress-induced change | -0.024 (0.026) | [-0.074, 0.028] | -0.004 (0.022) | [-0.047, 0.039] | 0.274 |
| Conflict | 0.008 (0.010) | [-0.012, 0.028] | -0.002 (0.009) | [-0.020, 0.017] | 0.764 |
| Conflict, stress-induced change | **0.033 (0.013)** | **[0.007, 0.056]** | 0.002 (0.012) | [-0.022, 0.025] | **0.961** |
| STN | 0.014 (0.010) | [-0.008, 0.033] | **-0.037 (0.009)** | **[-0.055, -0.020]** | **1.000** |
| STN, stress-induced change | **0.038 (0.015)** | **[0.008, 0.068]** | -0.007 (0.012) | [-0.030, 0.018] | **0.987** |
| Conflict × STN | -0.000 (0.009) | [-0.017, 0.018] | -0.002 (0.007) | [-0.017, 0.011] | 0.591 |
| Conflict × STN, stress-induced change | -0.015 (0.012) | [-0.039, 0.010] | -0.010 (0.009) | [-0.029, 0.008] | 0.381 |
| ***Starting point bias (z)*** |  |  |  |  |  |
| Intercept | **0.545 (0.010)** | **[0.526, 0.565]** | **0.543 (0.008)** | **[0.527, 0.559]** | 0.581 |
| Intercept, stress-induced change | 0.007 (0.010) | [-0.014, 0.027] | -0.009 (0.008) | [-0.026, 0.007] | 0.883 |
| Pavlovian incongruent | **0.027 (0.008)** | **[0.011, 0.044]** | **0.020 (0.007)** | **[0.005, 0.034]** | 0.745 |
| Pavlovian incongruent, stress-induced change | **0.023 (0.009)** | **[0.004, 0.041]** | 0.017 (0.009) | [-0.001, 0.035] | 0.671 |
| ***Nondecision time (Tₑᵣ)*** |  |  |  |  |  |
| Intercept | **0.653 (0.019)** | **[0.617, 0.691]** | **0.629 (0.016)** | **[0.597, 0.661]** | 0.840 |
| Intercept, stress-induced change | **-0.050 (0.018)** | **[-0.085, -0.014]** | **-0.053 (0.018)** | **[-0.089, -0.017]** | 0.553 |

Note. The post-stress condition served as the reference; terms without a condition label are post-stress estimates. Values are posterior means with standard deviations (SD) and 95% highest density intervals (HDI). Bold within-group values indicate that the 95% HDI excludes zero. PP[rMDD > HC] is the posterior probability that a parameter is larger in rMDD than in HC, defined as the proportion of the posterior distribution of the group difference lying above zero; bold values indicate PP >= 0.95 or PP <= 0.05. Terms are grouped by model parameter. Unless labelled positive domain, estimates refer to the negative domain, which served as the reference. Interactions are denoted by x. Terms labelled stress-induced change represent the difference between conditions (post minus pre-stress); terms labelled pre-stress are estimates for the pre-stress condition derived from the same posteriors.

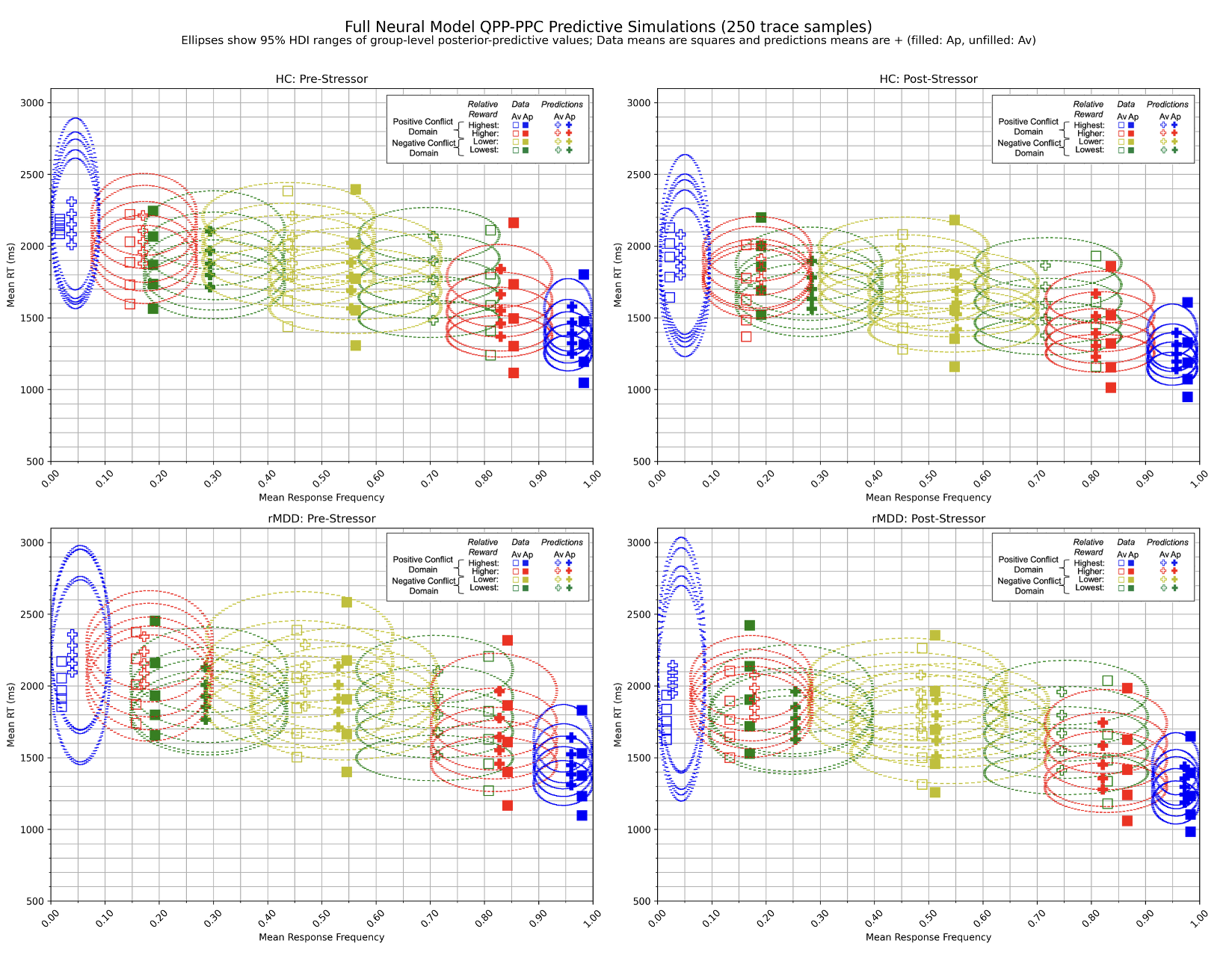

**Supplementary Fig. 10.** Posterior predictive checks by group of the full neural model (no terms removed) for HC pre-stress (top-left), HC post-stress (top-right), rMDD pre-stress (bottom-left), and rMDD post-stress (bottom-right). Ellipse horizontal widths reflect the 95% HDI for response frequency, while vertical widths represent the 95% HDI for mean RT percentiles at 10%, 30%, 50%, 70%, and 90%. Squares indicate observed data means and plus symbols indicate model-predicted means, with filled symbols denoting approach and unfilled symbols denoting avoidance choices. Colors reflect the bin of relative reward evidence that each data series belongs to (highest and higher in the positive domain, lower and lowest in the negative domain), with bin boundaries computed within each participant. 250 sample traces were used to simulate the entire dataset, yielding 250 simulated datasets.

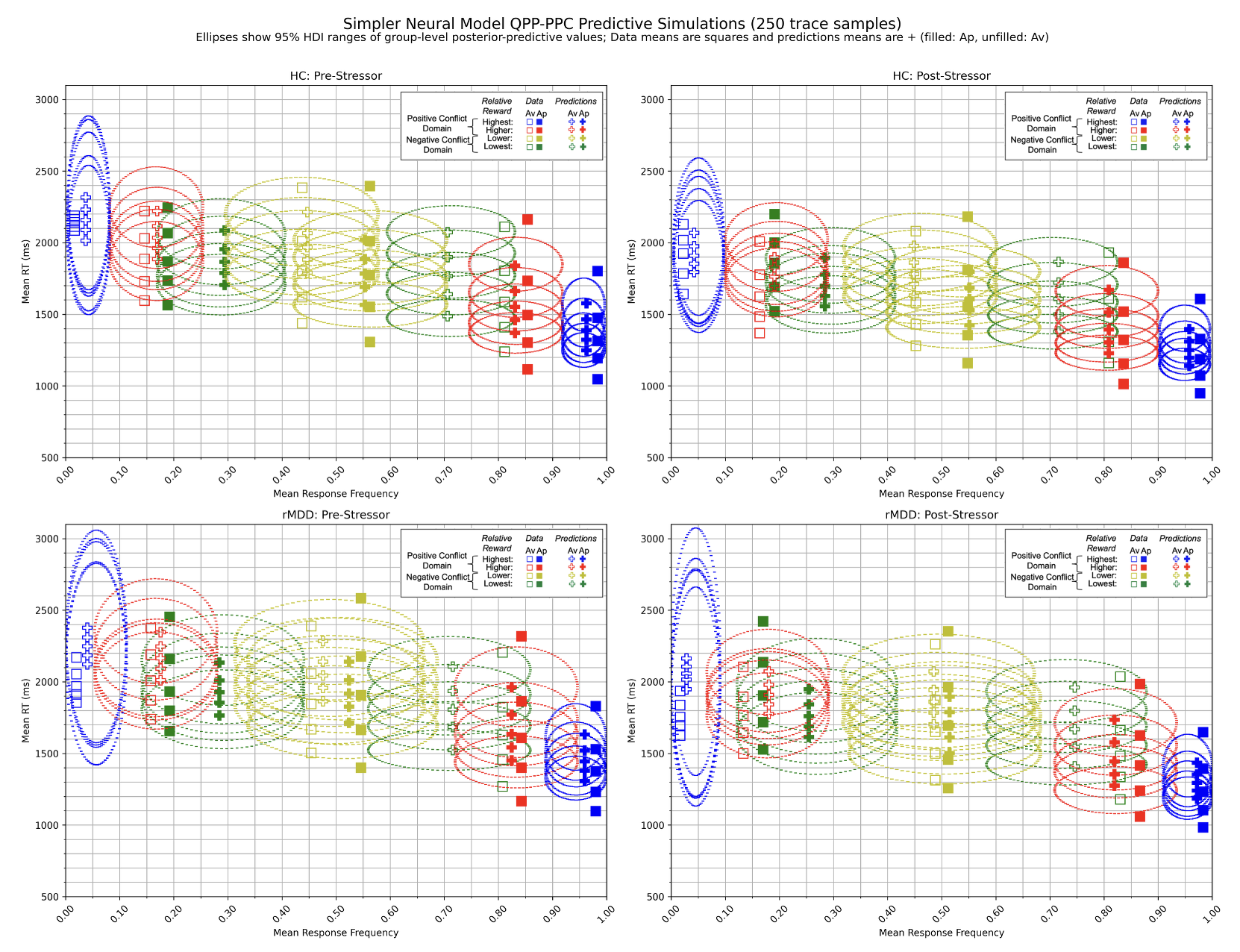

**Supplementary Fig. 11.** Posterior predictive checks by group of the reduced neural model for HC pre-stress (top-left), HC post-stress (top-right), rMDD pre-stress (bottom-left), and rMDD post-stress (bottom-right). Ellipse horizontal widths reflect the 95% HDI for response frequency, while vertical widths represent the 95% HDI for mean RT percentiles at 10%, 30%, 50%, 70%, and 90%. Squares indicate observed data means and plus symbols indicate model-predicted means, with filled symbols denoting approach and unfilled symbols denoting avoidance choices. Colors reflect the bin of relative reward evidence that each data series belongs to (highest and higher in the positive domain, lower and lowest in the negative domain), with bin boundaries computed within each participant. 250 sample traces were used to simulate the entire dataset, yielding 250 simulated datasets.

**Supplementary Table 4. Full neural model parameter estimates.**

| **Model term** | **rMDD mean (SD)** | **rMDD 95% HDI** | **HC mean (SD)** | **HC 95% HDI** | **PP[rMDD > HC]** |
| --- | --- | --- | --- | --- | --- |
| ***Drift rate (v)*** |  |  |  |  |  |
| Intercept, pre-stress | -0.002 (0.115) | [-0.230, 0.222] | 0.043 (0.102) | [-0.154, 0.244] | 0.385 |
| Intercept | 0.005 (0.108) | [-0.208, 0.217] | 0.101 (0.095) | [-0.082, 0.287] | 0.249 |
| Intercept, stress-induced change | 0.007 (0.064) | [-0.118, 0.136] | 0.057 (0.057) | [-0.054, 0.169] | 0.276 |
| Reward, pre-stress | **0.620 (0.050)** | **[0.521, 0.718]** | **0.661 (0.038)** | **[0.587, 0.738]** | 0.254 |
| Reward | **0.805 (0.049)** | **[0.710, 0.900]** | **0.759 (0.037)** | **[0.687, 0.834]** | 0.777 |
| Reward, stress-induced change | **0.185 (0.034)** | **[0.119, 0.254]** | **0.098 (0.028)** | **[0.041, 0.152]** | **0.978** |
| Aversion, pre-stress | **-0.623 (0.043)** | **[-0.709, -0.537]** | **-0.658 (0.037)** | **[-0.731, -0.586]** | 0.735 |
| Aversion | **-0.720 (0.044)** | **[-0.808, -0.634]** | **-0.672 (0.039)** | **[-0.748, -0.597]** | 0.205 |
| Aversion, stress-induced change | **-0.097 (0.038)** | **[-0.173, -0.022]** | -0.014 (0.033) | [-0.076, 0.055] | **0.043** |
| Positive domain, pre-stress | **1.076 (0.102)** | **[0.878, 1.277]** | **1.067 (0.098)** | **[0.874, 1.258]** | 0.530 |
| Positive domain | **1.346 (0.101)** | **[1.151, 1.546]** | **1.209 (0.090)** | **[1.033, 1.385]** | 0.845 |
| Positive domain, stress-induced change | **0.270 (0.075)** | **[0.119, 0.412]** | 0.142 (0.072) | [-0.000, 0.284] | 0.893 |
| Reward, positive domain, pre-stress | **-0.339 (0.060)** | **[-0.455, -0.221]** | **-0.400 (0.062)** | **[-0.521, -0.280]** | 0.763 |
| Reward, positive domain | **-0.565 (0.061)** | **[-0.685, -0.442]** | **-0.482 (0.058)** | **[-0.596, -0.367]** | 0.159 |
| Reward, positive domain, stress-induced change | **-0.226 (0.060)** | **[-0.337, -0.105]** | -0.082 (0.053) | [-0.184, 0.024] | **0.032** |
| Aversion, positive domain, pre-stress | **0.228 (0.064)** | **[0.103, 0.354]** | **0.277 (0.056)** | **[0.168, 0.390]** | 0.280 |
| Aversion, positive domain | **0.431 (0.065)** | **[0.304, 0.559]** | **0.392 (0.058)** | **[0.277, 0.506]** | 0.675 |
| Aversion, positive domain, stress-induced change | **0.202 (0.057)** | **[0.088, 0.316]** | **0.115 (0.045)** | **[0.014, 0.190]** | 0.899 |
| pgACC, pre-stress | 0.007 (0.032) | [-0.055, 0.068] | **-0.074 (0.027)** | **[-0.126, -0.021]** | **0.979** |
| pgACC | **-0.081 (0.027)** | **[-0.132, -0.028]** | -0.030 (0.025) | [-0.077, 0.020] | 0.082 |
| pgACC, stress-induced change | **-0.088 (0.040)** | **[-0.166, -0.011]** | 0.044 (0.037) | [-0.025, 0.113] | **0.003** |
| Reward × pgACC, pre-stress | -0.016 (0.027) | [-0.072, 0.037] | -0.038 (0.022) | [-0.081, 0.006] | 0.744 |
| Reward × pgACC | -0.002 (0.021) | [-0.048, 0.040] | 0.005 (0.019) | [-0.031, 0.046] | 0.432 |
| Reward × pgACC, stress-induced change | 0.014 (0.035) | [-0.055, 0.081] | 0.044 (0.029) | [-0.012, 0.101] | 0.263 |
| Aversion × pgACC, pre-stress | -0.049 (0.030) | [-0.106, 0.010] | 0.037 (0.024) | [-0.012, 0.085] | **0.005** |
| Aversion × pgACC | **0.055 (0.022)** | **[0.012, 0.100]** | **0.061 (0.023)** | **[0.016, 0.105]** | 0.412 |
| Aversion × pgACC, stress-induced change | **0.104 (0.036)** | **[0.035, 0.171]** | 0.024 (0.033) | [-0.040, 0.084] | **0.954** |
| pgACC, positive domain, pre-stress | -0.010 (0.050) | [-0.104, 0.090] | 0.079 (0.045) | [-0.012, 0.172] | 0.086 |
| pgACC, positive domain | **0.155 (0.043)** | **[0.068, 0.236]** | 0.048 (0.040) | [-0.031, 0.122] | **0.976** |
| pgACC, positive domain, stress-induced change | **0.165 (0.065)** | **[0.038, 0.290]** | -0.031 (0.060) | [-0.150, 0.090] | **0.987** |
| Reward × pgACC, positive domain, pre-stress | -0.011 (0.041) | [-0.090, 0.073] | 0.010 (0.038) | [-0.065, 0.084] | 0.349 |
| Reward × pgACC, positive domain | -0.014 (0.035) | [-0.078, 0.063] | -0.019 (0.028) | [-0.073, 0.038] | 0.516 |
| Reward × pgACC, positive domain, stress-induced change | -0.003 (0.052) | [-0.107, 0.104] | -0.030 (0.048) | [-0.125, 0.065] | 0.631 |
| Aversion × pgACC, positive domain, pre-stress | 0.021 (0.039) | [-0.057, 0.103] | -0.021 (0.035) | [-0.089, 0.045] | 0.800 |
| Aversion × pgACC, positive domain | -0.049 (0.031) | [-0.112, 0.011] | **-0.062 (0.027)** | **[-0.113, -0.007]** | 0.616 |
| Aversion × pgACC, positive domain, stress-induced change | -0.070 (0.048) | [-0.162, 0.027] | -0.042 (0.042) | [-0.119, 0.047] | 0.321 |
| ***Decision threshold (a)*** |  |  |  |  |  |
| Intercept, pre-stress | **1.890 (0.069)** | **[1.756, 2.027]** | **1.816 (0.054)** | **[1.712, 1.925]** | 0.803 |
| Intercept | **1.737 (0.067)** | **[1.611, 1.874]** | **1.684 (0.050)** | **[1.587, 1.784]** | 0.743 |
| Intercept, stress-induced change | **-0.153 (0.056)** | **[-0.262, -0.045]** | **-0.133 (0.045)** | **[-0.220, -0.046]** | 0.393 |
| STN, pre-stress | **0.051 (0.018)** | **[0.016, 0.086]** | **0.045 (0.014)** | **[0.018, 0.072]** | 0.593 |
| STN | **0.078 (0.015)** | **[0.049, 0.106]** | 0.022 (0.012) | [-0.001, 0.047] | **0.999** |
| STN, stress-induced change | 0.028 (0.022) | [-0.015, 0.070] | -0.024 (0.017) | [-0.056, 0.013] | **0.971** |
| ***Rate of threshold collapse (θ)*** |  |  |  |  |  |
| Intercept, pre-stress | **0.581 (0.025)** | **[0.531, 0.629]** | **0.584 (0.022)** | **[0.541, 0.626]** | 0.469 |
| Intercept | **0.556 (0.025)** | **[0.507, 0.604]** | **0.580 (0.020)** | **[0.541, 0.618]** | 0.216 |
| Intercept, stress-induced change | -0.025 (0.026) | [-0.076, 0.025] | -0.004 (0.022) | [-0.047, 0.040] | 0.255 |
| Conflict, pre-stress | **-0.025 (0.009)** | **[-0.042, -0.007]** | -0.004 (0.010) | [-0.022, 0.015] | 0.053 |
| Conflict | 0.008 (0.010) | [-0.011, 0.027] | -0.002 (0.009) | [-0.020, 0.017] | 0.761 |
| Conflict, stress-induced change | **0.033 (0.012)** | **[0.008, 0.056]** | 0.002 (0.012) | [-0.021, 0.025] | **0.962** |
| STN, pre-stress | **-0.024 (0.011)** | **[-0.046, -0.001]** | **-0.029 (0.009)** | **[-0.047, -0.011]** | 0.655 |
| STN | 0.012 (0.011) | [-0.009, 0.032] | **-0.039 (0.009)** | **[-0.056, -0.020]** | **1.000** |
| STN, stress-induced change | **0.035 (0.015)** | **[0.006, 0.065]** | -0.009 (0.012) | [-0.032, 0.015] | **0.993** |
| Conflict × STN, pre-stress | 0.015 (0.008) | [-0.002, 0.032] | 0.008 (0.007) | [-0.007, 0.022] | 0.745 |
| Conflict × STN | 0.001 (0.009) | [-0.016, 0.017] | -0.002 (0.007) | [-0.016, 0.012] | 0.599 |
| Conflict × STN, stress-induced change | -0.014 (0.011) | [-0.037, 0.008] | -0.010 (0.009) | [-0.027, 0.009] | 0.389 |
| ***Starting point bias (z)*** |  |  |  |  |  |
| Intercept, pre-stress | **0.545 (0.009)** | **[0.527, 0.563]** | **0.545 (0.008)** | **[0.530, 0.560]** | 0.487 |
| Intercept | **0.547 (0.010)** | **[0.527, 0.566]** | **0.543 (0.008)** | **[0.527, 0.559]** | 0.589 |
| Intercept, stress-induced change | 0.007 (0.010) | [-0.013, 0.028] | -0.009 (0.008) | [-0.025, 0.006] | 0.892 |
| Pavlovian incongruent, pre-stress | 0.004 (0.009) | [-0.014, 0.022] | 0.003 (0.009) | [-0.015, 0.021] | 0.540 |
| Pavlovian incongruent | **0.027 (0.009)** | **[0.011, 0.044]** | **0.020 (0.007)** | **[0.005, 0.034]** | 0.750 |
| Pavlovian incongruent, stress-induced change | **0.023 (0.010)** | **[0.004, 0.043]** | 0.017 (0.009) | [-0.001, 0.035] | 0.677 |
| ***Nondecision time (Tₑᵣ)*** |  |  |  |  |  |
| Intercept, pre-stress | **0.704 (0.022)** | **[0.660, 0.748]** | **0.682 (0.021)** | **[0.640, 0.725]** | 0.748 |
| Intercept | **0.654 (0.019)** | **[0.617, 0.692]** | **0.629 (0.016)** | **[0.597, 0.661]** | 0.853 |
| Intercept, stress-induced change | **-0.049 (0.018)** | **[-0.085, -0.014]** | **-0.054 (0.018)** | **[-0.089, -0.018]** | 0.572 |

Note. Estimates from the full neural model, including all higher-order interactions, retained as a robustness check. Values are posterior means with standard deviations (SD) and 95% highest density intervals (HDI). Bold within-group values indicate that the 95% HDI excludes zero. PP[rMDD > HC] is the posterior probability that a parameter is larger in rMDD than in HC, defined as the proportion of the posterior distribution of the group difference lying above zero; bold values indicate PP >= 0.95 or PP <= 0.05. Terms are grouped by model parameter. Unless labelled positive domain, estimates refer to the negative domain, which served as the reference. Terms labelled stress-induced change represent the difference between conditions (post minus pre-stress); terms labelled pre-stress are estimates for the pre-stress condition derived from the same posteriors.

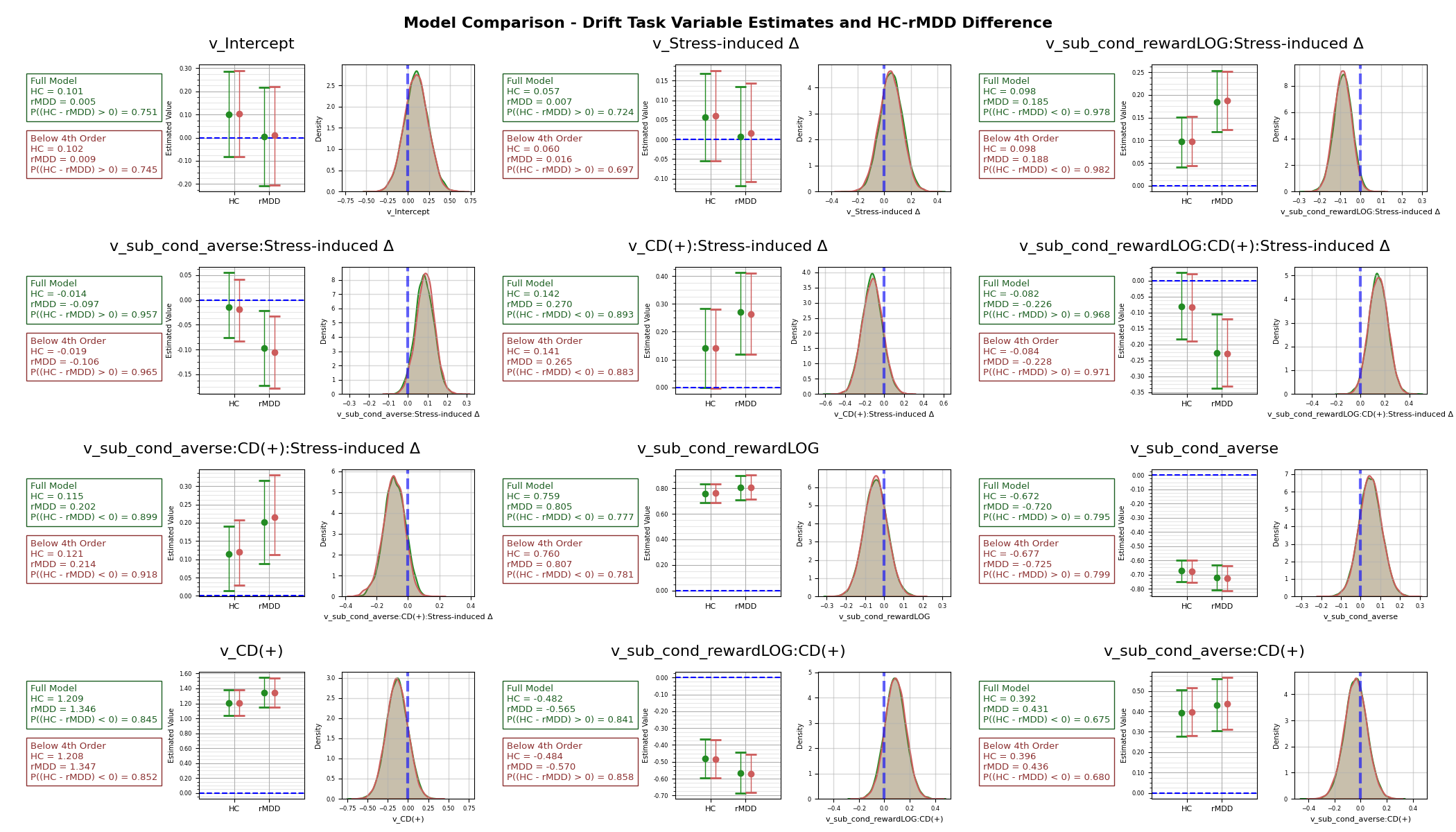

**Supplementary Fig. 12.** Comparison between the full and reduced neural models for drift parameters based on task variables. Plots on the left represent the estimates for each group for each model, while the distribution plots on the right show the directionality of the difference between HC and rMDD. The distribution plots are visualizations of the posterior distributions of the sampled difference between HC and rMDD for the indicated parameter.

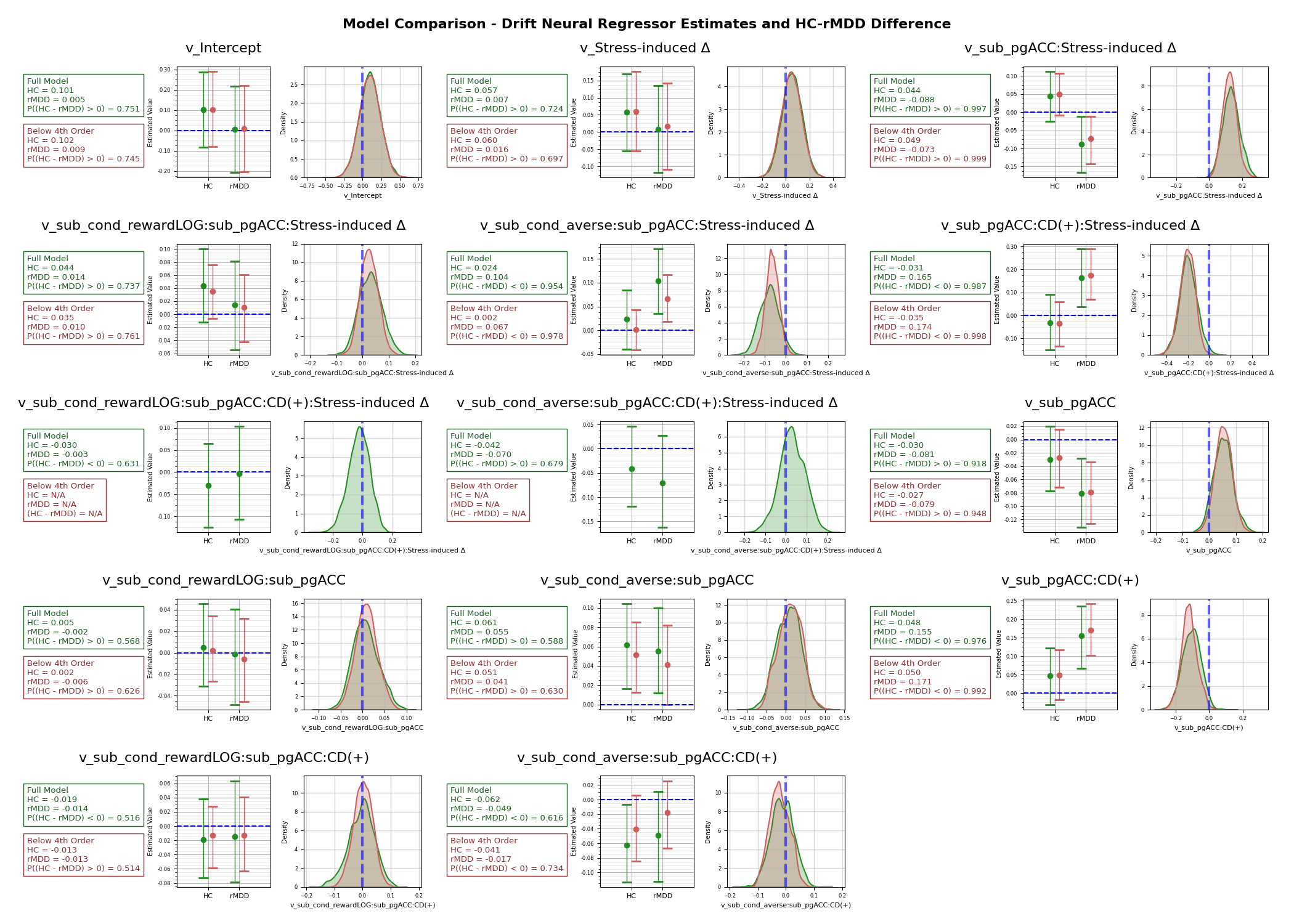

**Supplementary Fig. 13.** Comparison between the full and reduced neural models for drift parameters related to the neural regressor pgACC on drift rate. Plots on the left represent the estimates for each group for each model, while the distribution plots on the right show the directionality of the difference between HC and rMDD. The distribution plots are visualizations of the posterior distributions of the sampled difference between HC and rMDD for the indicated parameter. Panels showing only the full model correspond to fourth-order interaction terms, which by definition are not estimated in the reduced model.

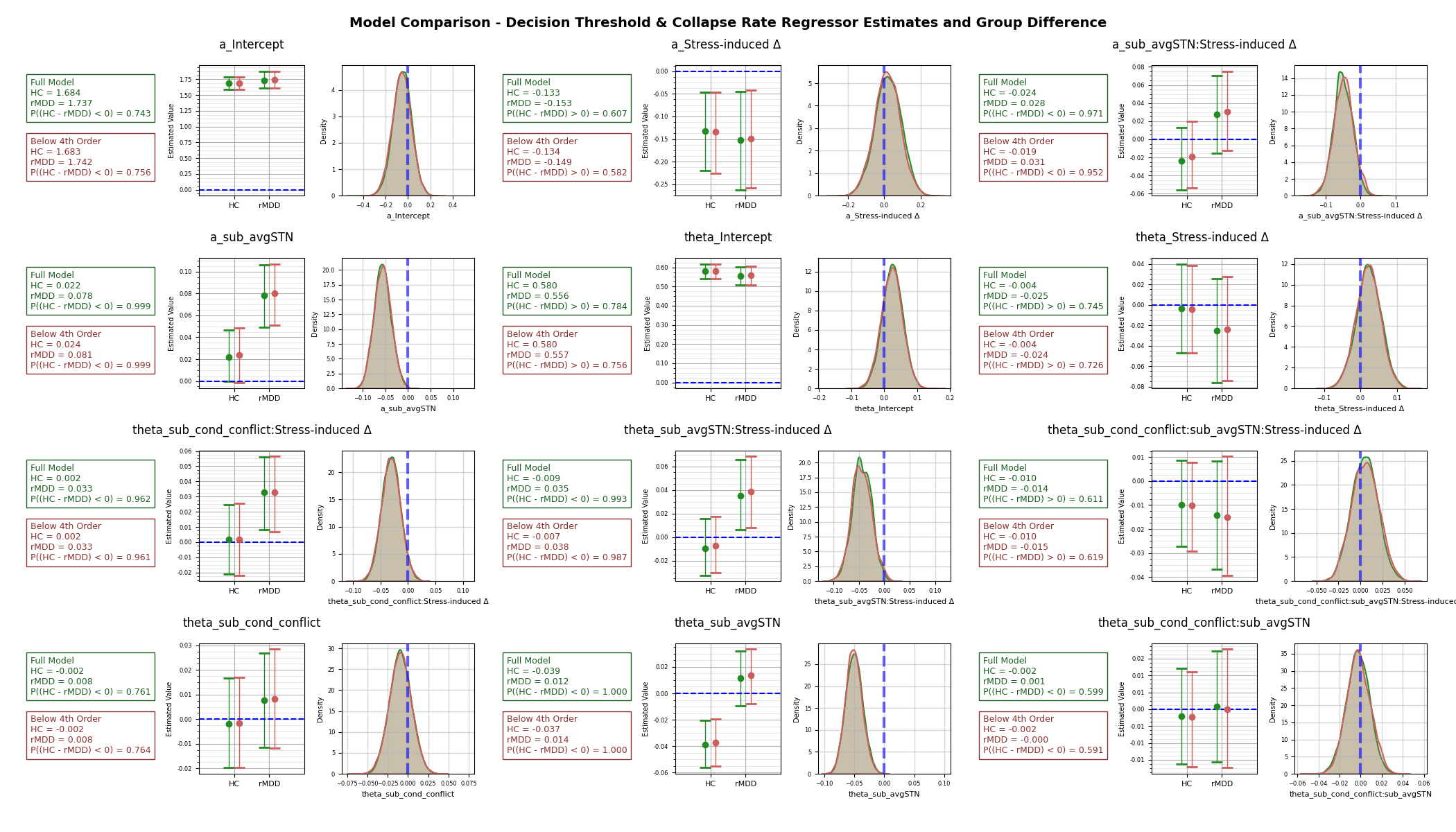

**Supplementary Fig. 14.** Comparison between the full and reduced neural models for regressors on decision threshold (a) and rate of threshold collapse (θ). Plots on the left represent the estimates for each group for each model, while the distribution plots on the right show the directionality of the difference between HC and rMDD. The distribution plots are visualizations of the posterior distributions of the sampled difference between HC and rMDD for the indicated parameter.

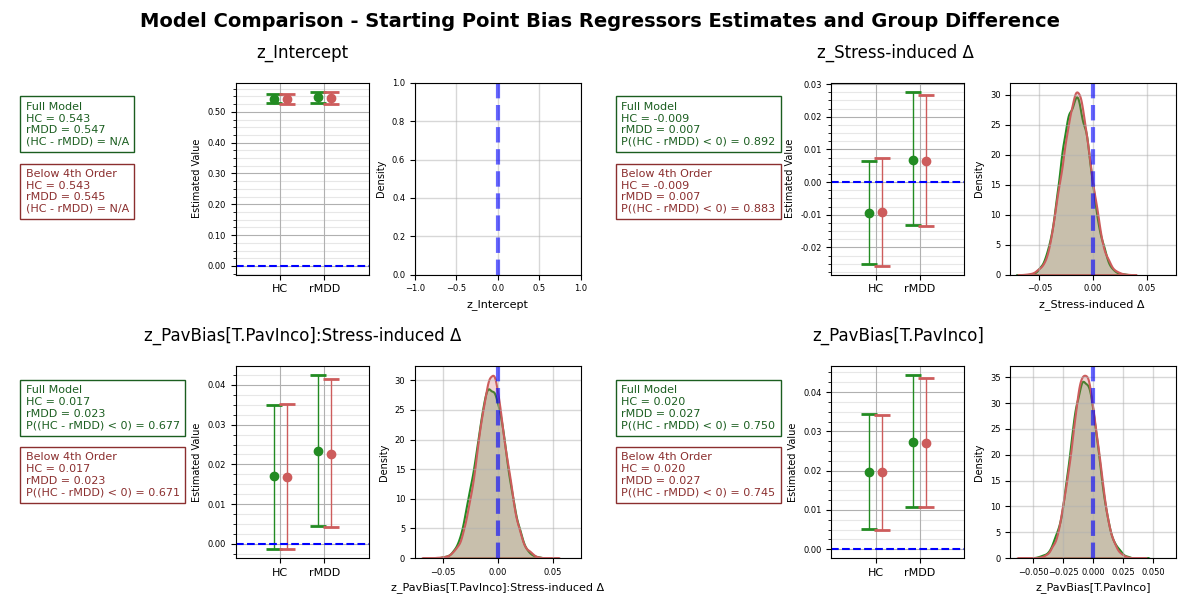

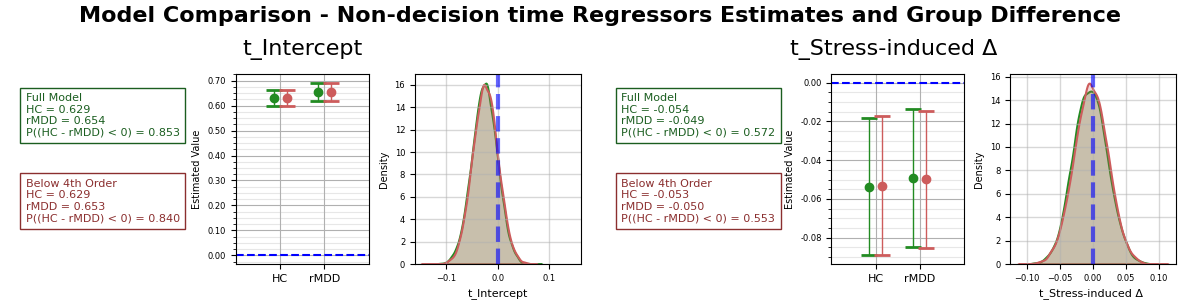

**Supplementary Fig. 15.** Comparison between the full and reduced neural models for regressors on starting point bias and non-decision time. Plots on the left represent the estimates for each group for each model, while the distribution plots on the right show the directionality of the difference between HC and rMDD. The distribution plots are visualizations of the posterior distributions of the sampled difference between HC and rMDD for the indicated parameter. No group-difference distribution is shown for the starting point intercept. Its group comparison is reported in Supplementary Tables 2 to 4.

**Supplementary Table 5. Generalized estimating equation models of HAMD across baseline, 6 and 12 months, net-aversive domain.**

| **Model term** | ***b*** | **SE** | **95% CI** | ***p*** |
| --- | --- | --- | --- | --- |
| ***Remitted MDD (142 observations, 50 participants)*** | | | | |
| Intercept | 1.148 | 0.315 | [0.530, 1.765] | <.001*** |
| Reward change | 0.094 | 0.249 | [−0.393, 0.582] | .704 |
| Time (months) | 0.084 | 0.043 | [−0.000, 0.169] | .051 |
| Reward change × Time | 0.108 | 0.054 | [0.002, 0.214] | .045* |
| ***Healthy controls (196 observations, 74 participants)*** | | | | |
| Intercept | 0.482 | 0.130 | [0.227, 0.737] | <.001*** |
| Reward change | 0.044 | 0.137 | [−0.225, 0.312] | .751 |
| Time (months) | 0.022 | 0.015 | [−0.009, 0.052] | .160 |
| Reward change × Time | −0.013 | 0.020 | [−0.052, 0.026] | .521 |
| ***Both groups (338 observations, 124 participants)*** | | | | |
| Intercept | 0.482 | 0.130 | [0.227, 0.737] | <.001*** |
| Reward change | 0.044 | 0.137 | [−0.225, 0.313] | .751 |
| Time (months) | 0.022 | 0.015 | [−0.008, 0.052] | .159 |
| Group (rMDD) | 0.666 | 0.341 | [−0.002, 1.334] | .051 |
| Reward change × Time | −0.013 | 0.020 | [−0.052, 0.026] | .522 |
| Reward change × Group | 0.051 | 0.284 | [−0.506, 0.608] | .857 |
| Time × Group | 0.062 | 0.046 | [−0.027, 0.152] | .173 |
| Reward change × Time × Group | 0.121 | 0.058 | [0.008, 0.234] | .036* |

Note. Generalized estimating equations with a Gaussian distribution, an exchangeable working correlation and robust sandwich standard errors, with participant as the clustering unit. Reward change is the stress-induced change in the influence of reward magnitude on drift rate for net-aversive offers, standardized across the whole sample. Time is in months from baseline, so coefficients are in HAMD points per month. In the both-groups model, healthy controls are the reference category. HAMD scores in healthy controls remained close to floor at every visit (means 0.47, 0.61 and 0.83 at baseline, 6 and 12 months; Table 2), so an absent association in that group may reflect the restricted range of symptoms. *p < .05, **p < .01, ***p < .001. HAMD = Hamilton Depression Rating Scale.

**Supplementary Table 6. Baseline-adjusted models of HAMD at 6 and 12 months.**

| **Effect** | ***b*** | **SE** | **95% CI** | ***F*** | ***p*** |
| --- | --- | --- | --- | --- | --- |
| ***6-month follow-up (n = 109)*** | | | | | |
| Intercept | 0.775 | 0.308 | [0.172, 1.379] | 6.34 | .012* |
| Reward change | 0.156 | 0.335 | [−0.502, 0.813] | 0.22 | .643 |
| Group (rMDD) | 1.095 | 0.506 | [0.104, 2.087] | 4.69 | .030* |
| Baseline HAMD | 0.298 | 0.164 | [−0.024, 0.620] | 3.30 | .069 |
| Reward change × Group | 0.430 | 0.497 | [−0.544, 1.404] | 0.75 | .387 |
| ***12-month follow-up (n = 105)*** | | | | | |
| Intercept | 0.843 | 0.447 | [−0.034, 1.719] | 3.55 | .060 |
| Reward change | −0.189 | 0.483 | [−1.137, 0.758] | 0.15 | .695 |
| Group (rMDD) | 1.073 | 0.729 | [−0.356, 2.502] | 2.17 | .141 |
| Baseline HAMD | 0.463 | 0.238 | [−0.005, 0.930] | 3.76 | .052 |
| Reward change × Group | 1.566 | 0.704 | [0.186, 2.947] | 4.95 | .026* |

Note. Ordinary least squares models of HAMD at each follow-up visit with baseline HAMD as a covariate, fitted separately at 6 and 12 months. Reward change and baseline HAMD are mean-centered, so the Group coefficient is the adjusted group difference at the sample mean. Healthy controls are the reference category. *p < .05, **p < .01, ***p < .001. HAMD = Hamilton Depression Rating Scale.

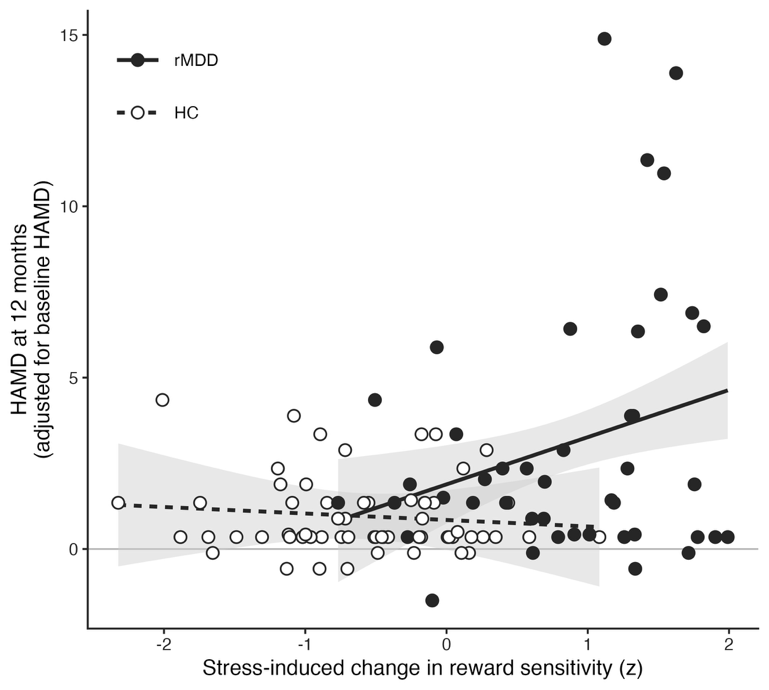

**Supplementary Fig. 16.** Association between the stress-induced change in reward sensitivity and HAMD at 12 months. HAMD at 12 months, adjusted for baseline HAMD, plotted against the stress-induced change in the influence of reward magnitude on drift rate for net-aversive offers, standardized across the whole sample, in 45 rMDD (filled circles, solid line) and 60 HC participants (open circles, dashed line). Lines show each group's slope from the baseline-adjusted model in Supplementary Table 6, drawn over the range of the predictor observed in that group, with shaded 95% confidence bands. Scores are adjusted to the sample-mean baseline HAMD, so participants with above-average baseline scores can fall below zero. The slopes differed between groups (b = 1.57, F(1,100) = 4.95, p = .026). HAMD = Hamilton Depression Rating Scale.

**Supplementary Table 7. Association between stress-induced reward sensitivity and the subsequent course of symptoms, by group and domain**

|  |  | **Within rMDD** | | **Within HC** | | **Predictor × time × group** | |
| --- | --- | --- | --- | --- | --- | --- | --- |
| **Measure** | **Domain** | ***b [95% CI]*** | ***p*** | ***b [95% CI]*** | ***p*** | ***b [95% CI]*** | ***p*** |
| HAMD | Net-aversive | 0.108 [0.002, 0.214] | 0.045* | −0.013 [−0.052, 0.026] | 0.521 | 0.121 [0.008, 0.234] | 0.036* |
|  | Net-rewarding | 0.080 [−0.009, 0.168] | 0.078 | −0.004 [−0.014, 0.005] | 0.374 | 0.084 [−0.005, 0.173] | 0.065 |
| BDI-II | Net-aversive | 0.155 [0.003, 0.307] | 0.046* | −0.020 [−0.110, 0.070] | 0.665 | 0.175 [−0.001, 0.352] | 0.052 |
|  | Net-rewarding | 0.092 [−0.024, 0.208] | 0.121 | −0.007 [−0.040, 0.025] | 0.648 | 0.099 [−0.021, 0.220] | 0.106 |
| QIDS-SR16 | Net-aversive | 0.076 [0.002, 0.149] | 0.043* | 0.015 [−0.013, 0.042] | 0.306 | 0.061 [−0.017, 0.140] | 0.125 |
|  | Net-rewarding | 0.059 [−0.001, 0.119] | 0.053 | −0.000 [−0.009, 0.009] | 0.955 | 0.059 [−0.001, 0.120] | 0.055 |
| SHAPS | Net-aversive | −0.090 [−0.366, 0.186] | 0.523 | 0.073 [−0.117, 0.262] | 0.451 | −0.147 [−0.482, 0.187] | 0.389 |
|  | Net-rewarding | −0.062 [−0.298, 0.173] | 0.602 | 0.010 [−0.034, 0.053] | 0.665 | −0.063 [−0.303, 0.177] | 0.609 |
| PSS-14 | Net-aversive | 0.054 [−0.210, 0.318] | 0.687 | 0.020 [−0.157, 0.196] | 0.827 | 0.049 [−0.269, 0.367] | 0.761 |
|  | Net-rewarding | 0.004 [−0.208, 0.216] | 0.970 | −0.025 [−0.076, 0.026] | 0.333 | 0.038 [−0.180, 0.255] | 0.734 |
| Q-LES-Q-SF | Net-aversive | −0.114 [−0.329, 0.101] | 0.298 | −0.073 [−0.202, 0.056] | 0.268 | −0.042 [−0.292, 0.208] | 0.742 |
|  | Net-rewarding | −0.094 [−0.269, 0.081] | 0.292 | 0.021 [−0.021, 0.062] | 0.325 | −0.116 [−0.296, 0.064] | 0.205 |

Note. Each coefficient represents the interaction between the stress-induced change in reward sensitivity and time, in points on the measure per month per SD of the predictor. The Within rMDD column reports that interaction fitted in the rMDD group alone (n = 50), the Within HC column the same model fitted in the control group (n = 74), and the Predictor × time × group column the three-way interaction in the full sample (n = 124), testing whether the association differs between groups. Both domains are expressed on a common scale, the pooled SD of the net-aversive parameter, so a unit means the same raw change in either domain. Models are generalized estimating equations with an exchangeable working correlation and robust standard errors. *p < .05, **p < .01, ***p < .001. HAMD = Hamilton Depression Rating Scale; BDI-II = Beck Depression Inventory-II; QIDS-SR16 = Quick Inventory of Depressive Symptomatology, Self-Report, 16 items; SHAPS = Snaith-Hamilton Pleasure Scale; PSS-14 = Perceived Stress Scale; Q-LES-Q-SF = Quality of Life Enjoyment and Satisfaction Questionnaire, Short Form.

**Supplementary References**

1.	Smeets, T. *et al.* Introducing the Maastricht Acute Stress Test (MAST): a quick and non-invasive approach to elicit robust autonomic and glucocorticoid stress responses. *Psychoneuroendocrinology* **37**, 1998–2008 (2012).

2.	Pizzagalli, D. A., Jahn, A. L. & O’Shea, J. P. Toward an objective characterization of an anhedonic phenotype: A signal-detection approach. *Biological Psychiatry* **57**, 319–327 (2005).

3.	Treadway, M. T., Buckholtz, J. W., Schwartzman, A. N., Lambert, W. E. & Zald, D. H. Worth the ‘EEfRT’? The Effort Expenditure for Rewards Task as an Objective Measure of Motivation and Anhedonia. *PLoS ONE* **4**, e6598 (2009).

4.	Watson, D., Clark, L. A. & Tellegen, A. Development and validation of brief measures of positive and negative affect: The PANAS scales. *Journal of Personality and Social Psychology* **54**, 1063–1070 (1988).

5.	Mendes, W. B., Blascovich, J., Major, B. & Seery, M. Challenge and threat responses during downward and upward social comparisons. *Euro J Social Psych* **31**, 477–497 (2001).

6.	Ironside, M. *et al.* Association of Lower Rostral Anterior Cingulate GABA+ and Dysregulated Cortisol Stress Response With Altered Functional Connectivity in Young Adults With Lifetime Depression: A Multimodal Imaging Investigation of Trait and State Effects. *Am J Psychiatry* appiajp20230382 (2024) doi:10.1176/appi.ajp.20230382.

7.	Whitton, A. E. *et al.* Acute stress impairs frontocingulate activation during error monitoring in remitted depression. *Psychoneuroendocrinology* **75**, 164–172 (2017).

8.	Admon, R. *et al.* Distinct Trajectories of Cortisol Response to Prolonged Acute Stress Are Linked to Affective Responses and Hippocampal Gray Matter Volume in Healthy Females. *J. Neurosci.* **37**, 7994–8002 (2017).

9.	Ging-Jehli, N. R. *et al.* Cognitive Signatures of Depressive and Anhedonic Symptoms and Affective States Using Computational Modeling and Neurocognitive Testing. *Biological Psychiatry: Cognitive Neuroscience and Neuroimaging* **9**, 726–736 (2024).

10.	Ironside, M. *et al.* Approach-Avoidance Conflict in Major Depressive Disorder: Congruent Neural Findings in Humans and Nonhuman Primates. *Biological Psychiatry* **87**, 399–408 (2020).

11.	Esteban, O. *et al.* fMRIPrep: a robust preprocessing pipeline for functional MRI. *Nat Methods* **16**, 111–116 (2019).

12.	Markiewicz, C. J. *et al.* fMRIPrep: a robust preprocessing pipeline for functional MRI. Zenodo https://doi.org/10.5281/zenodo.12774221 (2024).

13.	Gorgolewski, K. *et al.* Nipype: A Flexible, Lightweight and Extensible Neuroimaging Data Processing Framework in Python. *Front. Neuroinform.* **5**, (2011).

14.	Andersson, J. L. R., Skare, S. & Ashburner, J. How to correct susceptibility distortions in spin-echo echo-planar images: application to diffusion tensor imaging. *NeuroImage* **20**, 870–888 (2003).

15.	Tustison, N. J. *et al.* N4ITK: Improved N3 Bias Correction. *IEEE Transactions on Medical Imaging* **29**, 1310–1320 (2010).

16.	Avants, B. B., Epstein, C. L., Grossman, M. & Gee, J. C. Symmetric diffeomorphic image registration with cross-correlation: Evaluating automated labeling of elderly and neurodegenerative brain. *Medical Image Analysis* **12**, 26–41 (2008).

17.	Zhang, Y., Brady, M. & Smith, S. Segmentation of brain MR images through a hidden Markov random field model and the expectation-maximization algorithm. *IEEE Transactions on Medical Imaging* **20**, 45–57 (2001).

18.	Dale, A. M., Fischl, B. & Sereno, M. I. Cortical Surface-Based Analysis: I. Segmentation and Surface Reconstruction. *NeuroImage* **9**, 179–194 (1999).

19.	Klein, A. *et al.* Mindboggling morphometry of human brains. *PLOS Computational Biology* **13**, e1005350 (2017).

20.	Ciric, R. *et al.* TemplateFlow: FAIR-sharing of multi-scale, multi-species brain models. *Nat Methods* **19**, 1568–1571 (2022).

21.	Evans, A. C., Janke, A. L., Collins, D. L. & Baillet, S. Brain templates and atlases. *NeuroImage* **62**, 911–922 (2012).

22.	Fonov, V., Evans, A., McKinstry, R., Almli, C. & Collins, D. Unbiased nonlinear average age-appropriate brain templates from birth to adulthood. *NeuroImage* **47**, S102 (2009).

23.	Jenkinson, M., Bannister, P., Brady, M. & Smith, S. Improved Optimization for the Robust and Accurate Linear Registration and Motion Correction of Brain Images. *NeuroImage* **17**, 825–841 (2002).

24.	Cox, R. W. & Hyde, J. S. Software tools for analysis and visualization of fMRI data. *NMR in Biomedicine* **10**, 171–178 (1997).

25.	Greve, D. N. & Fischl, B. Accurate and robust brain image alignment using boundary-based registration. *NeuroImage* **48**, 63–72 (2009).

26.	Power, J. D. *et al.* Methods to detect, characterize, and remove motion artifact in resting state fMRI. *NeuroImage* **84**, 320–341 (2014).

27.	Behzadi, Y., Restom, K., Liau, J. & Liu, T. T. A component based noise correction method (CompCor) for BOLD and perfusion based fMRI. *NeuroImage* **37**, 90–101 (2007).

28.	Satterthwaite, T. D. *et al.* An improved framework for confound regression and filtering for control of motion artifact in the preprocessing of resting-state functional connectivity data. *NeuroImage* **64**, 240–256 (2013).

29.	Patriat, R., Reynolds, R. C. & Birn, R. M. An improved model of motion-related signal changes in fMRI. *NeuroImage* **144**, 74–82 (2017).

30.	Lanczos, C. Evaluation of Noisy Data. *Journal of the Society for Industrial and Applied Mathematics Series B Numerical Analysis* **1**, 76–85 (1964).

31.	Kuhn, M. *et al.* Psychobiological Stress Response Profiles in Current and Remitted Depression: A Person-Centered, Multisystem Approach. *Biological Psychiatry Global Open Science* **5**, 100400 (2025).

32.	Berghorst, L. H., Bogdan, R., Frank, M. J. & Pizzagalli, D. A. Acute stress selectively reduces reward sensitivity. *Front. Hum. Neurosci.* **7**, (2013).

33.	Buuren, S. van & Groothuis-Oudshoorn, K. mice: Multivariate Imputation by Chained Equations in R. *Journal of Statistical Software* **45**, 1–67 (2011).

34.	Pedersen, M. L. *et al.* Computational phenotyping of brain-behavior dynamics underlying approach-avoidance conflict in major depressive disorder. *PLoS Comput Biol* **17**, e1008955 (2021).
